# Epigenetic disruption of the RARγ complex impairs its function to bookmark AR enhancer interactions required for enzalutamide sensitivity in prostate cancer

**DOI:** 10.1101/2023.12.15.571947

**Authors:** Sajad A Wani, Shahid Hussain, Jaimie S Gray, Debasis Nayak, Hancong Tang, Lillian M Perez, Mark D Long, Manjunath Siddappa, Christopher J McCabe, Lara E Sucheston-Campbell, Michael R Freeman, Moray J Campbell

## Abstract

The current study in prostate cancer (PCa) focused on the genomic mechanisms at the cross-roads of pro-differentiation signals and the emergence of lineage plasticity. We explored an understudied cistromic mechanism involving RARγ’s ability to govern AR cistrome-transcriptome relationships, including those associated with more aggressive PCa features. The RARγ complex in PCa cell models was enriched for canonical cofactors, as well as proteins involved in RNA processing and bookmarking. Identifying the repertoire of miR-96 bound and regulated gene targets, including those recognition elements marked by m6A, revealed their significant enrichment in the RARγ complex. RARγ significantly enhanced the AR cistrome, particularly in active enhancers and super-enhancers, and overlapped with the binding of bookmarking factors. Furthermore, RARγ expression led to nucleosome-free chromatin enriched with H3K27ac, and significantly enhanced the AR cistrome in G_2_/M cells. RARγ functions also antagonized the transcriptional actions of the lineage master regulator ONECUT2. Similarly, gene programs regulated by either miR-96 or antagonized by RARγ were enriched in alternative lineages and more aggressive PCa phenotypes. Together these findings reveal an under-investigated role for RARγ, modulated by miR-96, to bookmark enhancer sites during mitosis. These sites are required by the AR to promote transcriptional competence, and emphasize luminal differentiation, while antagonizing ONECUT2.

## INTRODUCTION

Patients with advanced prostate cancer (PCa) receive androgen receptor (NR3C4/AR) signaling inhibitors (ARSI), such as such Enzalutamide (Enza), which are initially successful but recurrent PCa frequently emerges (reviewed in(1)). Acquisition of therapy resistance is frequently driven by altered AR functions, including enhancer switching to novel gene regulatory regions(2) that favor growth promotion instead of differentiation(3–7) and the acquisition of alternative lineages, such as neuroendocrine (NE) PCa.

Identifying the mechanisms that allow this shift in cell fate may reveal novel routes to restore luminal phenotypes, perhaps by targeting stem cell-like components, and make tumors more amenable to therapy(8). Arguably, this is the goal of Bipolar Androgen Therapy for PCa, in which the non-malignant features of AR signaling are sustained to support luminal differentiation(9). Supportively, recent studies suggest that luminal-type features can be restored by targeting the AR-interacting pioneer factor FOXA2(10) and other coregulators that govern Enza sensitivity(11).

The AR interacts with multiple other nuclear receptors (NRs) and coregulators. More widely NRs form a genomic network to sense various inputs to regulate cell fate decisions and metabolism (reviewed in(12)). Type I NRs, such as the AR, bind ligand with high affinity, which leads to nuclear translocation and target gene regulation. By contrast, Type II NRs such as retinoic acid receptor gamma (NR1B3/RARG, encodes RARγ) are nuclear resident, independent of ligand, with ligand exposure leading to genomic redistribution and modulation of target genes. The AR in the normal prostate gland promotes luminal phenotypes, survival and can induce proliferation (13), whereas RARγ promotes prostate growth restraint (14).

One mechanism for crosstalk arising between NRs is through shared genomic binding leading to combinatorial gene regulation. For example, NR-genomic interactions can converge on complex genomic sites such as high occupancy target (HOT) regions (15). In pluripotent cells so-called bookmarking functions have been revealed where mitotic complexes sustain nucleosome free positioning at genomic sites required for post-mitotic immediate transcriptional reactivation (16–19). There is some evidence to suggest that nuclear-resident Type II NRs can undertake bookmarking on mitotic chromatin (16,20–22). To date, NR-mediated bookmark mechanisms have neither been investigated in PCa nor cancer generally and could add further to understanding of crosstalk mechanisms between NRs.

There are well-studied examples of how the genomic interactions of the AR and other NRs are altered. Epigenomic mechanisms as a result of changes in expression of AR-interacting coregulators, such as the coactivator CBP/P300(23) and the corepressors NCOR1 and NCOR2(11,24,25) re-direct the AR to favor proliferative gene expression programs. Similarly, there are altered expression of microRNAs (miRNAs) target the pioneer factor FOXA1(26) and other lineage identity drivers. Across the PCa genome there are many infrequently mutated chromatin remodelers, the so-called “long-tail” mutant epigenetic drivers(27), which have the capacity to shape the wider transcriptional landscape. Intriguingly, it appears to be a relatively small group of transcription factors that are impacted by these widespread genomic and epigenomic events, perhaps fewer than 100(28). These include ONECUT2, a master regulator of metastasis and drug resistance in PCa(29).

We previously showed that RARγ was significantly down-regulated in 60% of localized PCa tumors compared to non-malignant samples(30). In non-malignant prostate cells miR-96-5p (miR-96) targets *RARG* and a known coactivator *TACC1*, and RARγ expression impacted DHT- signaling. Finally, RARγ binding sites were enriched for the ONECUT2 motif (31).

In the current study we sought to define the impact of the RARγ complex in the context of advanced PCa through three approaches (**Figure 1; Graphical Abstract**). Firstly, we undertook proteomic approaches to define the RARγ complex across PCa models. Given there are a wide number of identified miR-96 targets (32–37), we sought to define the prominence of the RARγ complex as a miR-96 target, and to test how the epi-modification N^6^-methyladenosine (m6A)(38,39) shaped the miR-96 targetome. Secondly, we examined how RARγ and TACC1 shaped the AR cistrome, and tested the possibility that RARγ could function to bookmark the AR cistrome and dictate the AR-dependent transcriptional response to DHT and Enza. Thirdly, we tested if the RARγ complex could antagonize the master regulator ONECUT2 to shape Enza responsiveness and limit transcriptional programs associated with advanced PCa.

**Figure 1:**
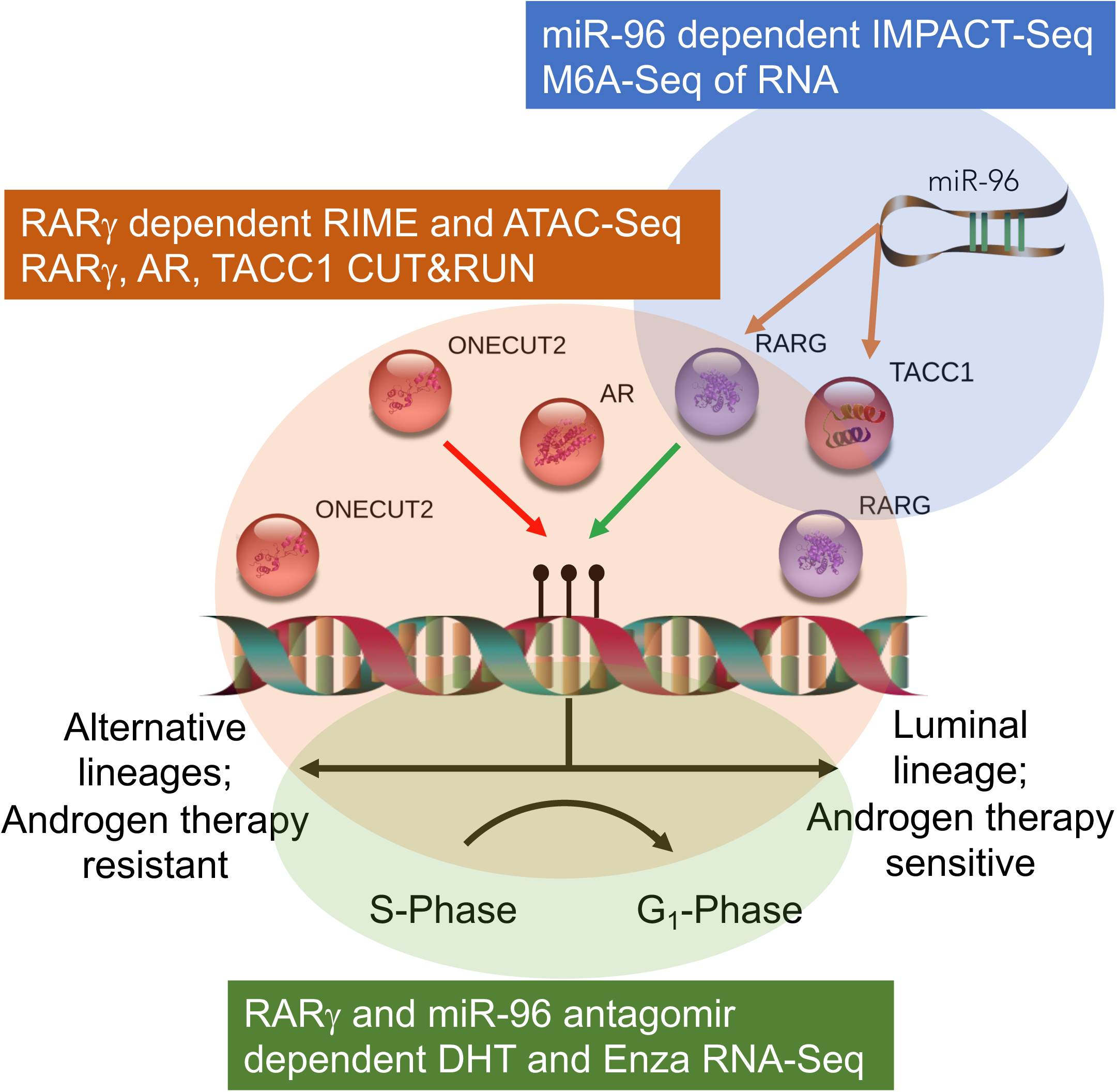
Graphical abstract of workflow and approaches.

Collectively our findings support a novel model in which the RARγ complex functions to bookmark the AR to limit aggressive phenotypes. In PCa progression RARγ is targeted by miR-96 to limit this capacity and to increase the interactions of AR with ONECUT2, as well as the regulation of transcriptional programs that associate with aggressive PCa phenotypes and alternative lineages.

## MATERIALS AND METHODS

### Reagents

DHT(Sigma), CD437 (Sigma), Enza (Sigma) mir-96 mimic and antagomir (Ambion, nocodazole (CST). For Western immunoblots the following antibodies were used; Anti-RARγ (D3A4) rabbit mAb #8965 Cell Signaling Technology (CST); Anti-Myc Tag (9B11); Anti-TACC1 mouse mAb #2276 (CST); Anti-AR-V7 rabbit mAb #ab198394 (Abcam); Anti-AR (D6F11) rabbit mAb #5153 (CST); Anti-PCNA rabbit polyclonal antibody #ab15497 (Abcam). For rapid immunoprecipitation mass spectrometry of endogenous protein (RIME) assay RARγ (D3A4) #8965 CST antibody was used. For CUT&RUN the following antibodies were used; Anti-Acetyl-Histone H3K27 (D5E4) XP® rabbit mAb #8173 (CST); anti-GFP antibody for RARγ, as previously(31) (ChIP Grade-ab290, Abcam); anti-AR (D6F11) XP® rabbit mAb #5153 (CST), anti-IgG #2729S (CST), rabbit anti-H3S10ph #39254 (Active Motif).

### Biological Resources

Prostate cancer cell lines LNCaP and 22Rv1 were obtained from the ATCC and maintained in RPMI-1640 media (Gibco) supplemented with 10% FBS and penicillin/ streptomycin. Non-malignant HPr1AR cells were maintained in keratinocyte serum free medium (SFM; Gibco) supplemented with penicillin/streptomycin/gentamicin(40). Cells were screened for mycoplasma every 2 months (Mycoplasma PCR Detection Kit (Sigma Aldrich) and authenticated regularly by short tandem repeat profiles at The Ohio State University Comprehensive Cancer Center Genomics Shared Resource.

### Lentiviral constructs and transduction

pLenti-CMV-RARγ-C vector (GFP) and pLenti-CMV-TACC1-C vector (MYC) and pLenti-III-Blank vector were custom synthesized (Applied Biological Materials). Lentiviral MOI were as per manufacturer’s protocol(41) and transfected into cells followed by positive selection with puromycin (1 μg/mL (Sigma)). TACC1 and RARγ expression was confirmed by qPCR and Western immunoblot.

These 22Rv1 isogenic cell variants are specifically 22Rv1 cells stably transfected with an empty vector (22Rv1-mock) or a GFP-tagged RARγ (22Rv1-RARγ). 22Rv1-RARγ-TACC1 cells were generated by transient transfection of TACC1 into 22Rv1-RARγ cells; we demonstrated TACC1 protein expression was sustained for ∼72h.

### Cell viability

Cells were plated in 96-well, white-walled plates as previously(31). At the indicated time points cellular ATP was measured using CellTiter-Glo^®^ (Promega) and luminescence detected with Synergy^TM^ 2 multi-mode microplate reader (BioTek® Instruments). Each experiment was performed in at least triplicate wells in triplicate experiments.

### Apoptosis Assay

Cells were treated with agent alone or with miR-96 inhibitor or scramble control, treated with Annexin-V-FLUOS (Roche Diagnostics) and analyzed by flow cytometer (Becton Dickinson).

### Cell cycle measurements

Cells were treated as for apoptosis, harvested, stained with PI and incubated (45 min in the dark) and cell cycle profiles analyzed on a flow cytometer (Becton Dickinson).

### Western Immunoblotting

Protein (30-60μg) were resolved via SDS polyacrylamide gel electrophoresis (SDS-PAGE) using precast 4-20% gradient polyacrylamide gels (Bio-Rad). Blocked membranes were probed with primary antibody for 3 hours at room temperature and subsequently for 1 hour species-appropriate secondary antibody at room temperature using enhanced chemiluminescence substrate (Pierce). Signal quantification was performed using the Protein Simple Fluorochem M Imager.

### RT-qPCR

Total RNA was isolated (TRIzol® reagent (Thermo Fisher Scientific)), complementary DNA (cDNA) prepared (iScriptTM cDNA Synthesis Kit (Bio-Rad)), and relative gene expression quantified via Applied Biosystems 7300 Real-Time PCR System (Applied Biosystems), for both TaqMan and SYBR Green (Thermo Fisher Scientific) applications. All SYBR Green primers were tested for specificity by melting curve analysis. Fold changes determined using the 2-ΔΔCt method. Gene expression master mix (Applied Biosystems) and the TaqMan assays RARγ (Hs01559230_m1), TACC1 (Hs00180691_m1), TWF1 (Hs00702289_s1), HMGA2 (Hs04397751_m1), MYC (Hs04397751_m1) were used to measure expression levels. 18s (Hs99999901_s1) and GAPDH (Hs02786624_g1) were used for normalization.

### Transfection of miRNA mimic and antagomir

Transient transfection of miRNA mimic (50nM) or inhibitor (100nM) (Ambion) was undertaken using Lipofectamine® 2000 in presence of Opti-MEM Reduced Serum Medium (Thermo fisher Scientific). Concentrations of miRNA mimic/inhibitor and transfection reagents were optimized using BLOCK-iTTM Alexa Fluor® Red Fluorescent Control (Ambion) as well as by mirVana™ miRNA mimic miR-1 positive control and mirVana™ miRNA Inhibitor, let-7c positive control.

### Identification of MRE by pull-down and alignment of captive transcripts—sequencing (IMPACT-Seq)

IMPACT-Seq(42) was undertaken on 5×10^6^ cells transfected with 3’ biotinylated miRIDIAN miR-96 mimic and scramble (50 nM, 24h and 48h) (Dharmacon, Inc.) using Lipofectamine 2000. RNA was then extracted from washed beads using Trizol LS and treated with T4 Polynucleotide Kinase (New England Biolabs) to obtain 5’ phosphate ends for subsequent ligations and passed through NucAway columns (Ambion) to remove RNAs <20 nt in length. Small RNA libraries were generated using NEBNext Multiplex Small RNA Library Prep (New England Biolabs) following the manufacturer’s recommendations for 100ng input material. PCR-amplified small RNA libraries were size selected to 130-200 bp fragments. Libraries characterized on an Agilent Tape Station on a DNA1000 High-Sensitivity Screen Tape and quantified by Qubit were pooled and sequenced at 75 bp read length on the MiSeq platform.

### m6A Sequencing

m6A seq was performed on 2×10^7^ cells using EpiMark N6-methyladenosine enrichment kit (New England Biolabs). The libraries were prepared from purified DNA, using the NEB Ultra II Library Prep Kit (New England Biolabs) per manufacturer’s instructions and the libraries were sequenced at 50 bp read length and were sequenced on an NovaSeq SP with a target of ∼40M PE reads per sample.

### Rapid immunoprecipitation mass spectrometry of endogenous protein (RIME)

RIME(43) was undertaken in presence or absence of CD437 (400nM, 6h). 2×10^7^ cells were crosslinked with 1% formaldehyde solution, quenched with glycine (0.1M, pH 7.5) and harvested in cold PBS. Nuclei were separated and subjected to sonication for genomic DNA fragmentation. Nuclear proteins were separated using 10% triton-X with high-speed centrifugation. RARγ or IgG antibody conjugated beads were incubated with nuclear lysates overnight and washed ten times with RIPA buffer then Ambic solution. Capillary-liquid chromatography-nanospray tandem mass spectrometry (LC-MS/MS) was acquired over a 2-hour separation using a data-dependent MS/MS approach at a resolving power to achieve high mass accuracy MS determination. Label free shotgun proteomics was used to eliminate nonspecific interactors(44).

### Assay for Transposase-Accessible Chromatin using sequencing (ATAC-Seq)

Omni-ATAC-seq(45) was undertaken in 5×10^4^ cells treated for 6h with DHT (10nM) or CD437 (400 nM). Libraries were PCR-amplified using the NEBNext Hi-Fidelity PCR Master Mix and Integrated DNA Technologies (IDT) 8 bp unique dual indexing (UDI) adapters. The PCR cycle number were optimized as indicated by qPCR fluorescence curves. Primer dimers and large >1,000 bp fragments were removed using AMPure XP beads (Beckman Coulter). The quality of libraries was checked on Agilent High Sensitivity DNA Bioanalysis and quantified by qubit. Libraries were sequenced on NovaSeq6000 S1 PE150bp (v1.5) with ∼60M reads per sample.

### Cleavage Under Targets and Release Using Nuclease (CUT&RUN)

CUT&RUN(46) was undertaken with antibodies to GFP (for RARγ), AR, H3K27ac, H3S10P and IgG. Briefly, 0.5×10^6^ cells were treated for DHT (10nM, 6h) or CD437 (400 nM, 6h). For TACC1, the pLenti-CMV-TACC1-C term RFP vector was transiently overexpressed for 48h followed by similar drug treatments and sustained protein expression confirmed. Harvested cells were washed with pre-activated Concanavalin A-coated beads mixed with antibody overnight at 4°C. CUTANA pAG-MNase (EpiCypher) was incubated with sample for 10 minutes at room temperature. Fragmented DNA samples was purified using the Monarch DNA Clean up Kit (New England Biolabs). The libraries were prepared from 5-10ng purified CUT&RUN-enriched DNA, using the NEB Ultra II Library Prep Kit (New England Biolabs) per manufacturer’s instructions and sequenced using Hiseq 4000 PE 150bp. Transcription factor modifications for sub-nucleosomal size (<120bp) DNA fragments were undertaken as per protocol.

### RNA-Seq

RNA Seq was undertaken in three experimental settings. 1. In the presence of miR-96 mimic/scramble (50nM) by transient transfection in HPr1-AR, LNCaP and 22Rv1 cells and RNA harvested at 48h. 2. In 22Rv1-RARγ-TACC1, 22Rv1-RARγ and 22Rv1-Mock cells treated with DHT (10nM) or CD437 (400nM) for 24h. 3. 22Rv1 cells were treated with the combination of ENZA and miR-96 inhibitor as above. Total RNA was extracted E.Z.N.A.® Total RNA Kit I, Omega Bio-tek®) and subjected to DNase treatment and purification by the RNA Clean & Concentrator kit (Zymo Research) to yield high-quality RNA for library preparation. RNA-seq libraries were prepared using Illumina’s TruSeq Stranded protocol. ∼70 million paired-end 150 bp sequence reads were generated for each library on the HiSeq 4000 platform.

### Label free quantitative proteomics (LFQ)

MiR-96 mimic/scramble (50nM) was transiently transfected into HPr1-AR, LNCaP and 22Rv1 cells for 48h. Protein concentration was measured and 70μg of protein was used for S-Trap Digestion. Capillary-LC/MS/MS was performed as for RIME

#### Data analyses and integration

All sequencing was performed at the Nationwide Children’s Hospital Institute for Genomic Medicine and FASTQC files were QC processed, adapters trimmed and aligned to Hg38 (Rsubread)(47).

### IMPACT Seq and m6A Seq

Differentially enriched miR-96 binding sites and m6A were identified with `csaw`(48) and annotated to genes (`ChIPpeakAnno`)(49), and only those retained that directly overlapped with a transcript. Shared binding was determined with a minimum of 1bp in common between peaks.

### RIME and LFQ

The mean spectral count results were generated by two protein inference engines (`Epifany`(50) and `FIDO`(51)) and processed in an `edgeR` workflow to identify differentially enriched proteins, visualized with volcano plots and network enrichment analyses (e.g. `stringdb`). Identified proteins were annotated enriched proteins by functional class (coactivator (CoA), corepressor (CoR), Mixed function coregulator (Mixed) or transcription factor (TF))(52). Hypergeometric tests were used to test enrichment for CoA, CoR, Mixed, TF, “long-tail” mutant epigenetic drivers(27), or miR-96 bound and regulated target genes, with or without m6A modification, in each RARγ complex.

### CUT&RUN

Cistromes were analyzed with `csaw` with appropriate window sizes for either TF (AR and RARγ) or larger ones for H3K27ac. Significantly differentially enriched regions (FDR < .1) were overlapped with ChromHMM(53) derived underlying epigenetic states in LNCaP (ref), and also mined for TF motif analyses `(MotifDb)`. `GIGGLE` was utilized to query the complete human transcription factor ChIP-seq dataset collection (10,361 and 10,031 datasets across 1,111 transcription factors and 75 histone marks, respectively) in Cistrome DB(54). Prostate specific filtering limited analysis to 681 datasets across 74 TFs and 238 datasets across 19 HMs. For each query dataset, the overlap with each experimental cistrome was determined. H3K27ac cistromes were analyzed for super-enhancers by combining with BRD4 cistrome from 22Rv1(55,56) using the `ROSE` algorithm(57).

### ATAC-Seq

ATAC-Seq data were separated into nucleosome free (NF), mono-, di- and tri-nucleosome compartments (ATACSeqQC)(58) and analyzed with csaw.

### RNA-Seq

Transcript abundance estimates were normalized and differentially expressed genes (DEGs) identified using a standard `edgeR` pipeline, and gene set enrichment analysis (`GSEA`) performed. For transcript-aware analyses, the FASTQ files were aligned with `salmon`(59) and differentially enriched transcripts were identified using `DRIMSeq`(60) in a similar workflow to `edgeR`.

### Cistrome-Transcriptome analyses

To test the significance of the ATAC-Seq or CUT&RUN cistromes to transcriptomes we applied a modification of the BETA method(61). Specifically, within each cell type we summed significance of the peaks within 100 kb of each annotated DEG multiplied by the absolute fold change for the same DEG and weighted by the peak distribution (proximal versus distal), or unweighted. We defined this score as the weighted cistrome-transcriptome (wt-C-T) and tested the difference between controls and modified cell backgrounds using a Wilcox test.

## RESULTS

### The RARγ complex is enriched for bookmarking and RNA splicing factors

Given RARγ levels are very low in 22Rv1 cells, we created an isogenic series of 22Rv1-mock, 22Rv1-RARγ and 22Rv1-RARγ-TACC1 sublines and treated the cells with the RARγ-selective ligand CD437. Phenotypically, 22Rv1-RARγ cells grew more slowly with longer G_1_ transit times than 22Rv1-Mock cells (data not shown).

Preliminary studies suggested RARγ and TACC1 interact (**Supplementary Figure 1A**), reflecting previous findings of others(62). To define the RARγ complex we undertook RIME. RARγ enrichment compared to IgG controls was broadly comparable in LNCaP and the 22Rv1 isogenic cells. In each case several hundred RARγ-interacting proteins were identified, including several known RARγ interacting proteins such as the heterodimer partners RXRβ and RXRα, the corepressor NCOR2, and other novel coregulators such as SSRP1, which is a member of the FACT chromatin remodeling complex(63) (**Supplementary Figure 1B**).

Next we categorized the RARγ-dependent proteins by measuring enrichment of NRs, and a comprehensive list of coregulators (coactivators (CoAs), corepressors (CoRs), mixed function coregulators (Mixed) or transcription and mRNA stabilization factors (TFs)(52)), as well as those coregulators that are mutated in PCa, known as so-called long-tail mutants(27) (**Supplementary Table 1A, B, C, Supplementary Figure 1B, C**). In 22Rv1-RARγ compared to LNCaP enriched NRs included the AR and the orphan receptor NR2C1/TR2 (**Supplementary Table 1B**). There was also enrichment of bookmarking factors, most clearly in 22Rv1-RARγ-TACC1 cells, including SMARCA4 and SMARCC1 (**Figure 2A**, salmon colored). Measuring group-level enrichment revealed that mixed function coregulators were highly enriched in basal 22Rv1-RARγ-TACC1 cells (**Supplementary Table 1A**). The chromatin remodelers in the long-tail mutants(11) were also significantly enriched in basal 22Rv1-RARγ-TACC1 cells (**Supplementary Table 1C**). Finally, although network enrichment unsurprisingly identified acetylation terms as enriched, other more novel terms included RNA binding and processing, including RNA splicing (**Figure 2B**, **Supplementary Figure 2**).

**Figure 2.**
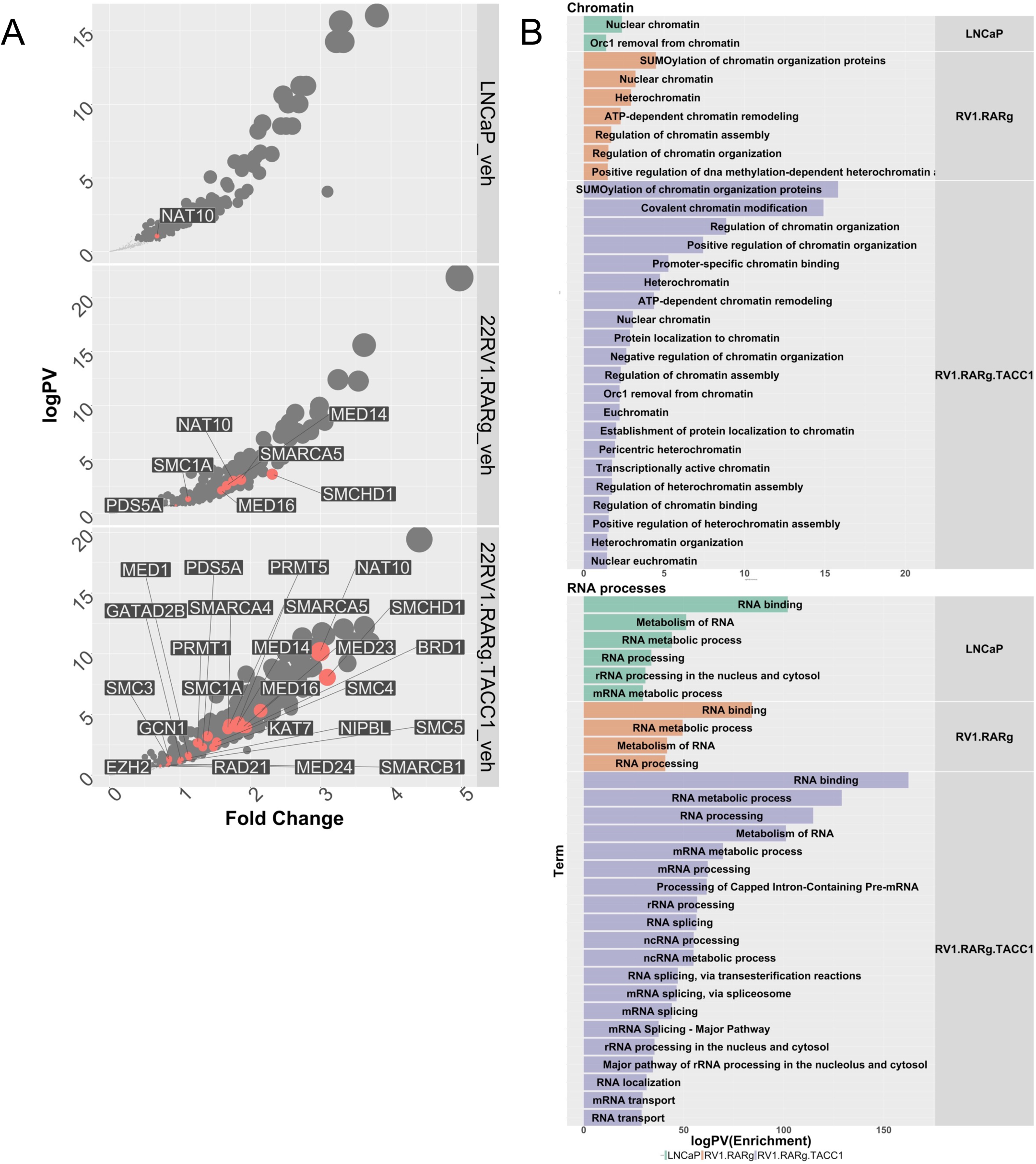
The RARγ complex in LNCaP and 22Rv1-RARγ isogenic variants. LNCaP, 22Rv1- mock, 22Rv1-RARγ and 22Rv1-RARγ-TACC1 cells were treated with CD437 (400 nM, 6h) or vehicle control in quadriplicates, and RIME analyses undertaken for RARγ in the cells and significantly different proteins were identified using an `edgeR` workflow to identify positively enriched proteins compared to IgG controls. **A.** Volcano plots depicting enrichment levels, compared to IgG controls (logPV > 1.3 & FC > .37), between the indicated cells in basal conditions. Proteins known to be associated with bookmarking functions (n=42) are indicated in salmon. **B.** The positively and significantly enriched proteins in each complex were analyzed by `StringDB` and the significantly enriched terms were grouped by the most frequently identified master terms (Chromatin, RNA Processes), and the significant terms are illustrated as waterfall plots.

Thus, the RARγ complex is significantly enriched with known and novel coregulators, and bookmarking and splicing factors as well as proteins involved with RNA-processing.

### MiR-96 targets are significantly enriched in the RARγ **complex**

To test whether the RARγ complex was enriched for miR-96 targets we used a biotinylated miR-96 approach (**Supplementary Figure 3**) to capture the repertoire of miR-96:mRNA interactions. We also aimed to capture miR-96 binding sites by testing their distribution in non-malignant HPr1AR and LNCaP cells, as well as examining the exposure time to miR-96, and the overlap with m6A sites. Finally, these approaches were coupled with miR-96 dependent transcriptomic and proteomic analyses.

The miR-96 targetome was highly variable. Naturally, more frequent miR-96 recognition elements MRE associated with greater numbers of genes (**Supplementary Table 2A, B**). Within cells, there was a significant overlap of MRE and m6A, for example the MRE significantly converged with m6A in HPr1AR at 48h (p=2.1e^-299^) and in LNCaP at 24h (p=1.8e^-14^). A 3’UTR enrichment of MRE was only pronounced when there were fewest MRE detected (**Supplementary Figure 4C**). More surprisingly there was only a modest overlap between HPr1AR and LNCaP cells in terms of either MRE or targeted genes (**Supplementary Figure 4A, B**). Interestingly, for protein-coding RNAs there was a 3’UTR bias of m6A-MRE in HPr1AR, whereas this was a 5’UTR bias in LNCaP (**Supplementary Figure 4D, 4E**). Categorization of miR-96 bound protein-coding genes revealed that TFs were significantly enriched in HPr1AR cells (**Supplementary Table 2C**). Examples of cell-specific MRE are illustrated for the bookmarking factor *SMARCC1* and the CoA *TACC1* (**Figure 3A**).

**Figure 3:**
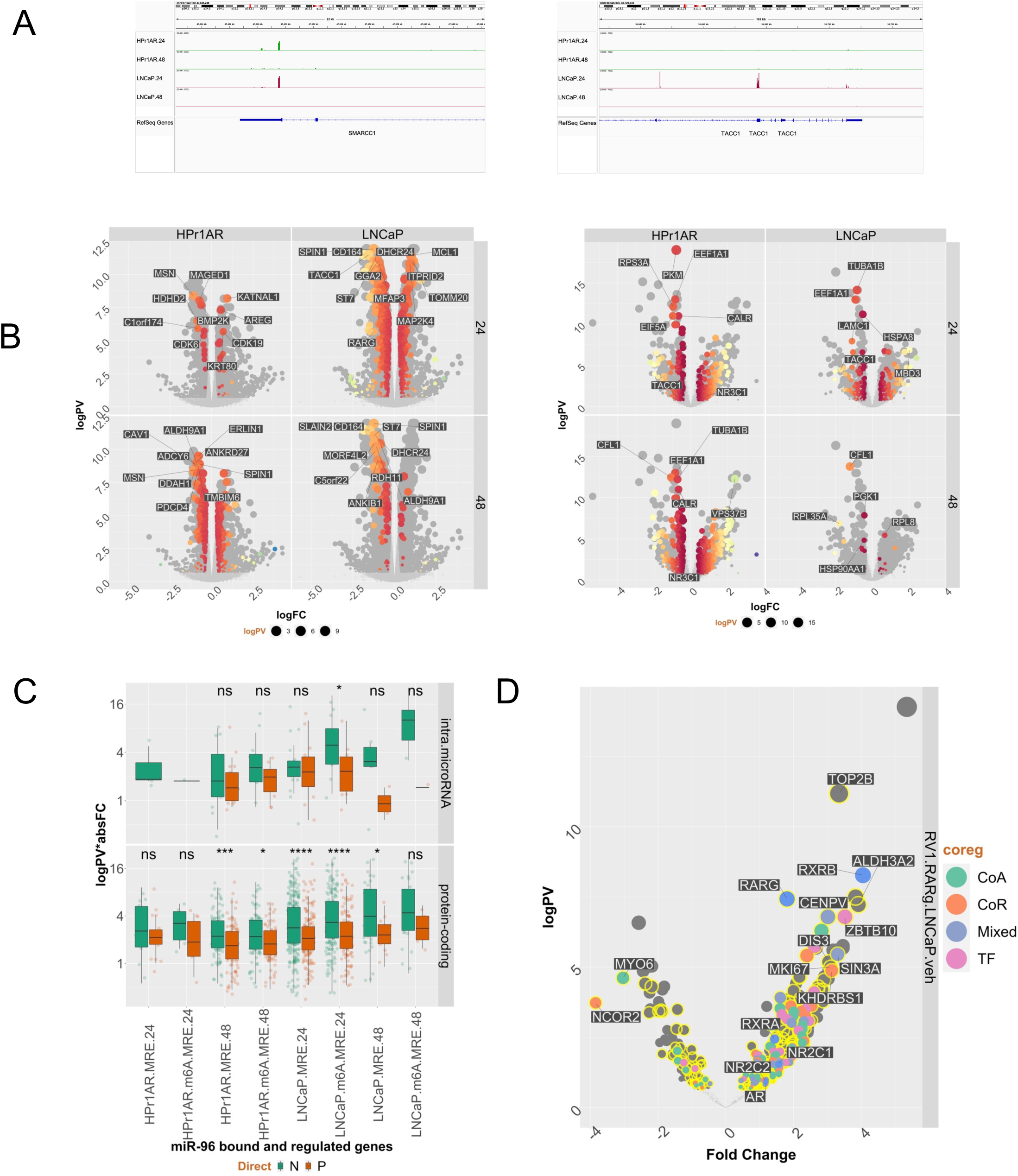
MiR-96 bound and regulated RNAs and proteins are frequently found in the RARγ complex in 22Rv1 cells. Non-malignant HPr1AR and LNCaP cells were treated in triplicate with 3’ biotinylated miRIDIAN miR-96 mimic or scramble control (50 nM, 24h and 48h) and IMPACT-Seq undertaken. RNA was captured from mid-exponential HPr1AR and LNCaP cells in triplicate for m6A-Seq. Briefly, `csaw` was used to identify differentially enriched regions of miR-96 binding sites (MREs) and m6A sites, and `bedtools` used to measure the overlap of m6A-MRE. Finally, HPr1AR and LNCaP cells were treated with miR-96 mimics (50nM, 48h) and RNA-Seq and Label free quantitative proteomics undertaken, and data processed with `edgeR` workflow to identify significant differentially enriched genes (DEGs) and differentially enriched proteins (logPV > 1.3 & absFC > .37). **A.** Representative genome browser views of MRE on *SMARCC1* and *TACC1* in HPr1AR and LNCaP cells. **B**. **Left.** Volcano plots of DEGs following miR-96 mimic treatment, and those that contained a MRE identified in the same cell background at either 24 or 48 h are shown in spectrum colors, with top 10 most significant genes labelled. **Right.** DETs were plotted as for DEGs. **C**. MRE or m6A-MRE-containing genes were classified as protein-coding or protein-coding genes containing an intronic miRNA, and the difference tested of the absolute fold change between negatively and positively-regulated targets. **D.** Volcano plot of significantly differentially enriched proteins from RIME data comparing 22Rv1-RARγ cells to LNCaP. Proteins were classified either as a Coactivator (CoA), Corepressor (CoR), Mixed function coregulator (Mixed) or transcription factor (TF) and whether they contained a MRE (yellow).

Next, we measured how MRE and m6A-MRE related to changes in mRNA and protein expression, by undertaking miR-96 mimic-dependent RNA-Seq and LFQ proteomics; PCA plots are shown in **Supplementary Figure 5**. Reflecting the MRE distribution, only ∼25% miR-96 dependent differentially expressed genes (DEGs) were common between HPr1AR and LNCaP, and 22Rv1 was most distinct. Fewer than 50% of all miR-96 mimic dependent DEGs or differentially enriched proteins (DEPs) contained a MRE identified in the same cell background. For example, in LNCaP at the mRNA level (**Figure 3B, left**) there were ∼470 MRE-containing DEGs, and in HPr1AR at the protein level (**Figure 3B, right**) there were 260 MRE-containing (DEPs). Furthermore, directly-bound miR-96 targets were regulated both positively and negatively, although MRE containing protein-coding genes in 48h in HPr1AR and 24h in LNCaP were significantly more down-regulated than up-regulated (**Figure 3C**), which appears to be enhanced in m6A-MRE (**Supplementary Figure 5B**). Classification of the DEGs with gene set enrichment analyses (GSEA) underscored the differential impact of miR-96 between cell lines, and all significantly enrichment terms contained a negative normalized enrichment score (NES) (**Supplementary Figure 5C**). For example, cell-cycle related terms and miR-96 targeted genes were unique to LNCaP.

Coregulator classification of MRE dependent DEGs and DEPs revealed in LNCaP the most negatively regulated MRE-containing CoA was *TACC1*, and amongst NRs *RARG* alongside the *AR* and *ESRRB* were bound and negatively regulated. At the protein level TACC1 was the most significantly regulated at both RNA and protein level, but only in LNCaP (**Supplementary Table 3A, B**). Selecting for miR-96 bound and negatively regulated genes in HPr1AR, LNCaP and 22Rv1 cells revealed 54 genes that included *RARG* and *TACC1* as well as *ALDH9A1*, *DHCR24* and *CAPNS1*, which are all related to retinoid functioning or signaling. Finally, enrichment for MRE-containing genes in the RARγ complex was most significant in the 22Rv1-RARγ variants (**Supplementary Figure 6, Supplementary Table 3C, 3D, 3E**, **Figure 3D**).

Together these data suggest that the choice of miR-96 binding sites is highly dynamic, and but that a consensus of directly bound and negatively regulated targets included RARγ, TACC1 and retinoid signaling associated targets is evident. In turn, these targets were most significantly enriched in the RARγ complex in 22Rv1 isogenic cell variants.

### The RARγ complex augments AR enhancer interactions

We measured how RARγ levels impact the AR cistrome using the 22Rv1 isogenic cell variants. Naturally, in 22Rv1-mock cells DHT treatment increased the size of the AR cistrome, but significantly larger basal and DHT-regulated AR cistromes were identified in 22Rv1-RARγ and 22Rv1-RARγ-TACC1 cells (**Supplementary Table 4A**, **Figure 4A**). Motif analyses(64) demonstrated a role for the RARγ-TACC1 complex to augment the AR cistrome (**Figure 4B**), in that vehicle treated 22Rv1-RARγ cells mirrored the effect of DHT on the AR cistrome in 22Rv1-mock cells. The presence of TACC1 led to an even more diverse motif repertoire (**Figure 4B**; **Supplementary Figure 7B**). H3K27ac cistromes were broadly comparable across the 22Rv1 isogenic variants but, again, motif enrichment and repertoire were increased RARγ (**Supplementary Figure 8A, 8B**).

**Figure 4:**
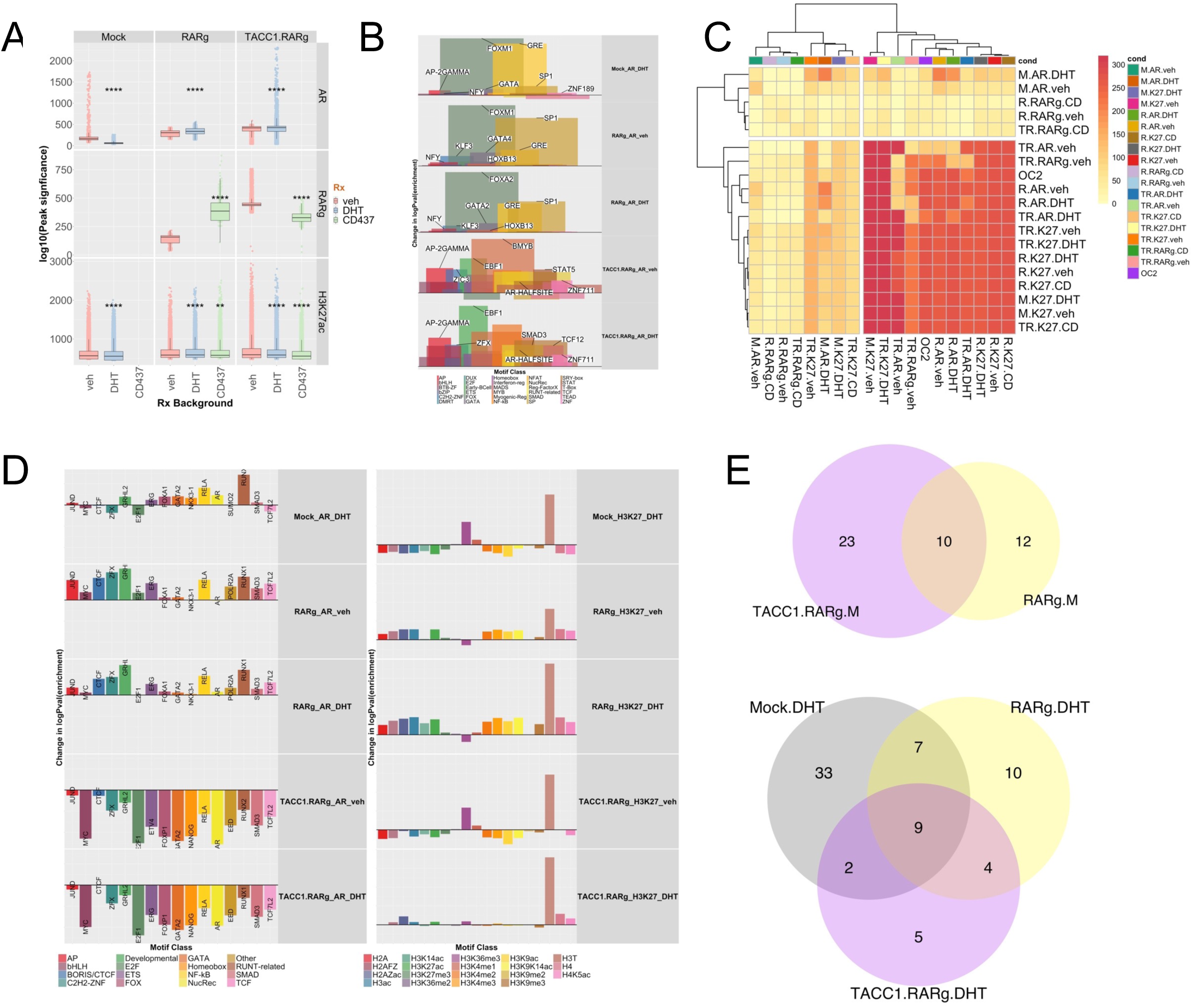
RARγ augments the shared AR and H3K27ac cistromes. 22Rv1-mock, 22Rv1-RARγ and 22Rv1-RARγ-TACC1 cells were treated with DHT (10nM), CD437(400nM) or vehicle for 6h and CUT&RUN undertaken using antibodies to either AR, GFP (for RARγ), H3K27ac or IgG in triplicate. Differential enrichment of regions compared to IgG controls (p.adj < .1) was measured with `csaw`. **A**. The mean of the log10(p.adj) of the significant peaks per cell/treatment condition are shown, and significant differences indicated. **B.** Motif analyses for AR, RARγ, H3K27ac cistromes was undertaken (`homer`) and motifs were grouped into the indicated transcription factor classes. For each motif the change in significance of enrichment was calculated across models compared to 22Rv1-mock cells treated with vehicle, from which within each class the mean delta logPval enrichment and mean percentage coverage calculated (indicated by peak width). The specific motif with the greatest delta per class is indicated. **C.** The genomic overlap with a minimum of 1 bp of the AR, RARγ, H3K27ac and ONECUT2 cistromes was measured between all cistrome pairs and significance determined with a Chi-squared test. From these analyses the -log10(p.adj) of the indicated overlaps was visualized as a clustered heatmap. **D**. The overlap of the AR, RARγ and H3K27ac cistromes with the cistrome collection in the CistromeDB was measured by `GIGGLE`. The change in enrichment was calculated for transcription factors and cofactors (**Left**) and histone modifications (**Right**) compared to 22Rv1-mock cells treated with vehicle. The factor with the greatest change is labelled. **E.** Super-enhancers were identified in the H3K27ac cistromes in the 22Rv1 variant using the `ROSE` algorithm also taking into account BRD4 cistrome from 22Rv1 cells as a pilot SE-enriched factor to generate high confidence sites. The Venn Diagrams indicate the number of unique and overlapping sites (minimum of 1bp) in the cells treated with either DHT of vehicle (M).

Next, we investigated the intersection of the RARγ, AR and H3K27ac cistromes (**Supplementary Figure 9**) and how these sites were shared with the 22Rv1 ONECUT2 cistrome (29), given that the ONECUT2 motif was enriched in the RARγ cistrome (31). Specifically, RARγ increased the significance of AR and H3K27ac overlaps (**Figure 4C**), whereas the overlaps between RARγ with AR only modestly changed. In a similar but opposite manner RARγ-TACC1 reduced the overlap between AR and ONECUT2 (**Supplementary Figure 10A, B**).

Overlap of the RARγ, AR and H3K27ac cistromes with those in the CistromeDB (54) (**Figure 4D**) also supported a role for RARγ to recapitulate the impact of DHT. For example, DHT treatment in 22Rv1-mock cells increased the overlap of AR with various TF including ETS (e.g. ERG), FOX (e.g. FOXA1), and RUNT-related (e.g. RUNX1), which was recapitulated in vehicle-treated 22Rv1-RARγ cells. Likewise, RARγ expression increased the diversity of H3K27ac cistrome overlaps with other histone modifications that are hallmarks of active transcription such as H3K36me3 (**Supplementary Figure 11**). These findings support the concept that the RARγ complex phenocopies DHT effects.

Extending these cistrome intersection analyses to ChromHMM-derived epigenetic states(65) again revealed that RARγ alone and with TACC1 shaped the AR cistrome by increasing enrichment at promoters and active enhancers (**Supplementary Table 4C**). These AR enriched enhancer binding sites were also more frequently associated with genes, for example for shared AR/H3K27ac binding sites in active enhancers (**Supplementary Figure 10C**). Finally, RARγ also notably increased the number and distribution of super enhancers (SE) in vehicle treated cells, and in the presence of DHT led to a different repertoire of SEs (**Figure 4E**). Annotation of SE-associated genes also supported a unique RARγ-mediated SE-dependent function to regulate other NRs, including NR4A1/NUR77 (**Supplementary Table 4C**).

These data support the concept that the RARγ complex phenocopies the impact of DHT treatment by increasing the number of AR binding sites, the repertoire of enriched motifs and shared TF binding. RARγ levels also directed the AR cistrome to active enhancers and promoters, and also directs the generation of unique SE sites.

### The RARγ complex bookmarks AR target genes and increases enzalutamide responses

Given the RARγ complex contained bookmarking factors and augmented AR enhancer interactions, we tested if RARγ exerts a bookmarking function for the AR during mitosis. The RARγ-dependent AR cistrome significantly overlapped with bookmarking factor cistromes (**Figure 5A**). RARγ also shaped the distribution of accessible chromatin, as measured by nucleosome-free (NF) regions that were enriched for H3K27ac (NF-H3K27ac). For example, there were ∼1950 unique RARγ-dependent NF-H3K27ac regions in 22Rv1-RARγ-TACC1 cells, and approximately 20% of these regions were shared by RARγ and AR (**Figure 5B**). There was a striking increase in the number of RARγ-dependent AR peaks activated by DHT in G_2_/M cells, further supporting a RARγ bookmarking function (**Supplementary Table 5, Figure 5C**).

**Figure 5:**
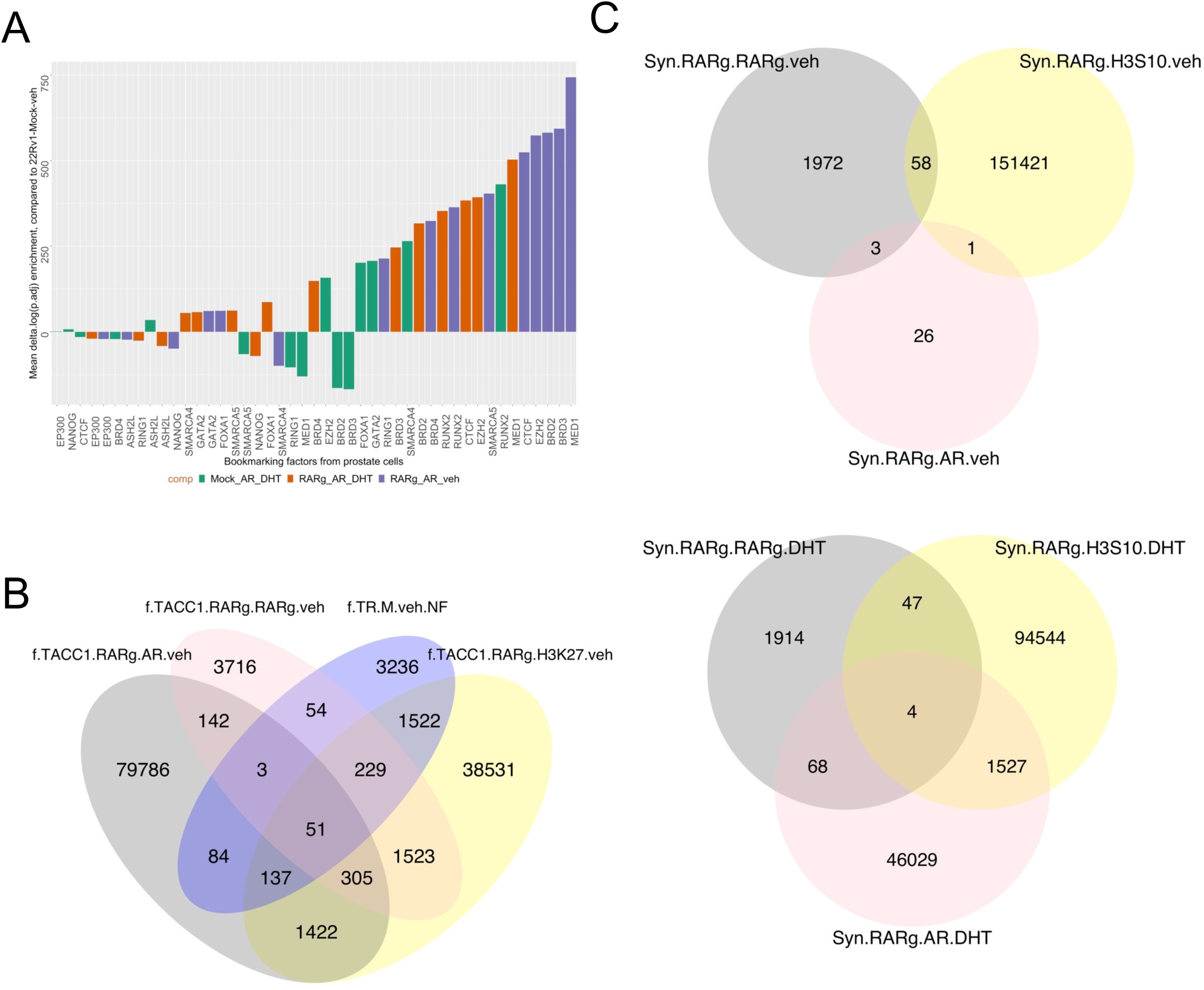
The role of RARγ to bookmark AR enhancers. **A.** In the 22Rv1 isogenic cell variants (in different colors) there was significant shared binding of AR cistromes with bookmarking factors as determined by `GIGGLE` analyses. **B.** ATAC-Seq was undertaken in triplicate in 22Rv1 cell isogenic cell variants and `ATACseqQC` used to define nucleosome free regions, and differential enrichment of chromatin accessibility measured with `csaw.` The significantly enriched nucleosome free regions (p.adj < .1) between 22Rv1-RARγ-TACC1 and 22Rv1-Mock cells (TR.M.veh.NF) were intersected with the indicated AR and RARγ cistromes also in 22Rv1-RARγ-TACC1 cells to generate the Venn diagrams of overlapping regions by a minimum of 1bp (`ChIPpeakAnno`). **C**. 22Rv1 isogenic cell variants were treated with nocodozole (60 ng/ml, 18h) or vehicle control, which led to ∼70% of cells in G_2_/M (labelled as “Syn”), and then treated with DHT (10nM) or vehicle for 6h and CUT&RUN undertaken using antibodies to AR, RARγ, H3S10P (as a marker of G_2_/M cells) or IgG in triplicate. Differential enrichment of regions compared to IgG controls (p.adj < .1) was measured with `csaw` and Venn diagrams generated of overlapping regions by a minimum of 1bp (`ChIPpeakAnno`).

We reasoned we could test the impact of a RARγ bookmarking function in AR-dependent transcriptional responses. Therefore, we measured the RARγ-dependent transcriptomic impact following treatments (DHT, ENZA or CD437) ((**Supplementary Figure 12A**, **Figure 6A**). GSEA was undertaken and ranking of all enriched phrases revealed the most common terms related to either NRs, epigenetics, and cell cycle; **Supplementary Figure 12B** is a waterfall plot of the most enriched NR terms across 22Rv1 isogenic variants. To reveal how the RARγ complex shifted the transcriptome we calculated the change in NES for the same terms in either the 22Rv1-RARγ or 22Rv1-RARγ-TACC1 cells compared to 22Rv1-mock cells (**Figure 6B**). This revealed that the RARγ complex shaped the DHT treatment enrichment terms, for example for estrogen-associated terms and, in 22Rv1-RARγ-TACC1 cells, terms associated with the synthetic retinoid, Tretinoin. Enza treatment positively enriched for tamoxifen-associated terms and negatively enriched for Tretinoin-dependent genes.

**Figure 6:**
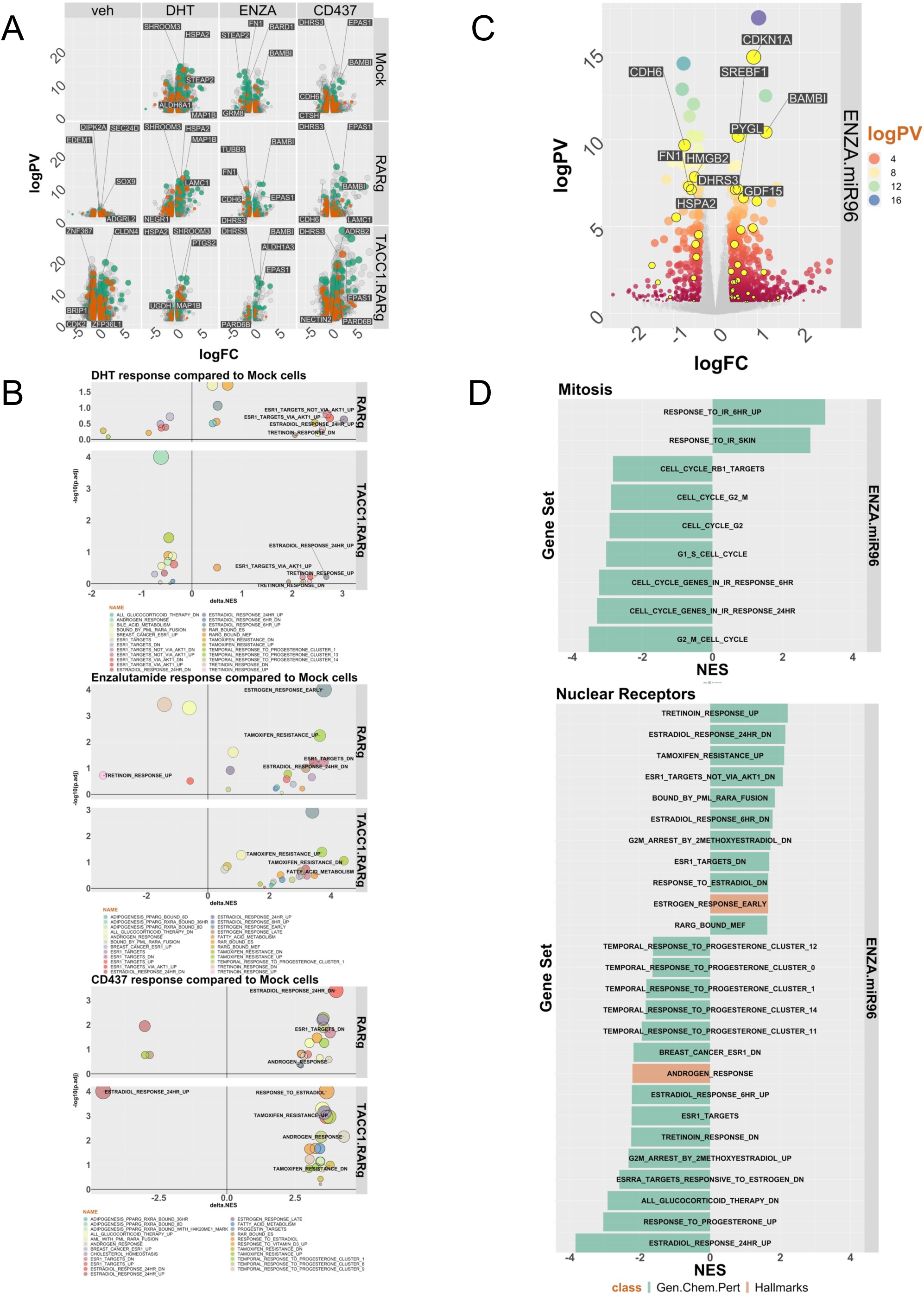
Expression of RARγ significantly shapes the DHT and Enza-dependent transcriptomes. 22Rv1 isogenic cell variants were treated with DHT (10nM), CD437 (400nM) or Enza (10μM) for 24h in triplicate and RNA-Seq undertaken. FASTQ files were QC processed, aligned to hg38 (Rsubread), and differentially expressed genes (DEGs)) (logPV > 1 & absFC > .37) identified with edgeR. **A.** Volcano plot depicting DEGs in 22Rv1 cell variants and treatments, with luminal (teal) and basal (brown) genes indicated. **B**. Pre-ranked gene set enrichment analysis (GSEA) was undertaken using the Hallmarks, and Chemical and Genetic Perturbations terms, and the most frequently enriched terms calculated (e.g. NRs, epigenetics and cell cycle terms). For the most frequent terms the delta normalized enrichment score (NES) between terms in either 22Rv1-RARγ or 22Rv1-RARγ-TACC1 cells compared to 22Rv1-mock cells are illustrated. **C**. Volcano plot depicting DEGs in 22Rv1 cells treated with Enza (10μM), miR-96 antagomir (100nM) or the combination. MiR-96 bound and regulated genes are indicated (yellow). **D**. GSEA analyses undertaken as **B**. and the most enriched terms in NRs or mitosis indicated in order of NES.

Further DEG classification (**Supplementary Table 6A**) revealed that prostate luminal gene signatures(66) were most frequently regulated, for example in 22Rv1-RARγ-TACC1 cells in responses to CD437, and to a lesser extent with Enza (**Supplementary Figure 12B**). Similarly, distinct cohorts of coregulators were regulated by DHT and NRs, as a class, were significantly enriched in all cell backgrounds (**Supplementary Table 6B**). However, miR-96 bound and regulated genes (termed miR96 BR; n∼660) were only significantly enriched in the DEGs from 22Rv1-RARγ and 22Rv1-RARγ-TACC1 cells (**Supplementary Table 6C**). Given that RIME analyses revealed enrichment of splicing factors in the RARγ complex (**Figure 2A**, **Supplementary Table 6D**) we also measured Differentially Expressed Transcripts (DETs). This revealed that the targets of alternative splicing were altered between cell backgrounds. For example, 22Rv1-RARγ cells there were significant changes in the mitotic spindle regulator SPAG5, and the AR itself. In support of this, in 22Rv1-RARγ cells there was reduced protein expression of ARv7 (**Supplementary Figure 13A, B**).

Finally, we tested if miR-96 antagomirs could sensitize 22Rv1 cells towards Enza, given that the RARγ complex was significantly enriched for miR-96 targets, and the RARγ complex shaped the AR cistrome and impacted the response to Enza. At the phenotypic level miR-96 antagomir plus Enza significantly decreased cell viability associated with a modest reduction of S-phase cells and an increase in necrotic cells at 48h (**Supplementary Figure 14A**). The co-treatment also impacted ∼1100 DEGs compared to individual treatments, including ∼50 MRE-containing miR-96 regulated genes (**Supplementary Figure 14B**, **Figure 6C** (yellow)). Again, GSEA analyses revealed enrichment of Estrogen and Tretoinin treatment terms, reflecting the impact of RARγ expression (**Figure 6B**), as well as genes regulated by the leukemic chimeric protein PML-RARA and bound by RARγ (**Figure 6D**). MiR-96 antagomir plus Enza impacted DETs were enriched for CoAs, notably including three SMARC genes (**Supplementary Table 7A, B**; **Supplementary Figure 15**).

Together these data further support a role for the RARγ complex to impact the AR functions through bookmarking the transcriptional functions, including to promote luminal differentiation gene programs and to enrich for estrogen and tretinoin dependent transcriptomes, as well as for miR-96 targeted genes. Reflecting the relationships between miR-96, RARγ and AR, a miR-96 antagomir enhanced Enza responses in 22Rv1 cells.

### The RARγ complex strengthens the AR cistrome-transcriptome relationships that antagonize ONECUT2 and associate with alternative lineages and metastasis

Collectively, these findings support an interplay between RARγ, AR, and ONECUT2. RARγ augments the AR cistrome (**Figure 4A**) and significantly overlaps with ONECUT2 (**Supplementary Figure 8**), whereas RARγ-TACC1 reduced the overlap between AR and ONECUT2 (**Figure 4B**). Therefore, we sought to test the impact of RARγ on the relationships between AR cistromes, DHT/Enza-dependent transcriptomes and the relationship to ONECUT2 functions.

The AR and H3K27ac shared binding sites that were driven by RARγ were more frequently annotated to Enza-regulated genes (**Supplementary Figure 16A**), and justified using a weighted cistrome-transcriptome approach (wt-C-T)(11,61,67) to measure how RARγ-dependent AR cistromes significantly impacted the regulation of DHT and Enza-regulated genes (**Supplementary Figure 16B**). This revealed that RARγ-TACC1 significantly increased the strength of the relationship between AR binding in Active Enhancers or Promoters and both basal and regulated gene expression (**Figure 7A**). These relationships were more common for upregulated gene groups but did include downregulated genes. Therefore RARγ-TACC1 not only augments AR binding but also enhances its gene-regulatory capacity.

**Figure 7.**
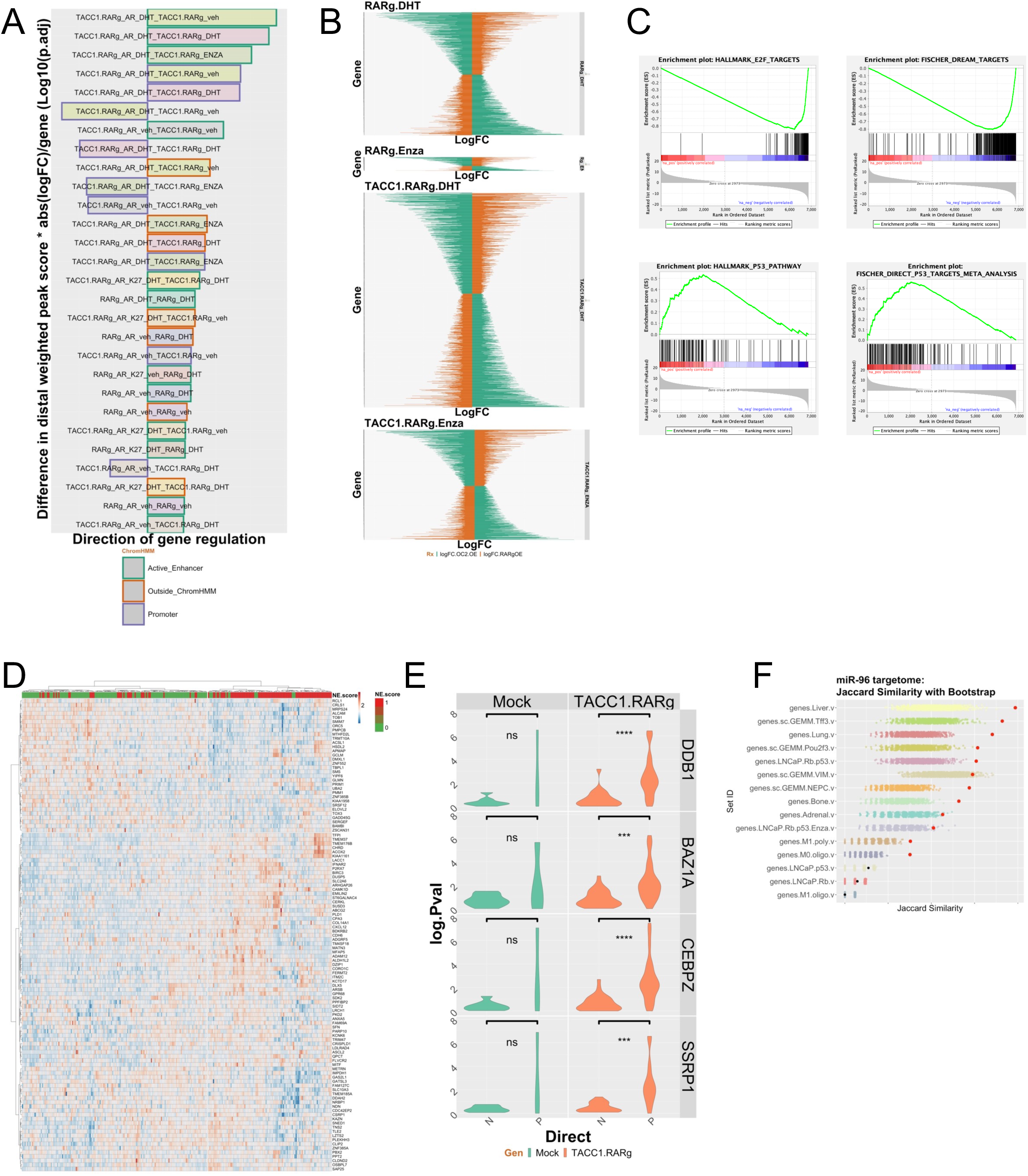
RARγ significantly determines AR cistrome-transcriptome relationships that are clinical significant. **A.** The cistrome in each cell was overlapped with ChromHMM defined epigenetic states and annotated to DEG within 100 kb, and the peak significance was summed for each gene, with a distal weighting, and multiplied by the absolute fold change for the same DEG. This score is the weighted cistrome-transcriptome (wt-C-T). A Wilcox test was used to compare the significant differences between the wt-C-T values to the comparable values in 22Rv1-mock cells. This revealed that the most significantly impacted cistrome-transcriptome relationships were TACC1-RARg-dependent. The name of bar refers to the cistrome first (e.g. TACC1_RARg_AR_DHT) and then the transcriptome it is matched to (e.g. TACC1_RARg_CD437). The direction of the bar refers to whether the genes were up or downregulated. The order refers to the most significance of change between the wt-C-T and that in 22Rv1-Mock cells. **B.** Antagonistic gene expression between either DHT or Enza regulated genes in 22Rv1-RARγ cells and 22Rv1-RARγ-TACC1 cells compared to gene regulation by over-expression of ONECUT2. Transcriptomic data in 22Rv1-RARγ and 22Rv1-RARγ-TACC1 cells was compared to that in 22Rv1 cells with ONECUT2 over-expression. Gene expression was defined as cooperative, if for a given gene the direction of change was the same between either DHT or Enza treatments and ONECUT2 over-expression, and antagonistic if the direction of change was opposite. The height of each panel is proportional to the number of events and indicates that in 22Rv1-RARγ-TACC1 cells there is more frequent Enza-dependent antagonism than in 22Rv1-RARγ cells. **C**. GSEA identified negative enrichment of E2F genes and positive enrichment of p53 genes in the Enza-regulated antagonized genes in 22Rv1-RARg-TACC1 cells. **D.** In SU2C tumors (n=293) upper quartile RARγ and lower quartile ONECUT2 were compared to the opposite and differential expressed genes determined. From these ∼ 3000 genes a lasso regression was used to identiffy genes that significantly associated with high/low (red/green) neuroendocrine feature. **E.** Partial correlation analyses in SU2C tumors between AR and RARγ-dependent AR-H3K27ac DHT regulated genes considering the impact of the indicated coregulators. The change in the correlation (delta.corr) was calculated as the difference between the Pearson correlation and Pearson partial correlations between AR and these target genes and each of the indicated coregulators. **F**. The miR-96 targetome of 663 MRE bound and miR-96 regulated genes were extracted from LNCaP cells to generate a ranked list. The similarity of these rank genes was compared to the ranked gene lists from the indicated experiments and similarity compared by Jaccard similarity measurements, and the significance determined by bootstrapping (n=1000). The symbols in red are significant similarity.

Next, we examined how these RARγ-dependent cistrome-transriptome relationships related to ONECUT2 functions by measuring the distribution of cooperative and antagonistic gene expression patterns between RARγ and ONECUT2 (29). This revealed a marked ability of Enza-regulated genes in the presence RARγ-TACC1 to antagonize ONECUT2-regulated genes (**Figure 7B**). By contrast there was no change in the frequency of cooperative genes (**Supplementary Figure 17A**). GSEA analyses of the antagonized Enza-regulated genes revealed positive enrichment for p53 targets, estrogen signaling and negative enrichment for E2F targets (**Figure 7C**). Genes that were antagonized between RARγ and ONECUT2 were significantly associated (logPV = 22.5) with unique AR binding sites, for example at the ONECUT2 locus.

Examining the relationships between RARγ, TACC1 and ONECUT2 in the SU2C cohort revealed *RARG* and *TACC1* positively correlated (Spearman’s correlation, .34; qval 2.6e^-6^) and *RARG* and *ONECUT2* negatively correlated, albeit modestly (-.2; qval=0.02). Differential expression analyses between upper quartile *RARG* and lower quartile *ONECUT2*, and *vice versa*, identified DEGs, which significantly related to high neuroendocrine score tumors (X^2^=8.5; pval = 0.003) (**Figure 7D**). We also measured how the RARγ complex impacted AR gene regulation and in the SU2C cohort and discovered that several coregulators including SSRPI(68) and BAZ1A(67), significantly and positively impacted the strength of the correlations between AR expression and AR enhancer dependent target genes (**Figure 7E**). Finally, we tested if miR96 BR genes (n∼660), and the RARγ-TACC1 and ONECUT2 antagonized genes (termed RARγvsOC2; n∼1400) were associated with PCa progression(69), metastasis(70), and alternative lineages(28). This revealed that both miR96 BR and RARγvsOC2 gene lists were enriched in different metastatic sites, notably liver, and enriched in both murine and human models with altered p53 and RB, including selectively enriched in different PCa lineages, most significantly in the Tff3 lineage (**Figure 7F, Supplementary Figure 17B**).

## DISCUSSION

The current study aimed to investigate crosstalk of the AR with other NRs to control cell lineage decisions. Specifically, we reasoned that nuclear resident Type II NRs could shape the actions of Type I NRs through a mechanism that centers on enhancer reprogramming. To investigate this mechanism, we dissected the relationship between RARγ and AR enhancer interactions and their targeted control by the oncomir miR-96 (31,37,71–73) through a series of high dimensional data approaches.

Composition analyses of the chromatin-bound RARγ complex in LNCaP and the three isogenic variants of 22Rv1 revealed the presence of well-established coregulators known to interact with NRs, including NCOR2 as well as the RXR heterodimeric partners of RARγ. A number of proteins were identified that have not previously been associated with RAR complexes, including the AR, alongside other proteins that impact AR function including NSUN2 which is required for translation of AR mRNA (74) as well as SART3, which is a splicing factor and regulates AR-dependent transcription(75) and has also been identified in leukemia as part of a fusion gene with RARG (SART3:RARG)(76). Reflecting these components, terms related to RNA processing and splicing process were enriched.

It is also clear that the RARγ complex in 22Rv1 cells, notably in the presence of TACC1, enhanced and diversified the coregulator repertoire and was enriched for bookmarking factors including SMARCC1 and SMARCA4. Recently(77), these BAF components have been demonstrated to undertake bookmarking in mitotic ES cells. Finally, the RARγ complex was significantly enriched with long-tail mutants including SMCHD1, which participates in DNA repair and binding(78). These components and coregulators were miR-96 targets. Although the miR-96 targetome was time-, cell-, and RNA-species specific, and impacted by coincidence with m6A, the targets of miR-96 were nonetheless enriched in the RARγ complex most significantly in 22Rv1-RARγ cells and 22Rv1-RARγ-TACC1 cells.

Next, we measured how RARγ levels impacted AR cistromes in the same three isogenic cell backgrounds, namely 22Rv1-Mock, 22Rv1-RARγ and 22Rv1-RARγ-TACC1 cells. The RARg cistrome was essentially unchanging in the different cell backgrounds, whereas RARg significantly enhanced the frequency of AR binding, for example at active enhancers, increased the repertoire of motifs enriched at these binding sites, the distribution of BRD4 positive super-enhancers (79,80), and increased how the AR and H3K27ac binding sites overlapped with other cistromes, notably including those for bookmarking factors. It is interesting to note that the AR cistrome is extremely flexible not only over the choice of binding site, but also the number of sites. For example, a recent study on primary PCa revealed that across tumor samples the AR cistrome ranged from ∼800 sites to more than 60,000 per sample, indicating considerable tailoring of the AR cistrome(81). The shift in AR cistrome overlap with other cistromes included both gain and loss, and indeed RARγ and TACC1 together reduced the significance of the overlap of ONECUT2 and AR. Together these data suggest that the impact of RARγ on AR is to phenocopy the effect of DHT exposure.

We reasoned, given the inclusion of bookmarking factors in the RARγ complex, that one function of this Type II NR is to bookmark AR binding sites in mitotic cells and, indeed, cells in G_2_/M were acutely sensitive to RARγ expression which markedly increased the DHT-dependent AR cistrome. Indeed RARγ directly and indirectly impacted ∼1500 accessible chromatin sites that were shared with H3K27ac in a RARγ-dependent manner but were not RARγ bound. We reason that these sites had been previously marked by RARγ or another bookmarking factor that is downstream of RARγ genomic functions. These findings fit with an emerging appreciation for other Type II NRs to initiate bookmarking in pluripotent cells(19,82,83), but to date neither bookmarking in general nor the functions of NRs to mediate this process have been investigated in prostate or other cancers.

The RARγ-dependent shift in AR enhancer interactions was measured at the transcriptomic level, where it led to significantly stronger cistrome-transcriptome relationships. Specifically, RARγ-TACC1 enhanced and redirected AR enhancer usage to impact DHT-dependent luminal differentiation programs, as well as the transcriptional responses to Enza, including the distribution of splice variants and notably reduced expression of the AR variant AR-v7. Indeed, we also demonstrated that Enza sensitivity was increased with an miR-96 antagomir.

The interplay between the RARγ complex, AR and ONECUT2 appears complex. RARγ and AR significantly overlap with the sites of ONECUT2 binding, but RARγ-TACC1 reduced the significance of these overlaps and increased the level of antagonistic control of gene expression between Enza and restored ONECUT2 expression. One explanation for this antagonism could be the epigenetic status of shared binding sites. Low density CpG methylation at enhancer regions can both repress and attract different classes of TFs. Of relevance here, increased CpG methylation attracts RARγ binding and repels ONECUT2(84); therefore, the status of CpG at enhancer sites will impact the functional antagonism between RARγ and ONECUT2. It is worth noting that RIME revealed that the RARγ complex in 22Rv1-RARγ-TACC1 cells was enriched for the DNA methyltransferase DNMT1. These findings are in keeping with the concept that changes in CpG methylation levels at enhancer regions are highly dynamic, critical in development(85,86) and appear to promote and sustain cell differentiation(87–94).

The genes that are either bound and regulated by miR-96 or antagonized between RARγ and ONECUT2 were significantly enriched in gene expression patterns associated with PCa progression and alternative lineages. Indeed, comparing tumors with high RARγ and low ONECUT2 to the opposite revealed gene signatures that were significantly enriched in the neuroendocrine phenotype.

In summary the current study supports a model whereby the RARγ complex exerts a bookmarking function for AR enhancers to shape cistromic functions, and in turn is regulated by miR-96 targeting of the RARγ complex. Androgen-indifferent lineages evoked by lineage plasticity drivers, such as ONECUT2, may be prevented by combinatorial strategies that induce RARγ and maintain RARγ complex bookmarking factors in favor of luminal differentiation programs at competing regulatory regions necessary for AR functions.

## DATA AVAILABILITY

All data are available (GSE254731)

## FUNDING

*MJC* also acknowledges National Institute of Health Cancer Center Support Grant (P30CA016058) to the OSUCCC The James. *MJC* also acknowledges institutional support from Cedars-Sinai Cancer, Cedars-Sinai Medical Center. *SAW*, *JSG* and *SH* each acknowledge support from Pelotonia at OSUCCC The James for individual fellowships.

## ACKNOWLEDGEMENTS

Consent for publication

Not applicable

## Availability of data and materials

The datasets generated and/or analyzed during the current study will be available on GEO

## Competing interests

The authors certify that they has NO affiliations with or involvement in any organization or entity with any financial interest (such as honoraria; educational grants; participation in speakers’ bureaus; membership, employment, consultancies, stock ownership, or other equity interest; and expert testimony or patent-licensing arrangements), or non-financial interest (such as personal or professional relationships, affiliations, knowledge or beliefs) in the subject matter or materials discussed in this manuscript.

## Authors’ contributions

*SAW* undertook IMPACT-Seq, m6A-Seq, RIME, CUT&RUN, ATAC-Seq, RNA-Seq and cell biology experiments; *SH* undertook synchronized CUT&RUN and cell biology experiments; *JSG* contributed to ATAC-Seq and cell biology experiments; *DN* undertook to cell biology experiments and contributed to manuscript preparation; *HT* undertook processing of genomic data; *LP* contributed to data interpretation and manuscript preparation; *MDL* undertook GIGGLE analyses; *MS* contributed to cell biology and genomic experiments; *CJM* contributed to data interpretation and manuscript preparation; *LESC* contributed to constructing the genomic workflow and data interpretation; *MF* contributed to interpretation of genomic findings in the context of advanced PCa; *SAW* and *MJC* jointly conceived of the study design; *MJC* oversaw the implementation of the study and undertook all other bioinformatic analyses, generated tables and figures and led data integration and manuscript preparation.

## Notes

### Competing Interest Statement

The authors have declared no competing interest.

### Summary of Updates

Updated figures and text

