## Supplementary material for "Epigenetic disruption of the RARγ complex impairs its function to bookmark AR enhancer interactions required for enzalutamide sensitivity in prostate cancer": Supp Figures

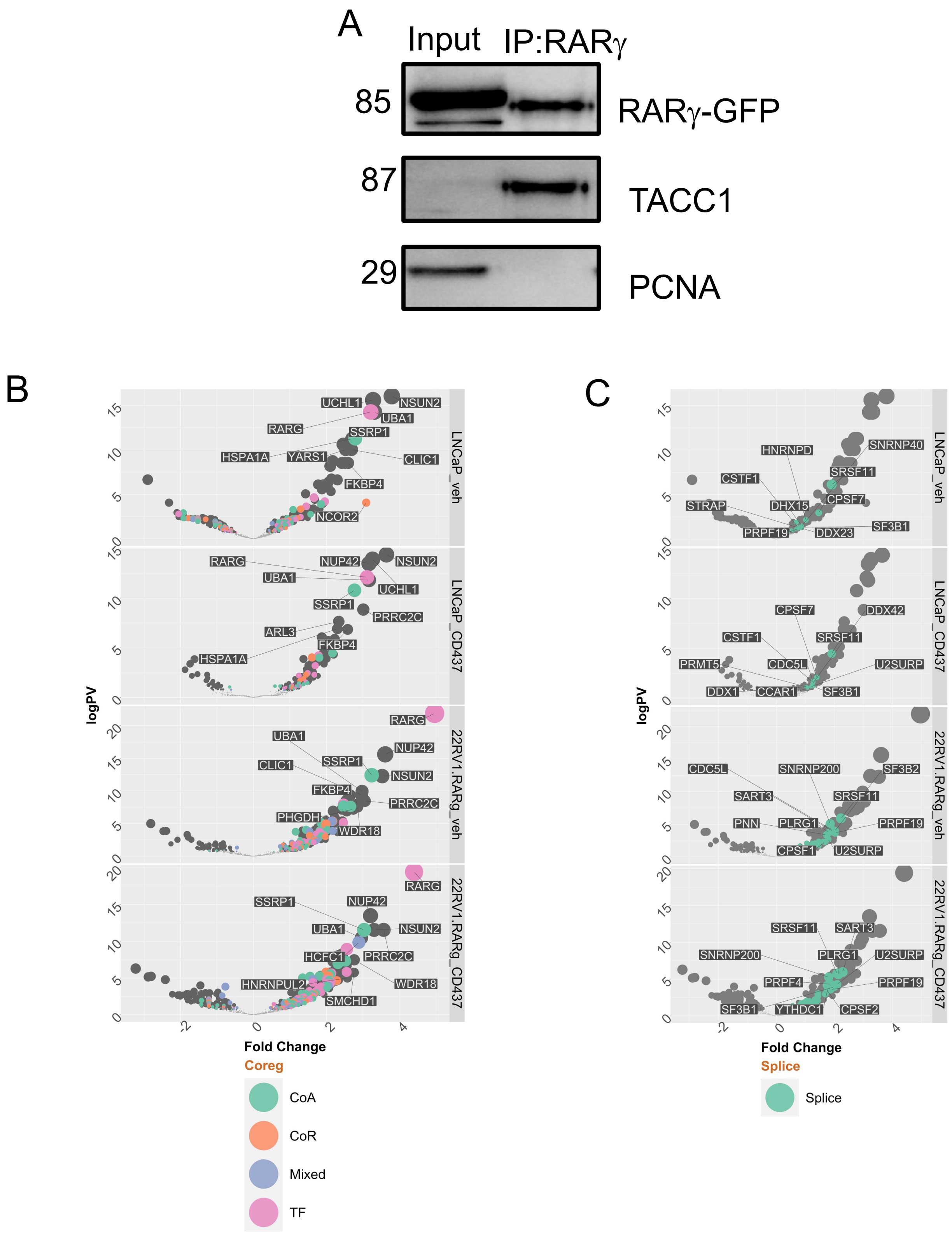


Supplementary Figure 1

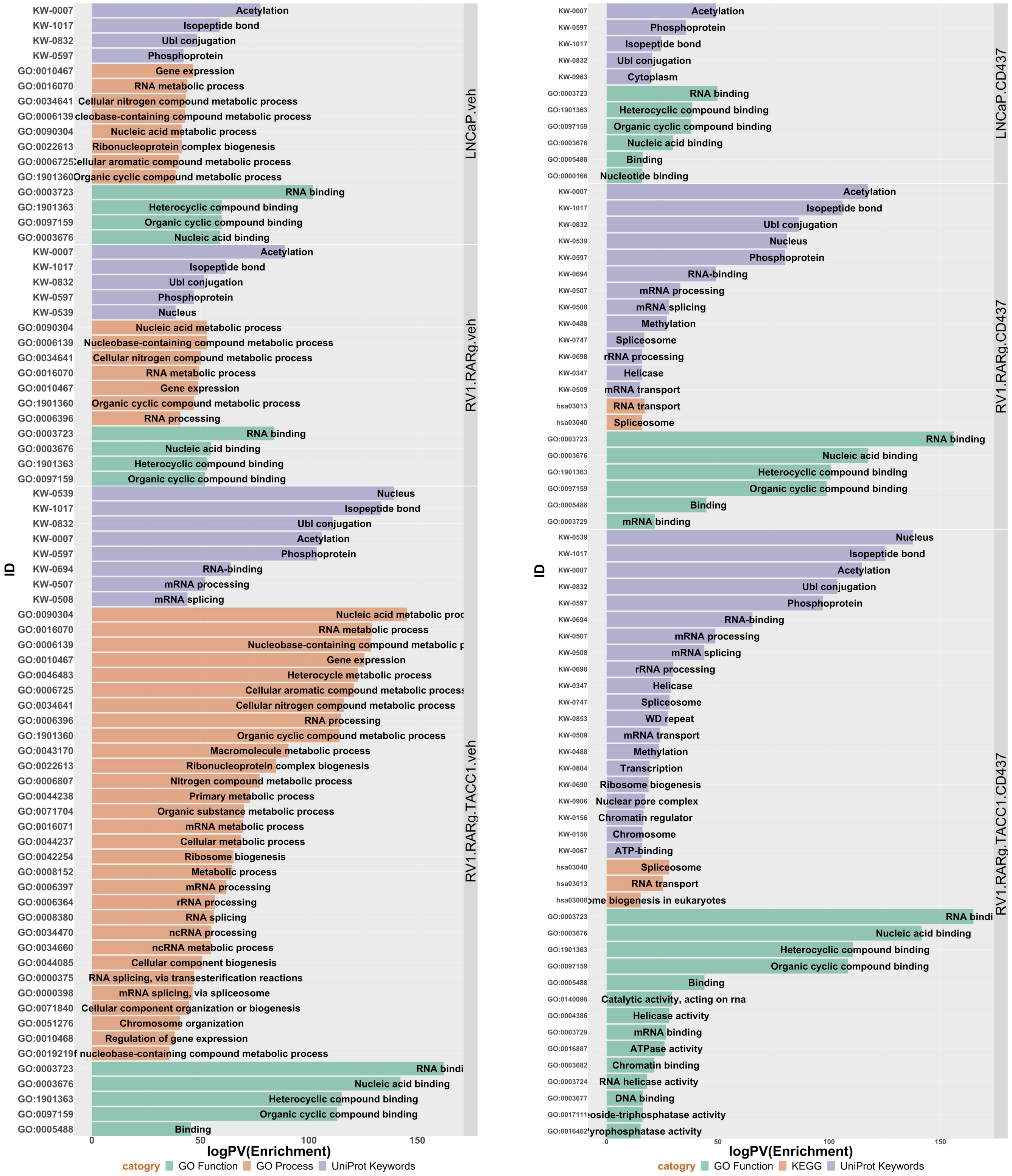


Supplementary Figure 2

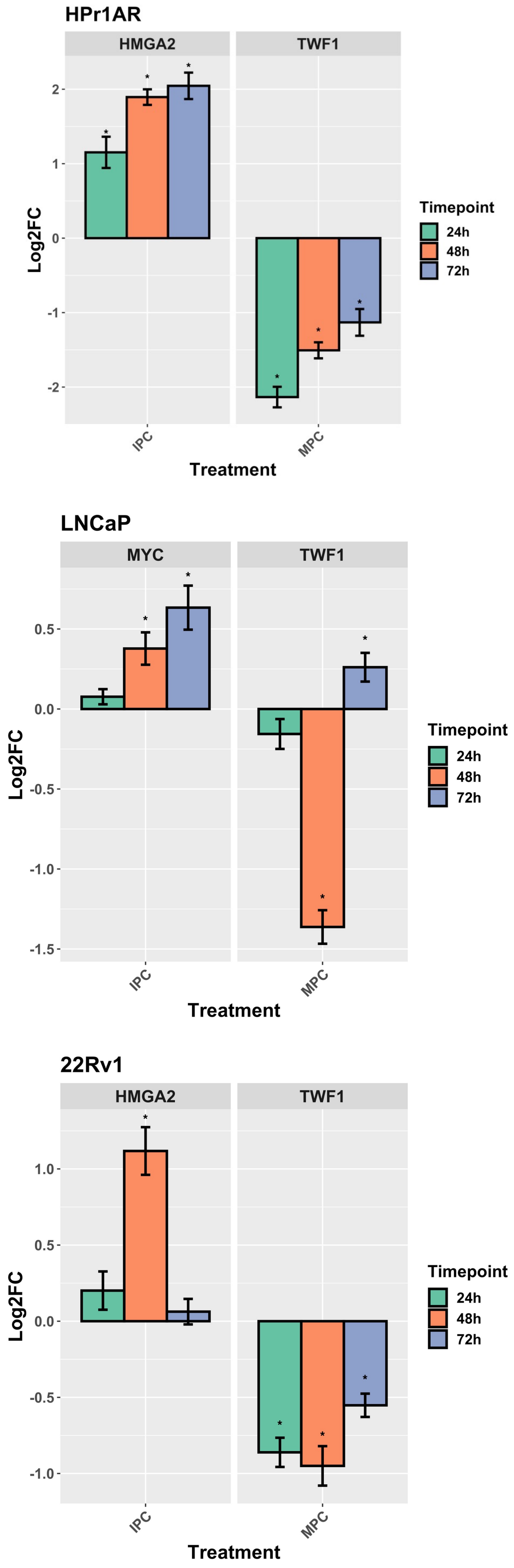


Supplementary Figure 3

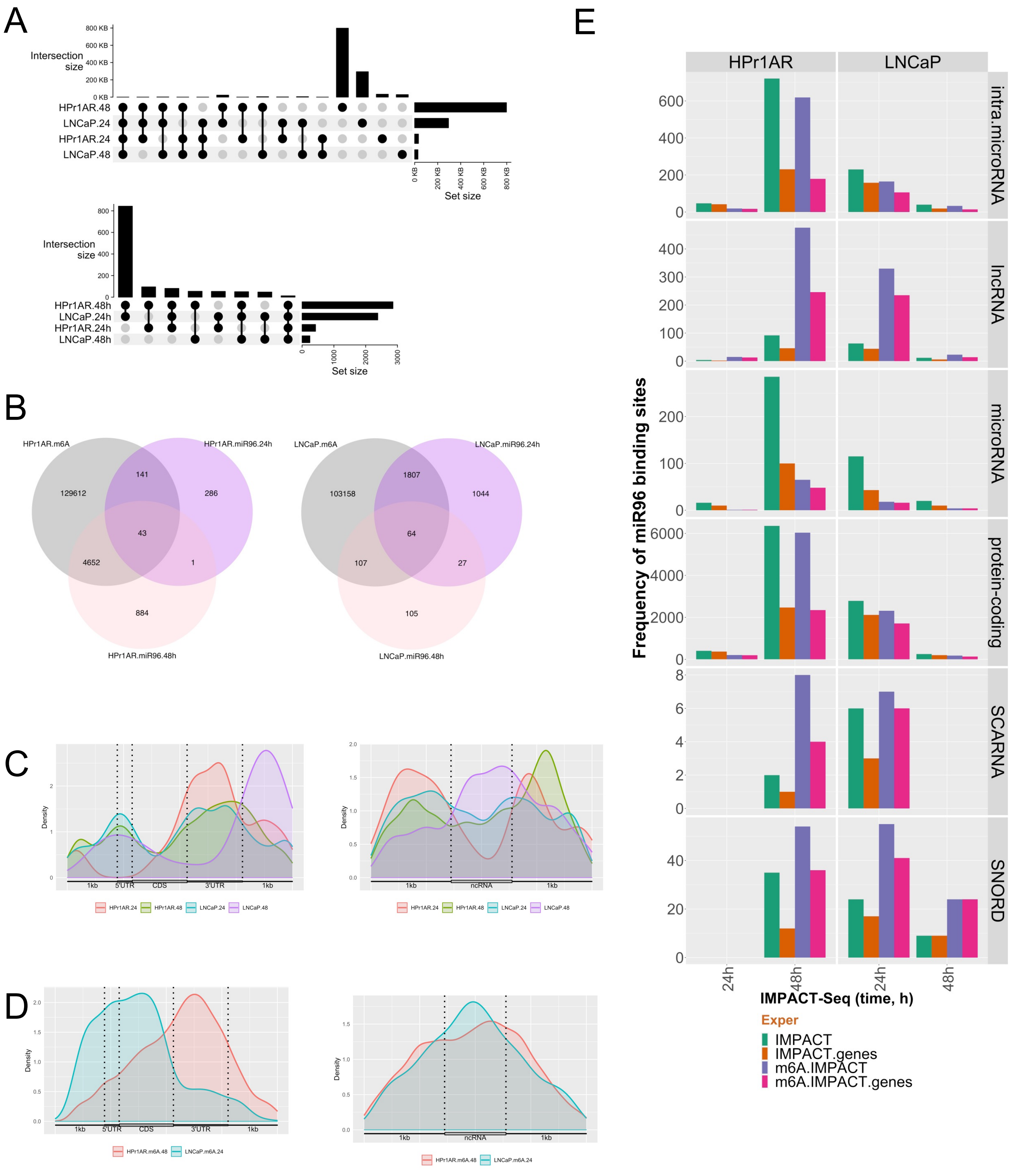


Supplementary Figure 4

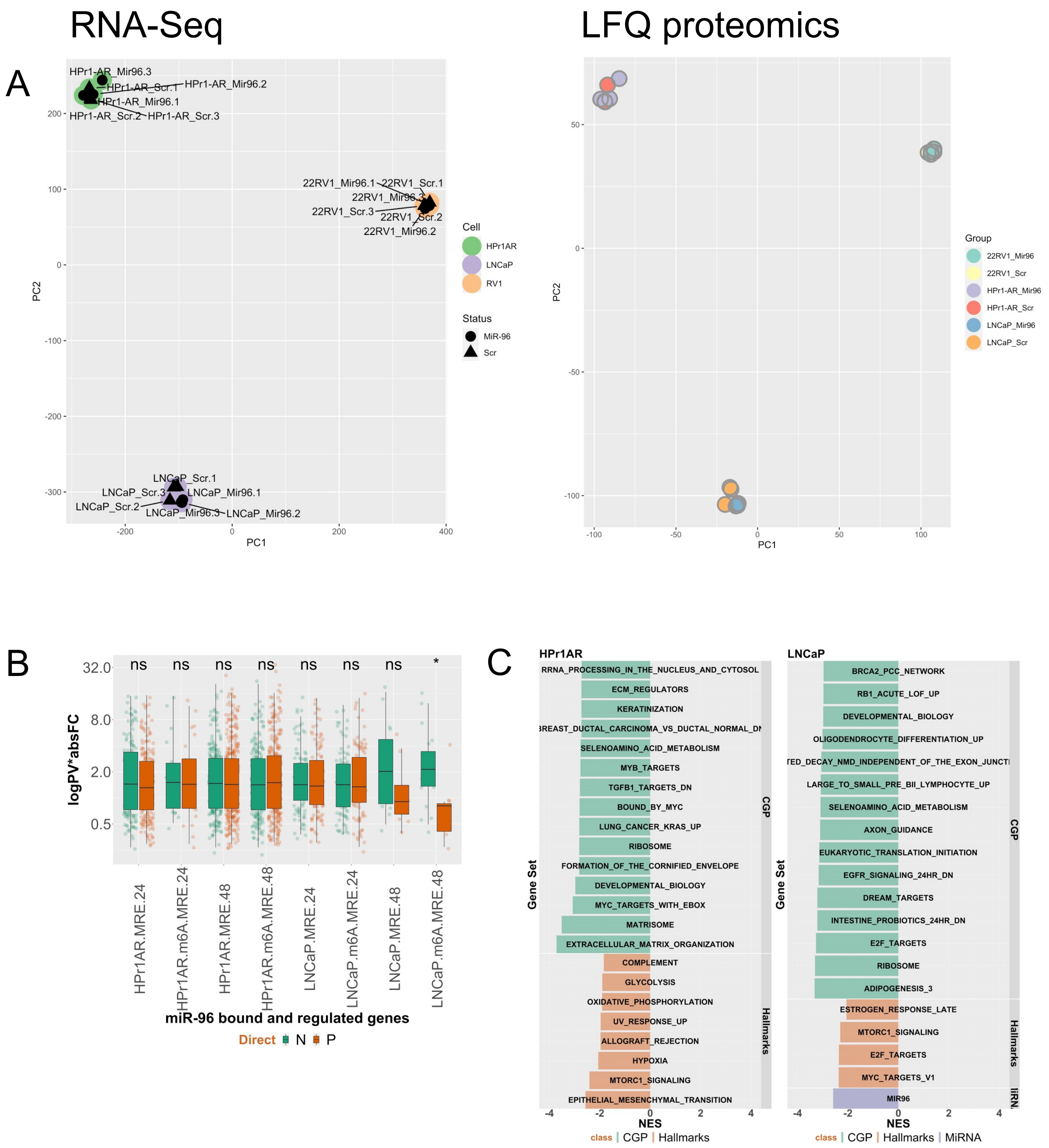


Supplementary Figure 5

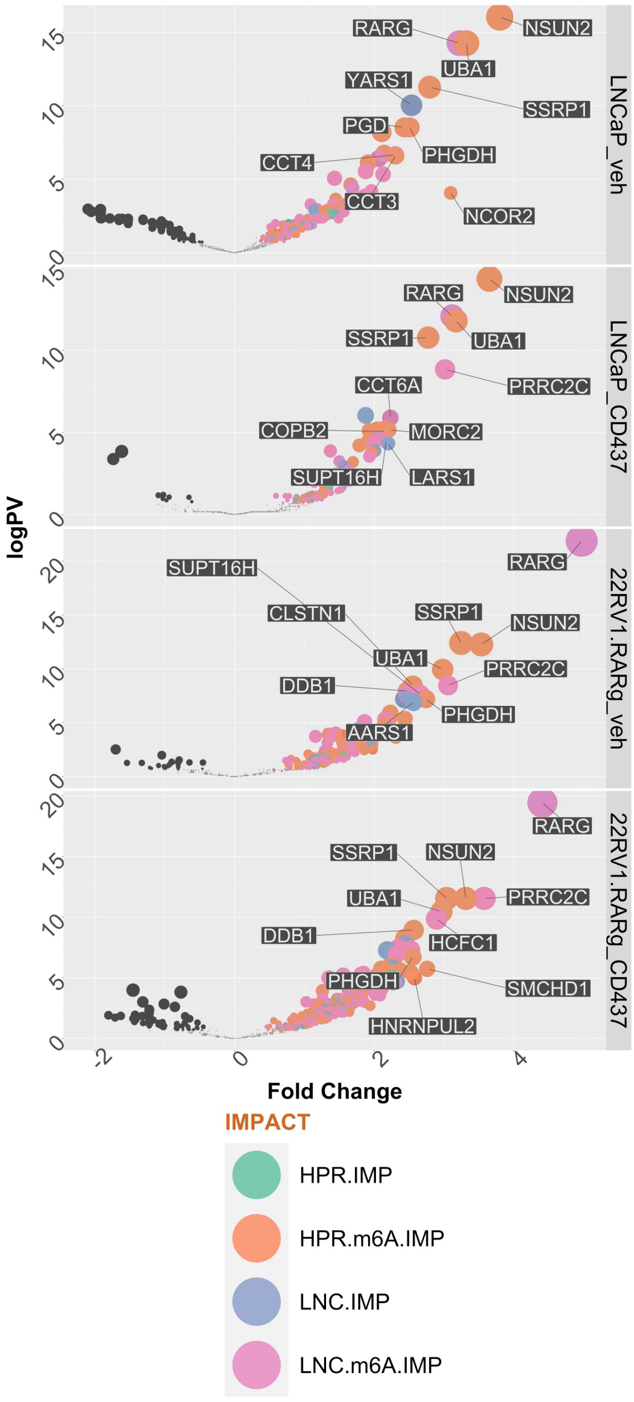


Supplementary Figure 6

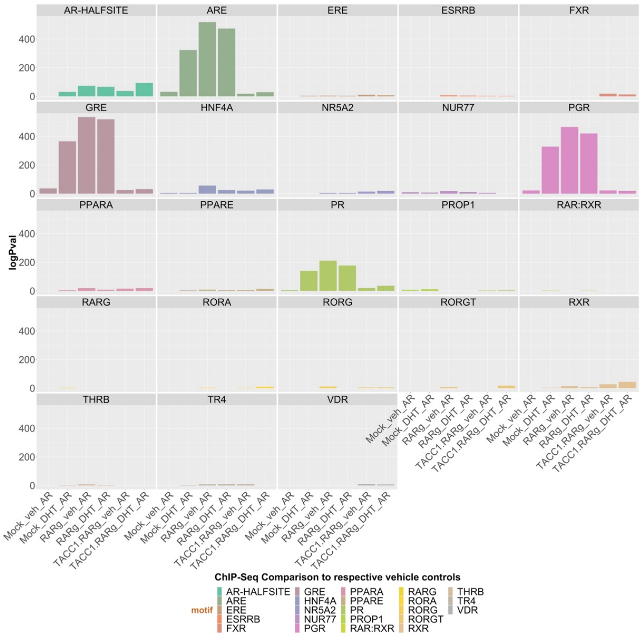


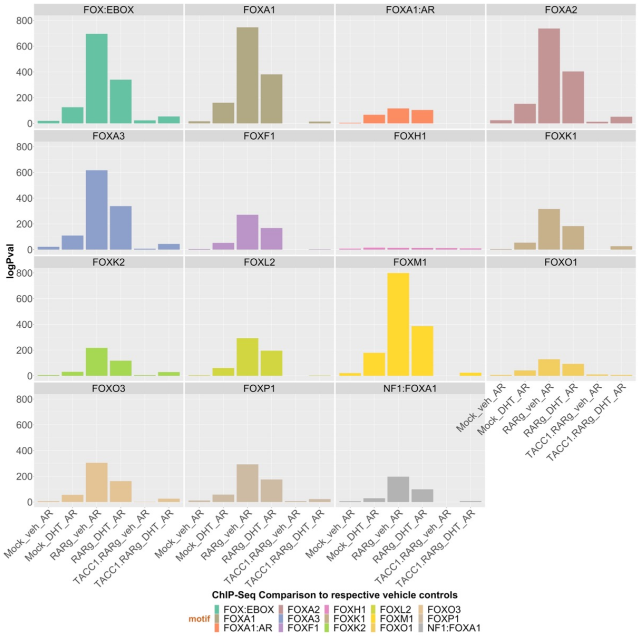


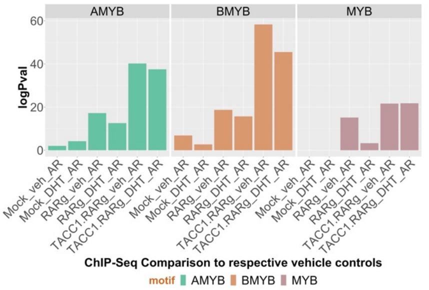


Supplementary Figure 7

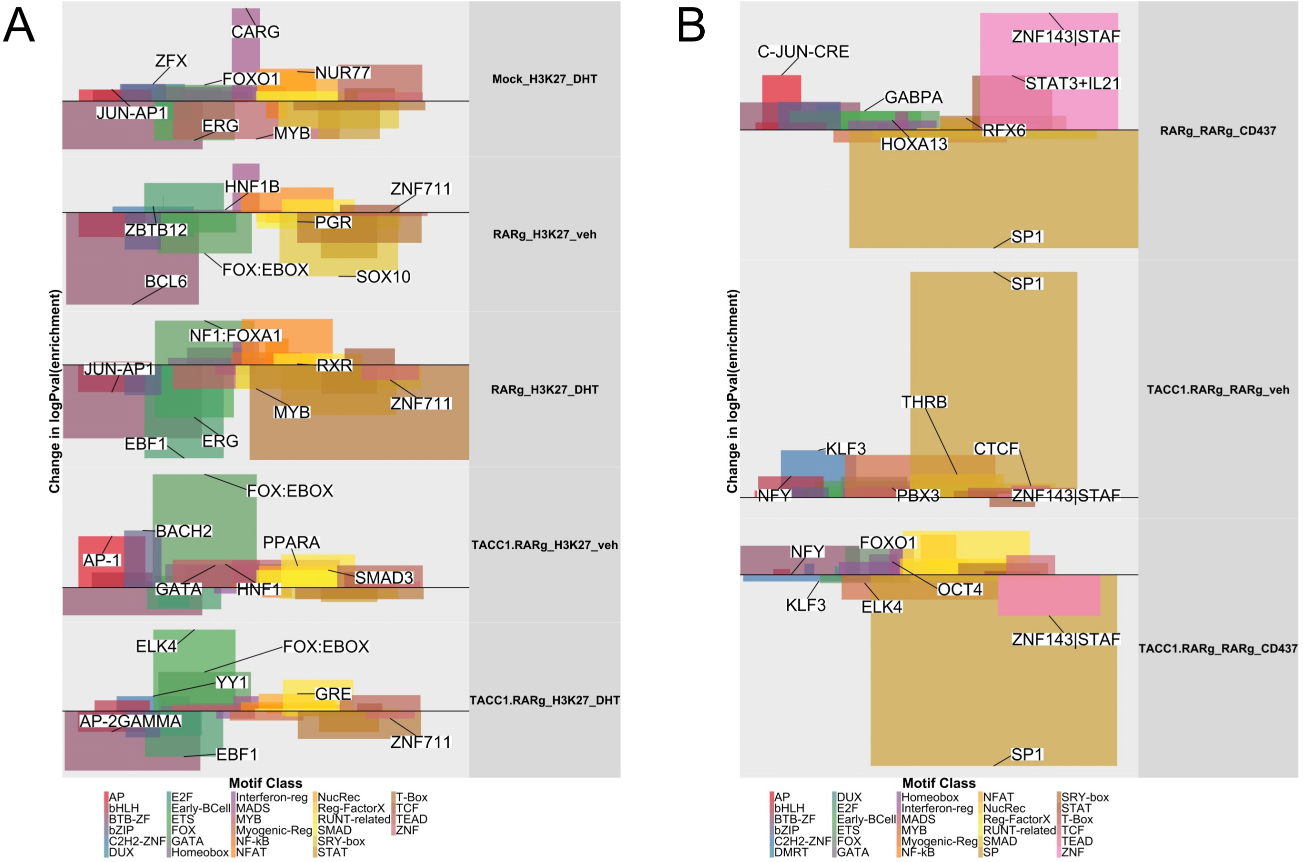


Supplementary Figure 8

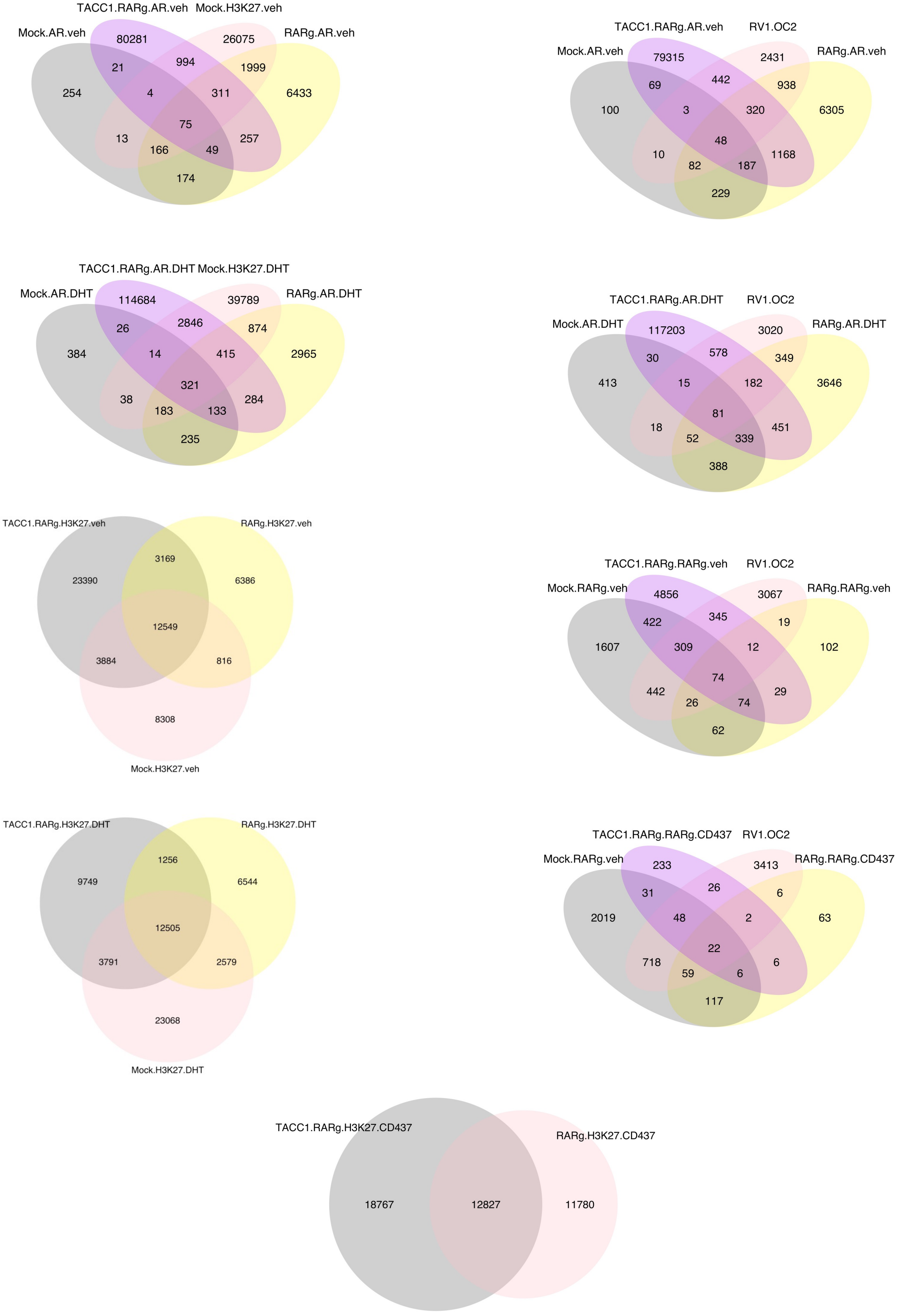


### Supplementary Figure 9

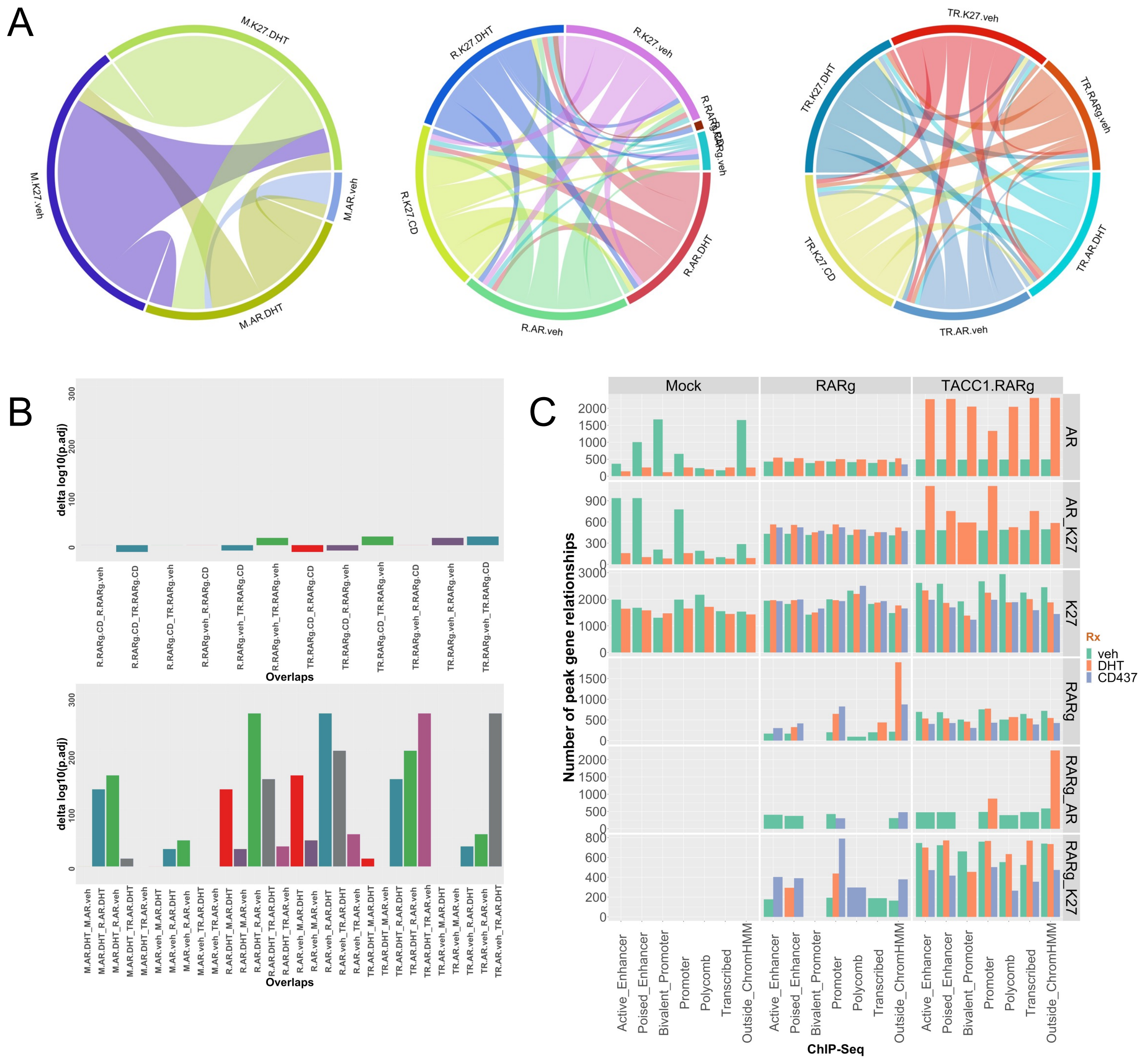


Supplementary Figure 10

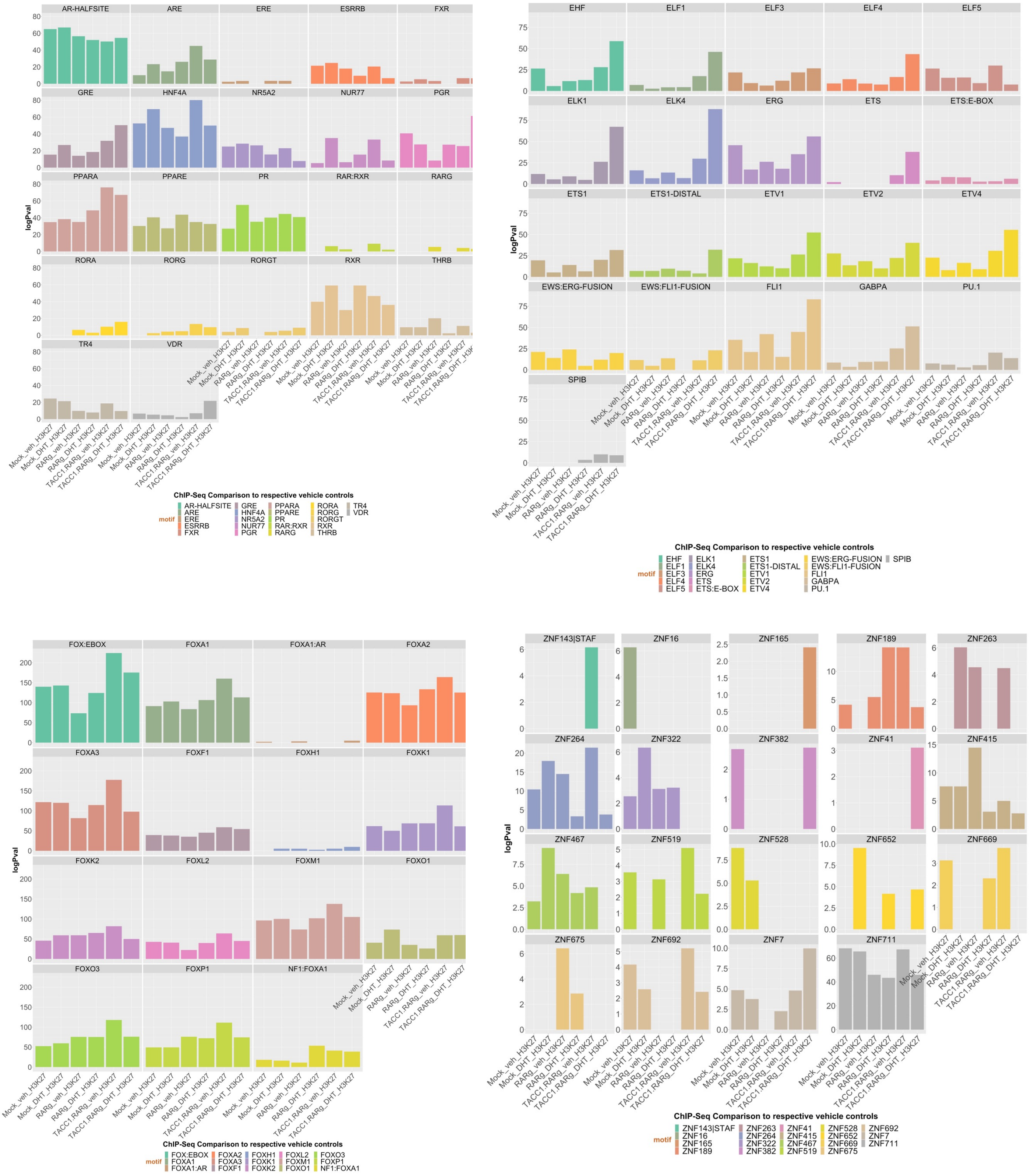


### Supplementary Figure 11

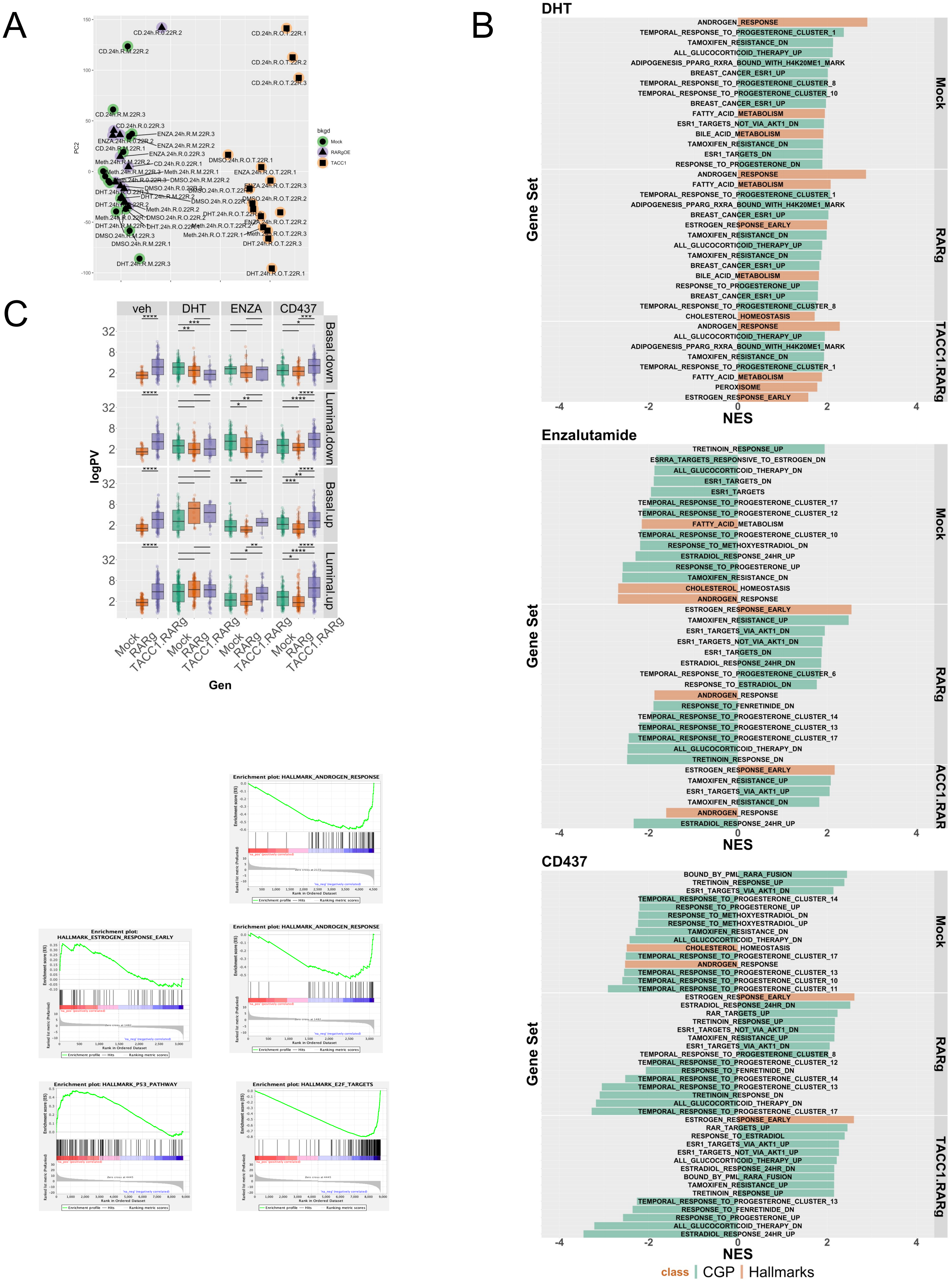


Supplementary Figure 12

A

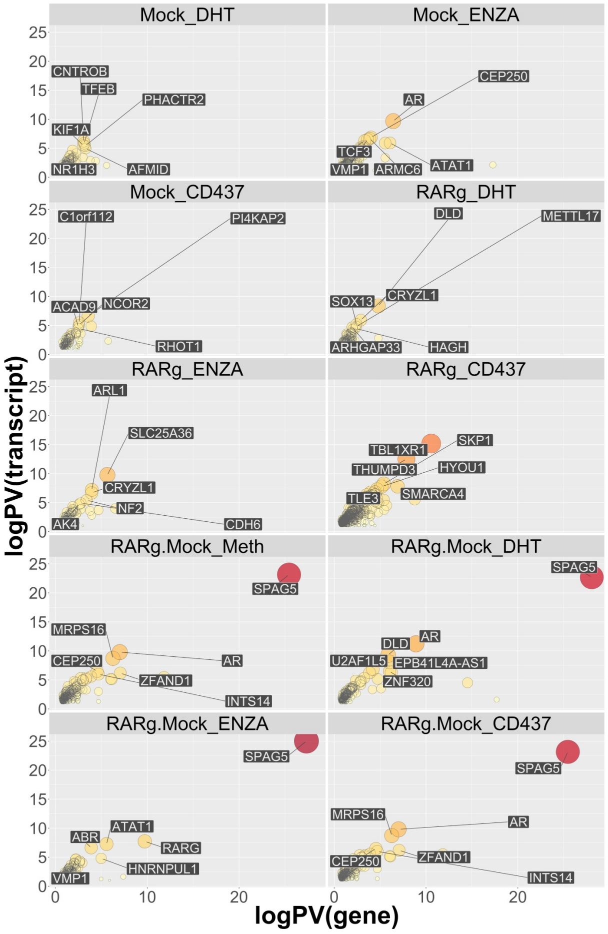


B

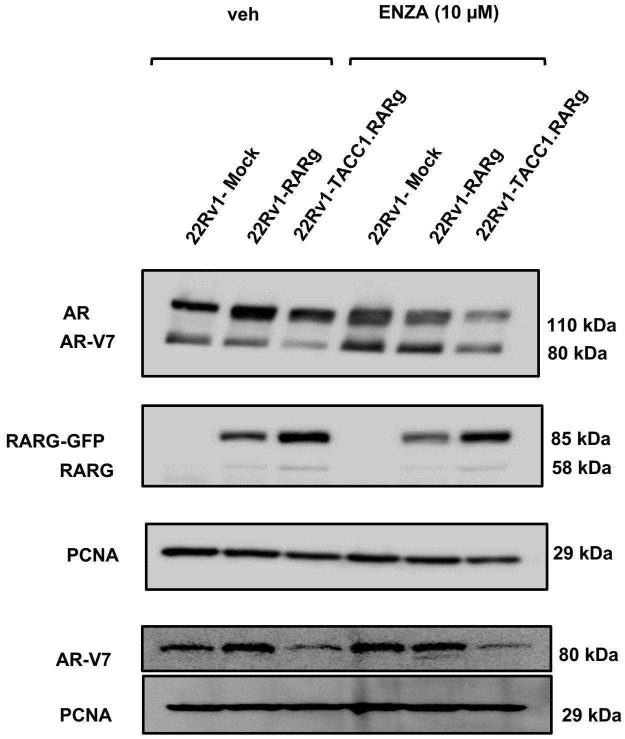


Supplementary Figure 13

A

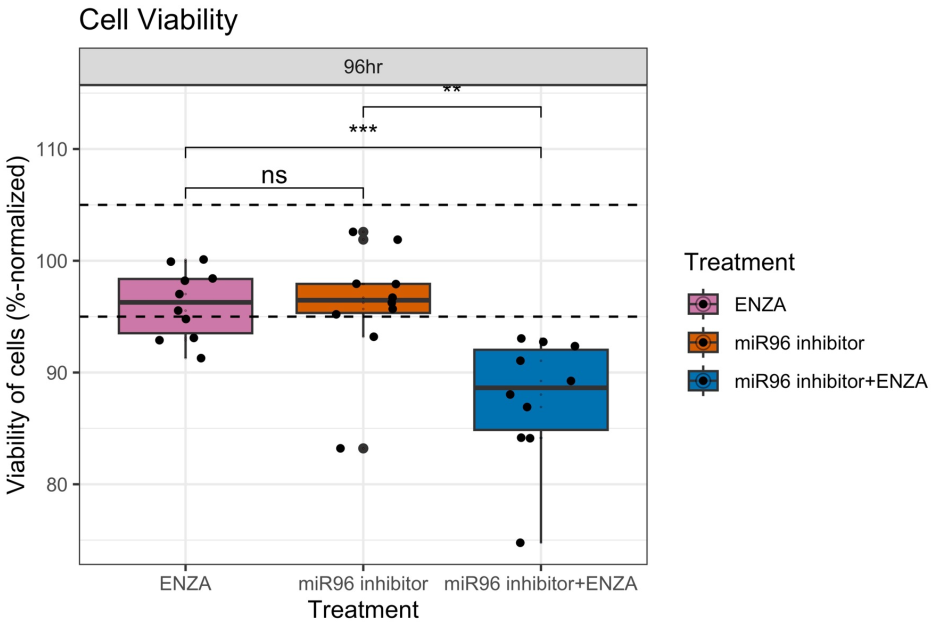


B

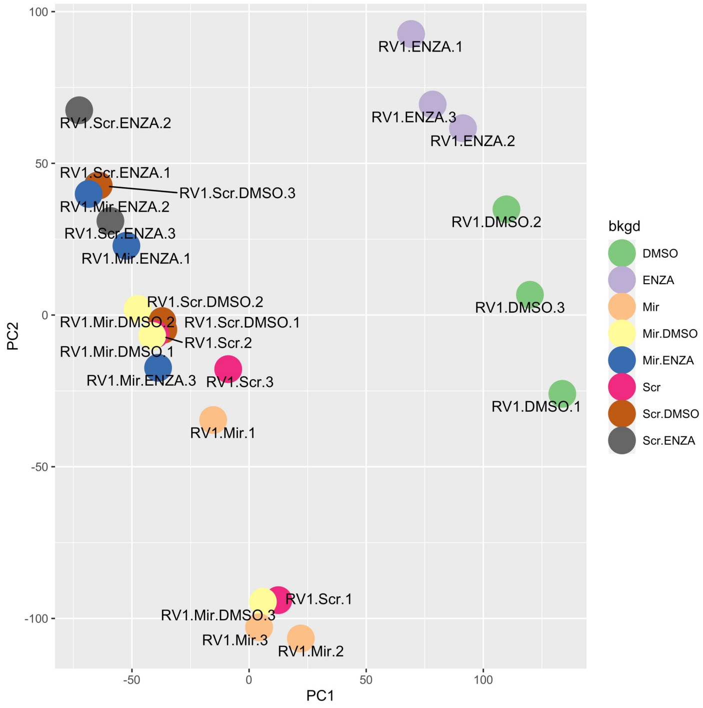


Supplementary Figure 14

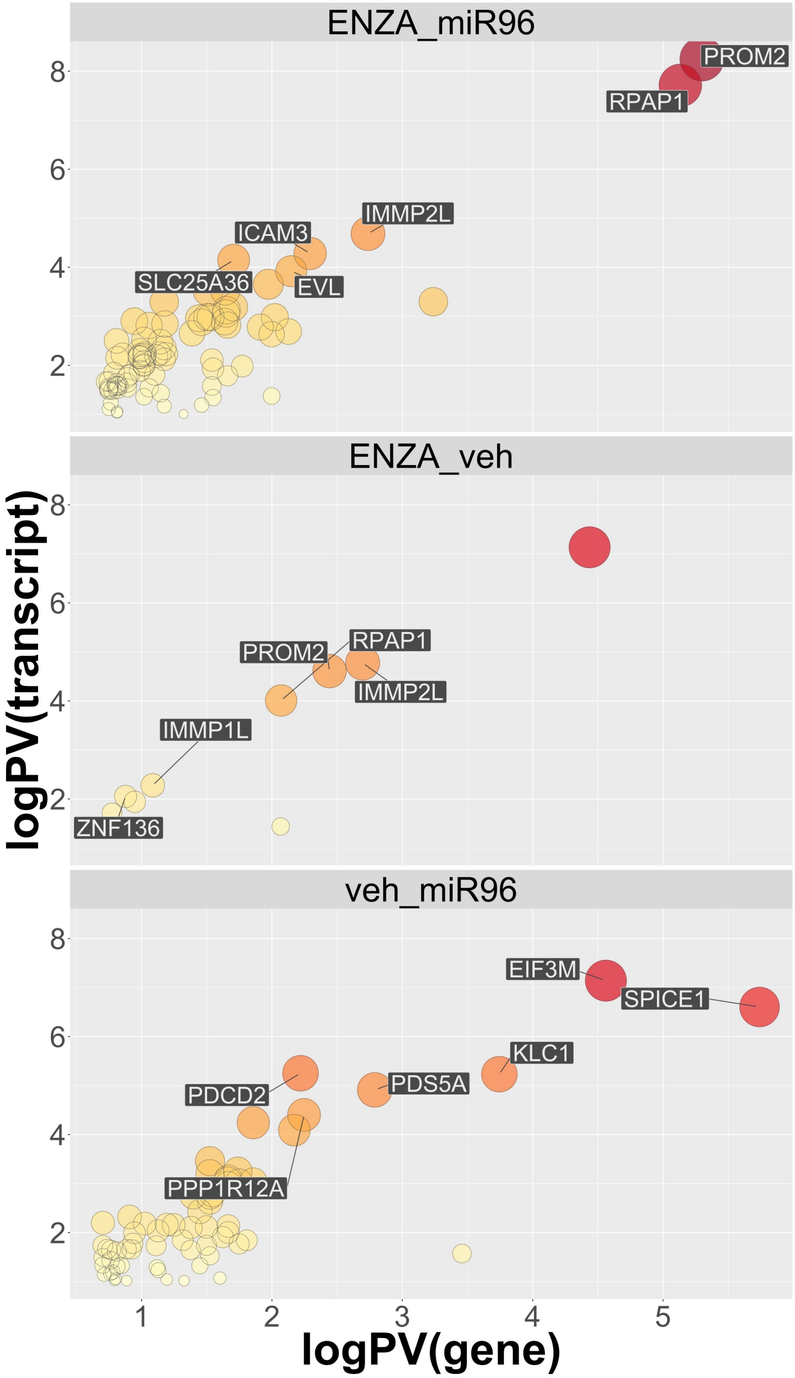


Supplementary Figure 15

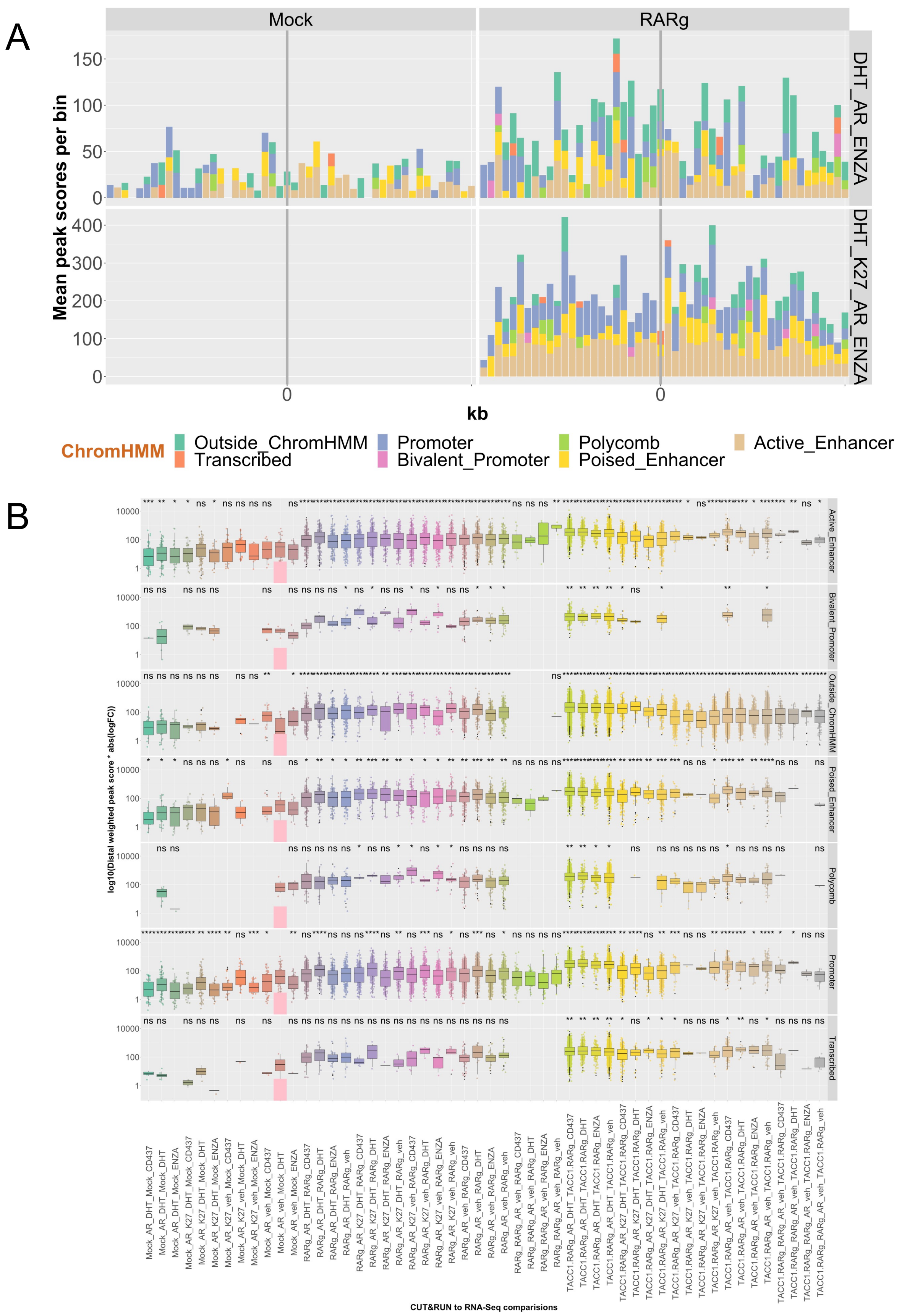


Supplementary Figure 16

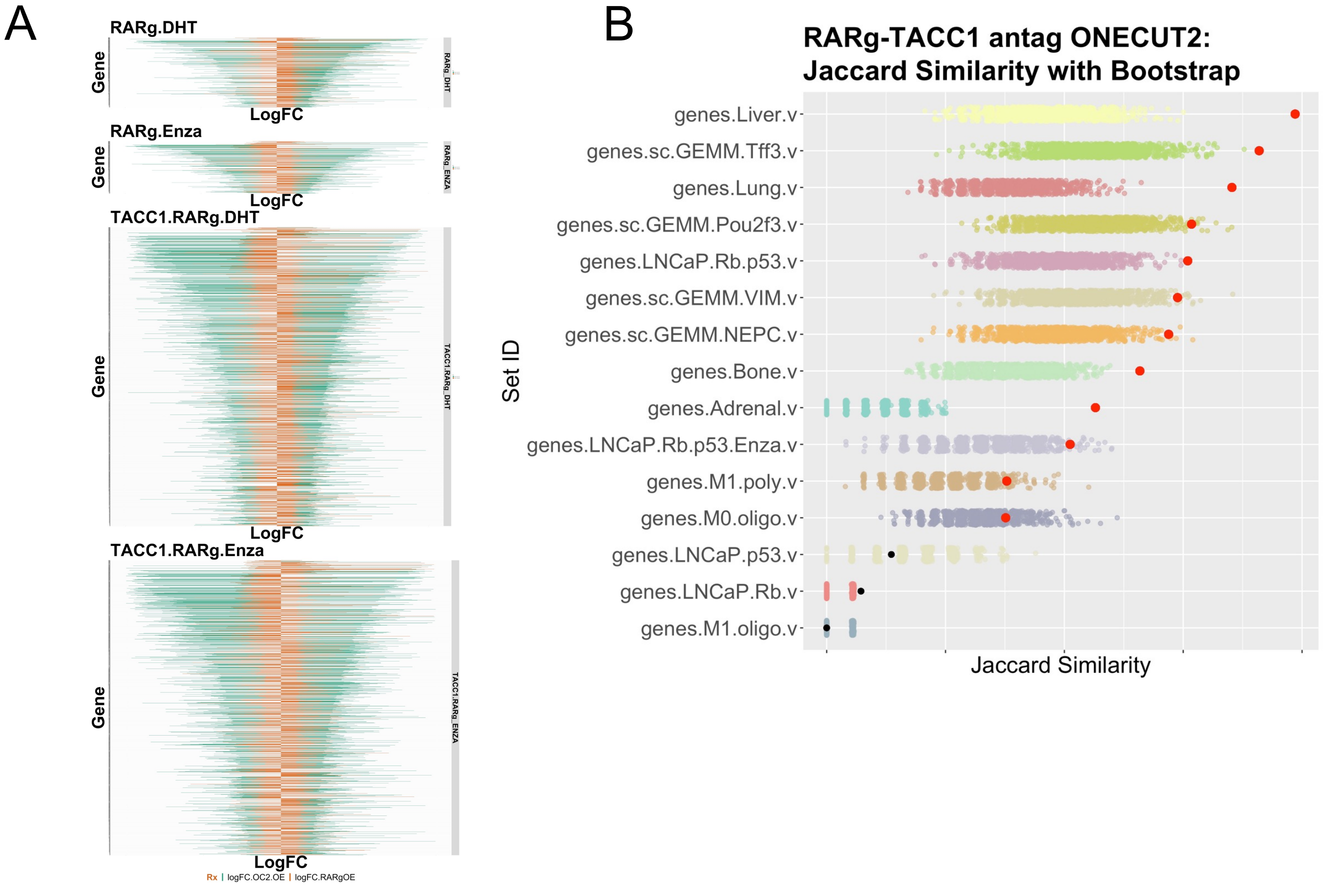


Supplementary Figure 17
