## Supplementary material for "Epigenetic disruption of the RARγ complex impairs its function to bookmark AR enhancer interactions required for enzalutamide sensitivity in prostate cancer": Supp Figures Legends

### Supplementary Figure Legends

SF1: RIME analyses of the RARγ complex. **A.** Co-immunoprecipitation of TACC1 with RARγ. **B.** Cells were treated with CD437 (400 nM, 6h) or vehicle control in quadriplicates and RIME analyses undertaken of RARγ in the indicated cells and significantly different proteins were identified using an `edgeR` workflow. Enriched proteins were classified as to whether they were coregulators (**B**) or splicing factors (**C**).

SF2: Enriched protein networks in RARγ complex. Differentially positively and significantly enriched proteins in the basal (**A**) or CD437-treated (**B**) cells were analyzed by `StringDB` and the logPV(Enrichment) calculated and visualized in each cell background.

SF3: Optimization of antagomir function. HPr1AR and LNCaP cells were treated with the indicated targeting antagomirs and mimics and effectiveness measured by transfecting the indicated cells at the indicated time points, in triplicate. Expression of TWF1 in miR-1 mimic positive control transfected and HMGA2/MYC in let-7c inhibitor positive control transfected cells was detected by quantitative polymerase chain reaction (qPCR) to optimize transfection conditions.

SF4: MiR-96 bound RNA and the overlap with m6A. HPr1AR and LNCaP cells were treated with 3' biotinylated miRIDIAN miR-96 mimic or scramble (50 nM, 24h and 48h) in triplicate and IMPACT-Seq undertaken, and differential enrichment of miR-96 binding sites (MREs) identified with `csaw`, and genes identified that contained one or more MREs. **A.** `UpsetR` plot of the overlap of the MRE regions (upper) or the MRE-containing genes (lower). **B.** In parallel, in triplicate m6A-Seq was undertaken in vehicle treated cells. `csaw` was used to identify differentially enriched regions of m6A sites, and the overlap of m6A-MRE was determined by `ChIPpeakAnno` in HPr1AR (left) or LNCaP (right). **C.** Guitar plots (`Guitar`) of the distribution of MRE for protein-coding and non-coding RNAs. **D.** Guitar plots (`Guitar`) of the distribution of m6A-MRE for protein-coding and non-coding RNAs. **E.** MRE- and m6A-MRE containing genes were classified by class of RNA and the frequency of binding sites calculated in HPr1AR and LNCaP cells.

SF5: MiR-96 regulated RNA and protein targets. HPr1AR and LNCaP cells were treated with 3' biotinylated miRIDIAN miR-96 mimic or scramble (50 nM, 24h and 48h) in triplicate RNA-Seq or LFQ undertaken. Both RNA and protein count matrices were processed with `edgeR` and PCA plots determined. **A. Left**. FASTQ files were QC processed, aligned to hg38 (`Rsubread`). **Right**. Data from capillary-liquid chromatography-nanospray tandem mass spectrometry (LC-MS/MS) was acquired over a 2-hour separation and mean spectral count results generated by two protein inference engines (`Epifany` and `FIDO`). This count matrix was processed in an `edgeR` workflow to identify differentially enriched proteins. **B**. Differentially expressed proteins were identified and filtered for those that contained MRE or m6A-MRE and the difference tested of the absolute fold change between negatively and positively-regulated targets. **C**. MRE-containing DEGs in HPr1AR and LNCaP were analyzed with GSEA using the Hallmarks, Chemical and

Genetic Perturbations and MiRNA libraries, identified 189 significantly enriched terms and the most significant terms (logPV > 3) were plotted, ordered by NES scores.

SF6: RIME analyses of the RARγ complex. Cells were treated with CD437 (400 nM, 6h) or vehicle control in quadriplicates and RIME analyses undertaken of RARγ in the indicated cells and significantly different proteins were identified using an `edgeR` workflow. Enriched proteins were classified as to whether they were contained MRE

SF7: AR, H3K27ac and RARγ cistrome data and motif enrichment. 22Rv1-mock, 22Rv1-RARγ and 22Rv1-RARγ-TACC1 cells treated with DHT (10nM), CD437(400nM) or vehicle for 6h and CUT&RUN undertaken using antibodies to AR, RARγ, H3K27ac or IgG in triplicate. Differential enrichment of regions compared to IgG controls (p.adj < .1) was measured with `csaw`. Motif analyses for AR, RARγ, H3K27ac cistromes was undertaken (`homer`) and motifs were grouped into the classes. The logPval enrichment for nuclear receptors (**Upper**) and FOX (**Lower**) family members are shown.

SF8: Motif enrichment in H3K27ac and RARγ cistromes. 22Rv1-mock, 22Rv1-RARγ and 22Rv1-RARγ-TACC1 cells treated with DHT (10nM), CD437(400nM) or vehicle for 6h and CUT&RUN undertaken using antibodies to AR, RARγ, H3K27ac or IgG in triplicate. Differential enrichment of regions compared to IgG controls (p.adj < .1) was measured with `csaw`. Motif analyses for H3K27ac (**A**) and RARγ (**B**) cistromes was undertaken (`homer`) and motifs were grouped into the indicated classes. For each motif the change in significance of enrichment was calculated across models compared to 22Rv1-mock cells treated with vehicle and per class the mean delta logPval enrichment and mean percentage coverage calculated (indicated by peak width), and the specific motif with the greatest delta per class is indicated.

SF9: Overlap of cistromes within cell models. 22Rv1-mock, 22Rv1-RARγ and 22Rv1-RARγ-TACC1 cells treated with DHT (10nM), CD437(400nM) or vehicle for 6h and CUT&RUN undertaken using antibodies to AR, RARγ, H3K27ac or IgG in triplicate. Differential enrichment of regions compared to IgG controls (p.adj < .1) was measured with `csaw`, and Venn diagrams generated of overlapping regions by a minimum of 1bp (`ChIPpeakAnno`).

SF10: Overlap of cistromes across cell models. 22Rv1-mock, 22Rv1-RARγ and 22Rv1-RARγ-TACC1 cells treated with DHT (10nM), CD437(400nM) or vehicle for 6h and CUT&RUN undertaken using antibodies to AR, RARγ, H3K27ac or IgG in triplicate. Differential enrichment of regions compared to IgG controls (p.adj < .1) was measured with `csaw`. **A.** Circos plots of the top 50% of log10(p.adj) values illustrate connectivity. **B.** The change in significance of overlap of the indicated overlaps compared to 22Rv1-RARγ +/- CD437 (Upper) or 22Rv1-mock +/- DHT (Lower). **C.** Each peak in each cistrome (unique and shared) was overlapped with ChromHMM states and annotated to target genes (within 100kb) and the frequency peak:gene relationships per cistrome per ChromHMM state was calculated.

SF11: Histone cistrome motif enrichment. 22Rv1-mock, 22Rv1-RARγ and 22Rv1-RARγ-TACC1 cells treated with DHT (10nM), CD437(400nM) or vehicle for 6h and CUT&RUN undertaken using antibodies to H3K27ac or IgG in triplicate. Differential enrichment of regions compared to IgG controls (p.adj < .1) was measured with `csaw`. The mean of the log(p.adj) of the significant peaks per cell/treatment condition are shown and significant differences measured compared to 22Rv1-mock cells. Motif analyses for H3K27ac cistromes was undertaken (`homer`) and motifs were grouped into the classes. The logPval enrichment for family members are shown.

SF12: RARγ dependent transcriptomes. **A.** FASTQ files from the indicated cell and treatments were QC processed, aligned to hg38 (`Rsubread`) and the count matrix processed in an `edgeR` workflow and PCA plots determined. **B**. Genes in prostate basal or luminal gene signatured were filtered from the significant DEGs in each cell and treatment condition and the differences in the magnitude of the absolute fold change measured. **C**. GSEA was undertaken and the terms were frequency analyzed and the most common group terms included cell cycle, nuclear receptors and epigenetics. The most significant NR-related terms are shown (p.adj > 0. 1) in ranked organization.

SF13: The RARγ-dependent alternatively spliced transcriptome in response to DHT, Enza or CD437 in 22Rv1 cell variants**.** 22Rv1 cell variants were treated with DHT (10nM), CD437 (400nM) or ENZA (10μM) for 24h in triplicate and RNA-Seq undertaken. FASTQ files were QC processed, aligned to hg38 (`Rsubread`), and differentially expressed transcripts (DETs) (logPV > 1) identified with `DRIM-Seq`. **A.** The logPV(gene) plotted against logPV(transcript). **B.** Western immunoblot measurements of AR and ARv7 levels following the indicated treatments or vehicle control.

SF14: The combination of Enza plus miR-96 antagomir in 22Rv1 cells. **A**. 22Rv1 cells were treated Enza (10nM), miR-96 antagomir (50nM) or the combination for 96h and proliferation measured. **B**. As above but for 24h in triplicate and RNA-Seq undertaken. FASTQ files were QC processed, aligned to hg38 (`Rsubread`) and PCA undertaken.

SF15: Enza plus miR-96 antagomir in 22Rv1 cells and differentially expressed transcripts (DETs) (logPV > 1) identified with `DRIM-Seq`. The logPV(gene) plotted against logPV(transcript).

SF16: Cistrome and transcriptome integration. 22Rv1-mock, 22Rv1-RARγ and 22Rv1-RARγ-TACC1 cells treated with DHT (10nM), CD437(400nM) or vehicle for 6h and CUT&RUN undertaken using antibodies to AR, RARγ, H3K27ac or IgG in triplicate, and treated in the same way for 24h in triplicate and RNA-Seq undertaken. In both cases FASTQ files were QC processed, aligned to hg38 (`Rsubread`), and differentially expressed peaks compared to IgG controls (p.adj < .1) identified by `csaw`, and differentially expressed genes (logPV > 1 & absFC > .37) identified with `edgeR`. Differentially enriched peaks were overlapped with ChromHMM regions (ChromHMM) identified in LNCaP using `bedtools`. **A**. The frequency of ChromHMM peak:gene relationships per 10K bin around target genes. The column heading is the cell background and the row is the cistrome-transcriptome relationship. DHT_AR_Enza is the DHT dependent AR cistrome annotated to Enza-regulated genes, and AR_K27 is the shared sites of AR and H3K27ac binding annotated to genes. B.

SF17. RARγ and ONECUT2. **A.** Antagonistic gene expression between either DHT or Enza regulated genes in 22Rv1-RARγ cells and 22Rv1- RARγ -TACC1 cells compared to gene regulation by over-expression of ONECUT2. Transcriptomic data in 22Rv1- RARγ cells to 22Rv1 cells with ONECUT2 over-expression, and defined gene expression as cooperative, if for a given gene the direction of change was the same, and antagonistic if the direction of change was opposite. The height of each panel is proportional to the number of events. **B.** The 704 RARγ-ONECUT2 antagonized genes were extracted from 22Rv1 cells to also generate a ranked list. The similarity of these rank genes was compared to the ranked gene lists from the indicated experiments and similarity measured by Jaccard similarity measurements, and the significance determined by bootstrapping (n=1000). The symbols in red are significant similarity.
