## Supplementary material for "Epigenetic disruption of the RARγ complex impairs its function to bookmark AR enhancer interactions required for enzalutamide sensitivity in prostate cancer": Supp Tables

Supplementary tables
Supplementary figure legends

| Cell.Rx | Direct | Coreg | Overlap | logPV.hyper | MostSignificant |
|---|---|---|---|---|---|
| 22RV1.RARg.TACC1_veh | P | Mixed | 272 | 290.68 | HCFC1 |
| LNCaP_veh | P | CoA | 216 | 256.71 | NSUN2 |
| 22RV1.RARg_CD437 | P | TF | 384 | 247.90 | RARG |
| 22RV1.RARg_CD437 | P | CoR | 232 | 234.59 | PA2G4 |
| 22RV1.RARg_CD437 | P | Mixed | 200 | 197.66 | HCFC1 |
| 22RV1.RARg_veh | P | CoR | 160 | 168.17 | DNMT1 |
| LNCaP_CD437 | P | CoA | 128 | 130.32 | NSUN2 |
| 22RV1.RARg_veh | P | TF | 192 | 92.86 | RARG |
| 22RV1.RARg_veh | P | Mixed | 104 | 89.14 | HCFC1 |
| LNCaP_veh | P | CoR | 96 | 76.21 | NCOR2 |
| LNCaP_CD437 | P | TF | 128 | 63.81 | RARG |
| LNCaP_veh | P | TF | 152 | 60.50 | RARG |
| LNCaP_veh | P | Mixed | 72 | 51.19 | RBM15 |
| LNCaP_CD437 | P | CoR | 48 | 29.67 | PA2G4 |
| LNCaP_CD437 | P | Mixed | 40 | 23.89 | RBM15 |

**Supplementary Table 1A**: Enrichment of coregulators in the RARγ complex. LNCaP, 22Rv1, 22Rv1-RARγ and 22Rv1-RARγ-TACC1 cells in quadruplicate were treated with vehicle or CD437 (400 nM, 6 h), and RIME undertaken to identify significantly and positively enriched proteins in the complex, which were then annotated as to whether they were either a coactivator (CoA), corepressor (CoR), mixed function coregulator (mixed) or a transcription factor (TF). A hypergeometric test was undertaken to examine if the overlaps for each group of proteins were significantly enriched in the RARγ complex, and the most positively enriched protein is indicated.

| Cell.Rx | Direct | MostSignificant |
|---|---|---|
| LNCaP.CD437 vs veh | P | RXRA |
| LNCaP.CD437 vs veh | P | ESRRA |
| LNCaP.CD437 vs veh | P | HNF4G |
| LNCaP.CD437 vs veh | P | NR2F2 |
| LNCaP.CD437 vs veh | P | NR2C1 |
| LNCaP.CD437 vs veh | P | NR2C2 |
| LNCaP.CD437 vs veh | P | AR |
| LNCaP.CD437 vs veh | P | RXRB |
| RV1 vs LNCaP RARg.veh | P | RXRB |
| RV1 vs LNCaP RARg.veh | P | RARG |
| RV1 vs LNCaP RARg.veh | P | RXRA |
| RV1 vs LNCaP RARg.veh | P | NR2C1 |
| RV1 vs LNCaP RARg.veh | P | NR2C2 |
| RV1 vs LNCaP RARg.veh | P | AR |

**Supplementary Table 1B**: Change in enrichment in nuclear receptor targets in the RARγ complex. LNCaP, 22Rv1, 22Rv1-RARγ and 22Rv1-RARγ-TACC1 cells in quadruplicate were treated with vehicle or CD437 (400 nM, 6 h) and RIME undertaken. In each case the change in enrichment was then calculated between the indicated conditions to identify differentially enriched proteins, both negatively and positively enriched in the complex, which were then annotated as to whether Nuclear Receptors. A hypergeometric test was undertaken to examine if the overlaps for each group of proteins were significantly enriched in these altered enrichments in the RARγ complex, and the most significantly altered protein is indicated.

| Cell.Rx | Overlap | logPV.hyper | Threshold.hyper | MostSignificant |
|---|---|---|---|---|
| 22RV1.RARg.TACC1_veh | 44 | 4.50 | Significant | SMCHD1 |
| 22RV1.RARg.TACC1_CD437 | 45 | 4.16 | Significant | TRRAP |
| 22RV1.RARg_veh | 20 | 2.07 | Significant | SMCHD1 |
| LNCaP_CD437 | 14 | 1.85 | Significant | PLEC |
| 22RV1.RARg_CD437 | 28 | 1.81 | Significant | SMCHD1 |

**Supplementary Table 1C**: Enrichment of coregulators in the RAR$\gamma$ complex. LNCaP, 22Rv1, 22Rv1-RAR$\gamma$ and 22Rv1-RAR$\gamma$-TACC1 cells in quadruplicate were treated with vehicle or CD437 (400 nM, 6 h), and RIME undertaken to identify significantly and positively enriched proteins in the complex, which were then annotated as to whether they were in the list of mutated chromatin remodelers; so-called long-tail mutants. A hypergeometric test was undertaken to examine if the overlaps for each group of proteins were significantly enriched in the RAR$\gamma$ complex, and the most positively enriched protein is indicated.

| Cell | Exper | Time | n |
|---|---|---|---|
| HPr1AR | IMPACT | 24h | 472 |
| HPr1AR | IMPACT | 48h | 7494 |
| HPr1AR | m6A.IMPACT | 24h | 241 |
| HPr1AR | m6A.IMPACT | 48h | 7258 |
| LNCaP | IMPACT | 24h | 3224 |
| LNCaP | IMPACT | 48h | 333 |
| LNCaP | m6A.IMPACT | 24h | 2888 |
| LNCaP | m6A.IMPACT | 48h | 263 |

**Supplementary Table 2A**. Frequency of miR-96 and m6A-miR-96 binding sites by time and cell background. MiR-96 binding sites were identified by IMPACT-Seq in HPr1AR and LNCaP cells by transfecting cells with a biotinylated miR-96 mimic (50 nM) and after 24 and 48 h cells were lysed and miR-96 interacting mRNA isolated by streptavidin beads, eluted, sequenced, and aligned to the transcriptome and discrete interacting sites (IMPACT) identified by `csaw` (n). In parallel, m6A-Seq was undertaken in the same cells and the overlap of significantly shared sites with m6A at a miR-96 binding sites (m6A.IMPACT) with `bedtools` and the frequency also enumerated (n).

| Cell | Exper | Time | n |
|---|---|---|---|
| HPr1AR | IMPACT.genes | 24h | 423 |
| HPr1AR | IMPACT.genes | 48h | 2859 |
| HPr1AR | m6A.IMPACT.genes | 24h | 227 |
| HPr1AR | m6A.IMPACT.genes | 48h | 2859 |
| LNCaP | IMPACT.genes | 24h | 2381 |
| LNCaP | IMPACT.genes | 48h | 245 |
| LNCaP | m6A.IMPACT.genes | 24h | 2113 |
| LNCaP | m6A.IMPACT.genes | 48h | 188 |

**Supplementary Table 2B**. Frequency of miR-96 and m6A-miR-96 targeted genes by time and cell background. The frequency of genes containing miR-96 binding sites (IMPACT.genes) and those shared with m6A (m6A.IMPACT.genes) were calculated at the indicated time points in each cell backgroun (n).

IMPACT-Seq: Coregulators and TFs

| | HPr1AR*24h* IMP N = 472[1] | HPr1AR*24h* m6A.IMP N = 241[1] | LNCaP*48h* IMP N = 333[1] | LNCaP*48h* m6A.IMP N = 263[1] | **p-value**[2] |
|---|---|---|---|---|---|
| **CoA** | 32 (6.8%) | 14 (5.8%) | 15 (4.5%) | 10 (3.8%) | 0.3 |
| **CoR** | 16 (3.4%) | 8 (3.3%) | 11 (3.3%) | 13 (4.9%) | 0.7 |
| **Mixed** | 17 (3.6%) | 9 (3.7%) | 7 (2.1%) | 4 (1.5%) | 0.3 |
| **TF** | 41 (8.7%) | 22 (9.1%) | 16 (4.8%) | 5 (1.9%) | <0.001 |
| **NR** | 2 (0.4%) | 0 (0%) | 0 (0%) | 0 (0%) | 0.5 |

[1]n (%)

[2]Fisher's Exact Test for Count Data with simulated p-value (based on 2000 replicates)

IMPACT-Seq: Coregulators and TFs

| | HPr1AR24*h* IMP.genes, N = 423[1] | HPr1AR*24h*m6A. IMP.genes, N = 227[1] | **LNCaP*48h* IMP.genes**, N = 245[1] | **LNCaP*48h*m6A. IMP.genes**, N = 188[1] | **p-value**[2] |
|---|---|---|---|---|---|
| **CoA** | 27 (6.4%) | 13 (5.7%) | 11 (4.5%) | 6 (3.2%) | 0.4 |
| **CoR** | 13 (3.1%) | 8 (3.5%) | 9 (3.7%) | 8 (4.3%) | 0.9 |
| **Mixed** | 15 (3.5%) | 8 (3.5%) | 6 (2.4%) | 4 (2.1%) | 0.8 |
| **TF** | 40 (9.5%) | 20 (8.8%) | 16 (6.5%) | 5 (2.7%) | 0.014 |
| **NR** | 2 (0.5%) | 0 (0%) | 0 (0%) | 0 (0%) | 0.7 |

[1]n (%)

[2]Fisher's Exact Test for Count Data with simulated p-value (based on 2000 replicates)

**Supplementary Table 2C.** Enrichment of different classes of protein coding genes in miR-96 targetomes. miR-96 targeted protein coding genes were categorized if they were coactivators (CoAs), corepressors (CoRs), mixed function coregulators (Mixed) or transcription and mRNA stabilization factors (TFs) and the frequency of the number of sites (**Top**) or frequency of targeted genes (**Bottom**) was determined and the significant differences measured with a Fisher's exact test.

| Cell | coreg | NR | Direct | HPr1AR. MRE | HPr1AR. m6A.MRE | LNCaP. m6A.MRE | LNCaP. MRE | Most Significant |
|---|---|---|---|---|---|---|---|---|
| HPr1AR | CoA | - | N | 3 | 3 | 0 | 0 | EEF2 |
| HPr1AR | CoA | - | P | 7 | 6 | 0 | 0 | GRHL3 |
| HPr1AR | CoR | - | N | 2 | 2 | 0 | 0 | PDCD4 |
| HPr1AR | CoR | - | P | 1 | 1 | 0 | 0 | SFMBT1 |
| HPr1AR | Mixed | - | N | 2 | 0 | 0 | 0 | MAGED1 |
| HPr1AR | Mixed | - | N | 0 | 1 | 0 | 0 | BRPF3 |
| HPr1AR | Mixed | - | P | 3 | 6 | 0 | 0 | RBPJ |
| HPr1AR | other | - | N | 0 | 84 | 0 | 0 | SLC39A1 |
| HPr1AR | other | - | N | 112 | 0 | 0 | 0 | MSN |
| HPr1AR | other | - | P | 0 | 64 | 0 | 0 | CDKN1A |
| HPr1AR | other | - | P | 105 | 0 | 0 | 0 | ALPK2 |
| HPr1AR | TF | - | N | 6 | 6 | 0 | 0 | ZFP36L1 |
| HPr1AR | TF | - | P | 14 | 14 | 0 | 0 | E2F7 |
| LNCaP | CoA | - | N | 0 | 0 | 11 | 14 | TACC1 |
| LNCaP | CoA | - | P | 0 | 0 | 0 | 10 | MYO6 |
| LNCaP | CoA | - | P | 0 | 0 | 12 | 0 | PPP2R5B |
| LNCaP | CoR | - | N | 0 | 0 | 7 | 0 | PDCD4 |
| LNCaP | CoR | - | N | 0 | 0 | 0 | 6 | CBX6 |
| LNCaP | CoR | - | P | 0 | 0 | 6 | 4 | TAF9B |
| LNCaP | Mixed | - | N | 0 | 0 | 5 | 7 | TBL1X |
| LNCaP | Mixed | - | P | 0 | 0 | 4 | 7 | RBPJ |
| LNCaP | other | - | N | 0 | 0 | 163 | 248 | CD164 |
| LNCaP | other | - | P | 0 | 0 | 104 | 154 | CBLN2 |
| LNCaP | TF | - | N | 0 | 0 | 13 | 20 | CREB3L2 |
| LNCaP | TF | Yes | N | 0 | 0 | 2 | 3 | RARG |
| LNCaP | TF | - | P | 0 | 0 | 11 | 13 | TRIB1 |

**Supplementary Table 3A**: Regulated mRNA targets of miR-96 mimics in HPr1AR and LNCaP cells. Cells were transfected with a miR-96 mimic or scrambled control (50nM, 48 h) and RNA-Seq undertaken to identify significantly differentially expressed genes (DEGs), which were then divided in negatively (N) and positively (P) regulated genes and annotated as to whether they were either a coactivator (CoA), corepressor (CoR), mixed function coregulator (mixed) or a transcription factor (TF), and if a TF, whether a nuclear receptor (NR). Genes in each of these groups were then annotated as to whether they were bound by miR-96 (MRE), or at a MRE of significant m6A enrichment (m6A-MRE). Finally, for each grouping the gene with the greatest significance was identified.

| Cell | coreg | NR | Direct | HPr1AR. MRE.24 | HPr1AR. MRE.48 | LNCaP. MRE.24 | LNCaP. MRE.48 | Most Significant |
|---|---|---|---|---|---|---|---|---|
| HPr1AR | CoA | - | N | 10 | 16 | 0 | 0 | EEF1A1 |
| HPr1AR | CoA | - | P | 14 | 24 | 0 | 0 | ATAD2 |
| HPr1AR | CoR | - | N | 5 | 9 | 0 | 0 | HSPA8 |
| HPr1AR | CoR | - | P | 0 | 22 | 0 | 0 | NELFCD |
| HPr1AR | CoR | - | P | 8 | 0 | 0 | 0 | UBE2I |
| HPr1AR | Mixed | - | N | 0 | 9 | 0 | 0 | HNRNPC |
| HPr1AR | Mixed | - | N | 7 | 0 | 0 | 0 | PHF6 |
| HPr1AR | Mixed | - | P | 4 | 16 | 0 | 0 | NIPBL |
| HPr1AR | other | - | N | 113 | 0 | 0 | 0 | PKM |
| HPr1AR | other | - | N | 0 | 169 | 0 | 0 | CFL1 |
| HPr1AR | other | - | P | 0 | 255 | 0 | 0 | VPS37B |
| HPr1AR | other | - | P | 172 | 0 | 0 | 0 | UBE2D3 |
| HPr1AR | TF | - | N | 8 | 11 | 0 | 0 | POLR2A |
| HPr1AR | TF | - | P | 9 | 15 | 0 | 0 | SUPT5H |
| HPr1AR | TF | NR | P | 1 | 1 | 0 | 0 | NR3C1 |
| LNCaP | CoA | - | N | 0 | 0 | 9 | 0 | EEF1A1 |
| LNCaP | CoA | - | N | 0 | 0 | 0 | 1 | MORF4L2 |
| LNCaP | CoA | - | P | 0 | 0 | 8 | 0 | HAX1 |
| LNCaP | CoA | - | P | 0 | 0 | 0 | 2 | BAZ2A |
| LNCaP | CoR | - | N | 0 | 0 | 2 | 0 | HSPA8 |
| LNCaP | CoR | - | P | 0 | 0 | 3 | 0 | MBD3 |
| LNCaP | Mixed | - | N | 0 | 0 | 1 | 0 | MECP2 |
| LNCaP | Mixed | - | P | 0 | 0 | 3 | 0 | GLS |
| LNCaP | Mixed | - | P | 0 | 0 | 0 | 1 | CHD8 |
| LNCaP | other | - | N | 0 | 0 | 82 | 0 | TUBA1B |
| LNCaP | other | - | N | 0 | 0 | 0 | 13 | CFL1 |
| LNCaP | other | - | P | 0 | 0 | 89 | 0 | PTCD3 |
| LNCaP | other | - | P | 0 | 0 | 0 | 6 | RPL8 |
| LNCaP | TF | - | N | 0 | 0 | 4 | 0 | NUCKS1 |
| LNCaP | TF | - | N | 0 | 0 | 0 | 1 | ENO1 |
| LNCaP | TF | - | P | 0 | 0 | 3 | 0 | CUX1 |

**Supplementary Table 3B**: Regulated protein targets of miR-96 mimics in HPr1AR and LNCaP cells. were transfected with a miR-96 mimic or scrambled control (50 nM, 48 h) and label free quantitative proteomics undertaken to identify significantly differentially expressed proteins (DEPs), which were then divided in negatively (N) and positively (P) regulated targets and annotated as to whether they were either a coactivator (CoA), corepressor (CoR), mixed function coregulator (mixed) or a transcription factor (TF), and if a TF, whether a nuclear receptor (NR). Genes in each of these groups were then annotated as to whether they were bound by miR-96 (MRE), or at a MRE of significant m6A enrichment (m6A-MRE). Finally, for each grouping the target with the greatest significance was identified.

| Cell.Rx | Direct | IMPACT | Overlap | logPV.hyper | MostSignificant |
|---|---|---|---|---|---|
| 22RV1.RARg.TACC1_veh | P | HPR.m6A.IMP | 250 | 50.07 | PRRC2C |
| 22RV1.RARg_CD437 | P | HPR.m6A.IMP | 212 | 48.68 | PRRC2C |
| 22RV1.RARg.TACC1_CD437 | P | HPR.m6A.IMP | 258 | 48.52 | DNMT1 |
| 22RV1.RARg_CD437 | P | HPR.IMP | 208 | 41.51 | PRRC2C |
| 22RV1.RARg.TACC1_CD437 | P | HPR.IMP | 255 | 41.48 | DNMT1 |
| 22RV1.RARg.TACC1_veh | P | HPR.IMP | 244 | 41.39 | PRRC2C |
| 22RV1.RARg_veh | P | HPR.m6A.IMP | 141 | 37.22 | SSRP1 |
| 22RV1.RARg_veh | P | HPR.IMP | 141 | 33.97 | SSRP1 |
| LNCaP_CD437 | P | HPR.IMP | 98 | 26.70 | NSUN2 |
| LNCaP_veh | P | HPR.m6A.IMP | 115 | 24.86 | NSUN2 |
| LNCaP_veh | P | HPR.IMP | 114 | 21.81 | NSUN2 |
| LNCaP_CD437 | P | HPR.m6A.IMP | 87 | 21.18 | NSUN2 |
| 22RV1.RARg_CD437 | P | LNC.IMP | 136 | 16.58 | RARG |
| 22RV1.RARg.TACC1_veh | P | LNC.m6A.IMP | 143 | 14.93 | RARG |
| 22RV1.RARg.TACC1_CD437 | P | LNC.m6A.IMP | 146 | 13.59 | RARG |
| 22RV1.RARg_CD437 | P | LNC.m6A.IMP | 117 | 13.53 | RARG |
| LNCaP_CD437 | P | LNC.IMP | 67 | 13.11 | RARG |
| 22RV1.RARg.TACC1_CD437 | P | LNC.IMP | 159 | 13.01 | RARG |
| 22RV1.RARg_veh | P | LNC.m6A.IMP | 81 | 12.44 | RARG |
| 22RV1.RARg.TACC1_veh | P | LNC.IMP | 145 | 10.90 | RARG |
| 22RV1.RARg_veh | P | LNC.IMP | 84 | 10.62 | RARG |
| LNCaP_veh | P | LNC.m6A.IMP | 72 | 10.50 | RARG |
| LNCaP_CD437 | P | LNC.m6A.IMP | 50 | 7.15 | RARG |
| LNCaP_veh | P | LNC.IMP | 69 | 6.78 | RARG |

**Supplementary Table 3C**: Enrichment of miR-96 MRE targets in the RARγ complex**.** LNCaP, 22Rv1, 22Rv1-RARγ and 22Rv1-RARγ-TACC1 cells in quadruplicate were treated with vehicle or CD437 (400 nM, 6 h) and RIME undertaken to identify significantly and positively enriched proteins in the complex, which were then annotated as to whether they enriched for proteins encoded by MRE-containing genes, and the RARγ complexes are ranked, and the most positively enriched protein is indicated.

| Cell.Rx | Direct | coreg | Number | MostSignificant |
|---|---|---|---|---|
| LNCaP.CD437 vs veh | N | Coactivator | 9 | PTBP1 |
| LNCaP.CD437 vs veh | N | Corepressor | 6 | NCOR2 |
| LNCaP.CD437 vs veh | N | Mixed | 11 | HNRNPA1 |
| LNCaP.CD437 vs veh | P | Coactivator | 49 | KMT2D |
| LNCaP.CD437 vs veh | P | Corepressor | 21 | RCOR3 |
| LNCaP.CD437 vs veh | P | Mixed | 26 | SRC |
| LNCaP.CD437 vs veh | P | TF | 51 | RXRA |
| RV1.RARg.CD437 vs veh | P | Corepressor | 1 | CBX4 |
| RV1.RARg.CD437 vs veh | P | TF | 2 | SUPT5H |
| RV1 vs LNCaP RARg.veh | N | Coactivator | 6 | MYO6 |
| RV1 vs LNCaP RARg.veh | N | Corepressor | 3 | NCOR2 |
| RV1 vs LNCaP RARg.veh | N | Mixed | 1 | HNRNPA0 |
| RV1 vs LNCaP RARg.veh | N | TF | 2 | POLR1C |
| RV1 vs LNCaP RARg.veh | P | Coactivator | 37 | PHIP |
| RV1 vs LNCaP RARg.veh | P | Corepressor | 16 | KHDRBS1 |
| RV1 vs LNCaP RARg.veh | P | Mixed | 23 | HNRNPH3 |
| RV1 vs LNCaP RARg.veh | P | TF | 31 | ZBTB10 |
| RV1.RARg.TACC1.CD437 vs veh | N | Coactivator | 2 | NSD1 |
| RV1.RARg.TACC1.CD437 vs veh | N | Corepressor | 2 | NCOR2 |

**Supplementary Table 3D**: Change in enrichment of coregulators in the RARγ complex**.** LNCaP, 22Rv1, 22Rv1-RARγ and 22Rv1-RARγ-TACC1 cells in quadruplicate were treated with vehicle or CD437 (400 nM, 6 h) and RIME undertaken. In each case the change in enrichment was then calculated between the indicated conditions to identify differentially enriched proteins, both negatively and positively enriched in the complex, which were then annotated as to whether they were either a coactivator (CoA), corepressor (CoR), mixed function coregulator (mixed) or a transcription factor (TF). A hypergeometric test was undertaken to examine if the overlaps for each group of proteins were significantly enriched in these altered enrichments in the RARγ complex, and the most significantly altered protein is indicated.

| Cell.Rx | Coreg | IMPACT | Number | MostSignificant |
|---|---|---|---|---|
| 22RV1.RARg.TACC1_CD437 | CoA | HPR.IMP | 46 | TRRAP |
| 22RV1.RARg.TACC1_veh | CoA | HPR.IMP | 46 | NSD1 |
| 22RV1.RARg.TACC1_veh | CoA | HPR.m6A.IMP | 46 | NSD1 |
| 22RV1.RARg.TACC1_CD437 | CoA | HPR.m6A.IMP | 45 | TRRAP |
| 22RV1.RARg_CD437 | CoA | HPR.IMP | 36 | NSUN2 |
| 22RV1.RARg_CD437 | CoA | HPR.m6A.IMP | 36 | NSUN2 |
| 22RV1.RARg.TACC1_veh | TF | HPR.m6A.IMP | 35 | CSDE1 |
| 22RV1.RARg.TACC1_CD437 | TF | HPR.m6A.IMP | 32 | SUPT5H |
| 22RV1.RARg.TACC1_veh | TF | HPR.IMP | 30 | CSDE1 |
| 22RV1.RARg.TACC1_CD437 | TF | HPR.IMP | 28 | SUPT5H |
| 22RV1.RARg.TACC1_veh | CoA | LNC.m6A.IMP | 28 | NSD1 |
| 22RV1.RARg_veh | CoA | HPR.m6A.IMP | 26 | NSUN2 |
| 22RV1.RARg.TACC1_CD437 | CoA | LNC.m6A.IMP | 26 | TRRAP |
| 22RV1.RARg.TACC1_CD437 | CoA | LNC.IMP | 25 | TRRAP |
| 22RV1.RARg_veh | CoA | HPR.IMP | 24 | NSUN2 |
| 22RV1.RARg.TACC1_veh | CoA | LNC.IMP | 24 | NSD1 |
| 22RV1.RARg.TACC1_CD437 | Mixed | HPR.m6A.IMP | 23 | HCFC1 |
| 22RV1.RARg_CD437 | TF | HPR.m6A.IMP | 22 | DDB1 |
| 22RV1.RARg_CD437 | CoA | LNC.IMP | 21 | SUPT16H |
| 22RV1.RARg.TACC1_CD437 | Mixed | HPR.IMP | 21 | HCFC1 |
| 22RV1.RARg.TACC1_veh | CoR | HPR.m6A.IMP | 21 | DNMT1 |
| 22RV1.RARg.TACC1_veh | Mixed | HPR.m6A.IMP | 21 | HCFC1 |
| 22RV1.RARg_CD437 | CoA | LNC.m6A.IMP | 20 | SUPT16H |
| 22RV1.RARg_CD437 | TF | HPR.IMP | 20 | DDB1 |
| 22RV1.RARg.TACC1_veh | Mixed | HPR.IMP | 20 | HCFC1 |
| 22RV1.RARg.TACC1_CD437 | CoR | HPR.m6A.IMP | 19 | DNMT1 |
| 22RV1.RARg.TACC1_veh | CoR | HPR.IMP | 19 | DNMT1 |
| LNCaP_veh | CoA | HPR.m6A.IMP | 19 | NSUN2 |
| LNCaP_veh | CoA | HPR.IMP | 18 | NSUN2 |
| 22RV1.RARg.TACC1_CD437 | CoR | HPR.IMP | 17 | DNMT1 |
| 22RV1.RARg_CD437 | CoR | HPR.m6A.IMP | 16 | XRCC5 |
| 22RV1.RARg_veh | CoA | LNC.m6A.IMP | 16 | SUPT16H |
| 22RV1.RARg.TACC1_veh | TF | LNC.IMP | 16 | RARG |
| 22RV1.RARg_CD437 | Mixed | HPR.m6A.IMP | 15 | HCFC1 |
| 22RV1.RARg.TACC1_CD437 | CoR | LNC.IMP | 15 | XRCC5 |
| 22RV1.RARg.TACC1_CD437 | Mixed | LNC.IMP | 15 | HCFC1 |
| 22RV1.RARg.TACC1_CD437 | TF | LNC.IMP | 15 | RARG |
| 22RV1.RARg.TACC1_veh | TF | LNC.m6A.IMP | 15 | RARG |
| 22RV1.RARg_CD437 | CoR | HPR.IMP | 14 | XRCC5 |
| 22RV1.RARg_veh | CoA | LNC.IMP | 14 | SUPT16H |
| 22RV1.RARg.TACC1_CD437 | CoR | LNC.m6A.IMP | 14 | XRCC5 |
| 22RV1.RARg.TACC1_veh | CoR | LNC.m6A.IMP | 14 | NACC1 |
| LNCaP_CD437 | CoA | HPR.m6A.IMP | 14 | NSUN2 |
| 22RV1.RARg_CD437 | Mixed | HPR.IMP | 13 | HCFC1 |
| 22RV1.RARg.TACC1_CD437 | Mixed | LNC.m6A.IMP | 13 | HCFC1 |
| 22RV1.RARg.TACC1_CD437 | TF | LNC.m6A.IMP | 13 | RARG |
| 22RV1.RARg.TACC1_veh | CoR | LNC.IMP | 13 | NACC1 |
| 22RV1.RARg.TACC1_veh | Mixed | LNC.IMP | 13 | HCFC1 |
| LNCaP_CD437 | CoA | HPR.IMP | 13 | NSUN2 |
| 22RV1.RARg_CD437 | TF | LNC.IMP | 12 | RARG |
| 22RV1.RARg_veh | CoR | HPR.m6A.IMP | 12 | DNMT1 |

| | | | | |
|---|---|---|---|---|
| 22RV1.RARg.TACC1_veh | Mixed | LNC.m6A.IMP | 12 | HCFC1 |
| LNCaP_veh | CoA | LNC.IMP | 12 | SUPT16H |
| 22RV1.RARg_CD437 | CoR | LNC.IMP | 11 | XRCC5 |
| 22RV1.RARg_CD437 | CoR | LNC.m6A.IMP | 11 | XRCC5 |
| 22RV1.RARg_CD437 | Mixed | LNC.IMP | 11 | HCFC1 |
| 22RV1.RARg_veh | CoR | HPR.IMP | 11 | DNMT1 |
| 22RV1.RARg_veh | TF | HPR.m6A.IMP | 11 | DDB1 |
| LNCaP_veh | CoA | LNC.m6A.IMP | 11 | SUPT16H |

**Supplementary Table 3D– Change in enrichment of coregulators that contain a miR-96 MRE in the RARγ complex.** LNCaP, 22Rv1, 22Rv1-RARγ and 22Rv1-RARγ-TACC1 cells in quadruplicate were treated with vehicle or CD437 (400 nM, 6 h) and RIME undertaken. In each case enriched proteins, compared to IgG, were annotated as to whether they were either a coactivator (CoA), corepressor (CoR), mixed function coregulator (mixed) or a transcription factor (TF) and contained a miR-96 binding site by IMPACT-Seq (IMP) that also contained a m6A modified region (m6A.IMP) in the indicated conditions. The number of proteins that meet each criterion is indicated and the most positively enriched protein is indicated.

| Cell | Target | Rx | n | logPval |
|---|---|---|---|---|
| Mock | AR | veh | 768 | 260.81 |
| Mock | AR | DHT | 1368 | 64.36 |
| RARg | AR | veh | 9649 | 291.65 |
| RARg | AR | DHT | 5554 | 330.78 |
| TACC1.RARg | AR | veh | 82090 | 375.79 |
| TACC1.RARg | AR | DHT | 119149 | 387.00 |
| RARg | RARγ | veh | 411 | 137.44 |
| RARg | RARγ | CD437 | 284 | 402.08 |
| TACC1.RARg | RARγ | veh | 6278 | 455.64 |
| TACC1.RARg | RARγ | CD437 | 378 | 332.87 |
| Mock | K27 | veh | 29755 | 628.63 |
| Mock | K27 | DHT | 44594 | 612.61 |
| RARg | K27 | veh | 26554 | 647.89 |
| RARg | K27 | DHT | 25446 | 658.48 |
| RARg | K27 | CD437 | 25838 | 652.64 |
| TACC1.RARg | K27 | veh | 43758 | 675.59 |
| TACC1.RARg | K27 | DHT | 29118 | 657.20 |
| TACC1.RARg | K27 | CD437 | 32828 | 621.60 |

**Supplementary Table 4A. Summary of significantly enriched binding sites for AR, RARγ and H3K27ac**. 22Rv1, 22Rv1-RARγ and 22Rv1-RARγ-TACC1 cells in triplicate were treated for 6 h with DHT (10nM), CD437 (400 nM) or vehicle, and CUT&RUN undertaken for AR, RARγ or H3K27ac. The number of differentially enriched peaks compared to IgG (n) was determined with `csaw`, and the mean logPV (calculated as -log10(p.adj) for each group) are indicated.

| Cell.ChIP.Rx | ChromHMM | logPV | Threshold |
|---|---|---|---|
| Mock.K27.DHT | Promoter | 317.17 | Significant |
| Mock.K27.DHT | Transcribed | 317.17 | Significant |
| Mock.K27.veh | Promoter | 317.17 | Significant |
| Mock.K27.veh | Transcribed | 317.17 | Significant |
| RARg.AR.DHT | Promoter | 317.17 | Significant |
| RARg.AR.veh | Promoter | 317.17 | Significant |
| RARg.K27.veh | Promoter | 317.17 | Significant |
| RARg.K27.DHT | Promoter | 317.17 | Significant |
| RARg.K27.CD437 | Promoter | 317.17 | Significant |
| RARg.K27.CD437 | Transcribed | 317.17 | Significant |
| TACC1.RARg.AR.DHT | Transcribed | 317.17 | Significant |
| TACC1.RARg.AR.DHT | Poised_Enhancer | 317.17 | Significant |
| TACC1.RARg.AR.DHT | Promoter | 317.17 | Significant |
| TACC1.RARg.AR.DHT | Active_Enhancer | 317.17 | Significant |
| TACC1.RARg.AR.DHT | Polycomb | 317.17 | Significant |
| TACC1.RARg.K27.CD437 | Promoter | 317.17 | Significant |
| TACC1.RARg.K27.CD437 | Transcribed | 317.17 | Significant |
| TACC1.RARg.K27.veh | Promoter | 317.17 | Significant |
| TACC1.RARg.K27.veh | Active_Enhancer | 317.17 | Significant |
| TACC1.RARg.K27.veh | Transcribed | 317.17 | Significant |
| TACC1.RARg.K27.DHT | Promoter | 317.17 | Significant |
| TACC1.RARg.K27.DHT | Transcribed | 317.17 | Significant |
| TACC1.RARg.K27.DHT | Active_Enhancer | 317.17 | Significant |
| TACC1.RARg.RARg.veh | Promoter | 317.17 | Significant |
| TACC1.RARg.RARg.veh | Poised_Enhancer | 317.17 | Significant |
| TACC1.RARg.RARg.veh | Active_Enhancer | 317.17 | Significant |
| TACC1.RARg.RARg.veh | Transcribed | 317.17 | Significant |
| TACC1.RARg.K27.CD437 | Active_Enhancer | 316.88 | Significant |
| RARg.RARg.veh | Promoter | 297.22 | Significant |
| TACC1.RARg.RARg.CD437 | Promoter | 244.78 | Significant |
| RARg.K27.veh | Transcribed | 225.56 | Significant |
| RARg.RARg.CD437 | Promoter | 207.81 | Significant |
| TACC1.RARg.RARg.veh | Polycomb | 143.32 | Significant |
| RARg.K27.DHT | Active_Enhancer | 116.49 | Significant |
| RARg.K27.DHT | Transcribed | 99.32 | Significant |
| RARg.AR.DHT | Active_Enhancer | 98.98 | Significant |
| RARg.K27.CD437 | Active_Enhancer | 98.22 | Significant |
| TACC1.RARg.AR.DHT | Bivalent_Promoter | 93.86 | Significant |
| Mock.AR.veh | Promoter | 91.28 | Significant |
| Mock.AR.DHT | Promoter | 86.14 | Significant |
| RARg.K27.veh | Active_Enhancer | 80.35 | Significant |
| Mock.K27.veh | Active_Enhancer | 60.28 | Significant |
| RARg.AR.veh | Active_Enhancer | 55.61 | Significant |
| Mock.AR.DHT | Active_Enhancer | 49.43 | Significant |
| Mock.K27.DHT | Active_Enhancer | 46.12 | Significant |
| TACC1.RARg.RARg.veh | Bivalent_Promoter | 5.63 | Significant |
| Mock.AR.veh | Active_Enhancer | 2.37 | Significant |

**Supplementary Table 4B**: Significant enrichment of AR, RARγ and H3K27ac cistromes in ChromHMM defined epigenetic states. 22Rv1-mock, 22Rv1-RARγ and 22Rv1-RARγ-TACC1 cells in triplicate were treated for 6h with vehicle DHT (10nM), or CD437 (400 nM), and CUT&RUN undertaken for AR, RARγ or H3K27ac. Differential enriched binding sites compared to IgG controls (p.adj < .1) were identified by `csaw` and were overlapped using `bedtools` with ChromHMM regions (ChromHMM) identified in LNCaP, and enrichment tested with a hypergeometric test (lower.tail = FALSE).

| Cell.Rx | class | coreg.freq | Nearest.Example |
|---|---|---|---|
| Mock_DHT | other | 470 | LPAR3 |
| Mock_DHT | CoR | 13 | SNX6 |
| Mock_DHT | TF | 12 | CASZ1 |
| Mock_DHT | Mixed | 10 | ZBTB16 |
| Mock_DHT | CoA | 6 | ZBTB7B |
| Mock_RARg_DHT | other | 45 | LPAR3 |
| Mock_RARg_DHT | TF | 2 | NR4A1 |
| Mock_RARg_DHT | CoR | 1 | SNX6 |
| Mock_RARg_M | other | 30 | MYBPC1 |
| Mock_RARg_M | TF | 2 | NR4A1 |
| Mock_RARg_TACC1.RARg_DHT | other | 52 | RP11-122G18.12 |
| Mock_RARg_TACC1.RARg_DHT | TF | 1 | THRB |
| Mock_RARg_TACC1.RARg_M | other | 12 | REXO1L7P |
| Mock_TACC1.RARg_DHT | other | 22 | KLK15 |
| Mock_TACC1.RARg_DHT | Mixed | 1 | ZBTB16 |
| Mock_TACC1.RARg_M | other | 71 | RP11-122G18.12 |
| Mock_TACC1.RARg_M | TF | 3 | THRB |
| Mock_TACC1.RARg_M | Mixed | 1 | ZBTB16 |
| RARg_DHT | other | 48 | RP11-554D13.1 |
| RARg_DHT | Mixed | 3 | HFE2 |
| RARg_DHT | CoR | 2 | CCND1 |
| RARg_DHT | CoA | 1 | BMPR1B |
| RARg_DHT | TF | 1 | FOXA1 |
| RARg_M | other | 46 | RP11-554D13.1 |
| RARg_M | CoR | 2 | TXNIP |
| RARg_M | Mixed | 2 | HFE2 |
| RARg_M | TF | 2 | HIST2H2BE |
| RARg_M | CoA | 2 | SF3B4 |
| TACC1.RARg_DHT | other | 53 | DBET |
| TACC1.RARg_DHT | TF | 20 | DUX4L8 |
| TACC1.RARg_M | other | 123 | REXO1L9P |
| TACC1.RARg_M | TF | 8 | FOXA1 |
| TACC1.RARg_M | CoR | 5 | CCND1 |
| TACC1.RARg_M | Mixed | 3 | NFIX |
| TACC1.RARg_M | CoA | 1 | KDM6B |
| TACC1.RARg_RARg_DHT | other | 21 | REXO1L8P |
| TACC1.RARg_RARg_DHT | TF | 1 | PPARA |
| TACC1.RARg_RARg_M | other | 36 | REXO1L8P |
| TACC1.RARg_RARg_M | TF | 5 | ZNF565 |

**Supplementary Table 4C**: Coregulator and transcription factor annotation of genes associated with super enhancers. 22Rv1, 22Rv1-RAR$\gamma$ and 22Rv1-RAR$\gamma$-TACC1 cells in triplicate were treated with vehicle or DHT (10nM, 6h) or vehicle, and CUT&RUN undertaken for H3K27ac, and differentially-enriched sites were analyzed, including those at the sites of shared binding, with the `ROSE` algorithm and using BRD4 as a enhancer-enriched factor to generate high confidence super enhancer sites (SE). The SE were annotated to genes within 100 kb and classified to identify coregulators (CoA, CoR, Mixed) and TFs, or other. The closest example is shown in each cell, treatment and coregulator class.

| Bkgd | ChIP | Rx | n | logPval |
|---|---|---|---|---|
| Syn | AR | DHT | 47695 | 1.24 |
| Syn | AR | veh | 30 | 1.66 |
| Asy | AR | DHT | 15545 | 1.54 |
| Asy | AR | veh | 6354 | 1.26 |
| Syn | RARγ | DHT | 2035 | 1.27 |
| Syn | RARγ | veh | 2037 | 1.42 |
| Syn | H3S10 | DHT | 96123 | 3.60 |
| Syn | H3S10 | veh | 151480 | 2.25 |
| Asy | H3S10 | veh | 1634 | 2.13 |
| Asy | RARγ | DHT | 388 | 1.41 |
| Asy | RARγ | veh | 295 | 1.41 |

**Supplementary Table 5**: Cistrome in G2/M synchronized and asynchronized cells. 22Rv1-RARγ cells in triplicate were treated with vehicle or nocodozole (*60 ng*, 18h) and then (DHT (10nM, 6 h) or vehicle, and CUT&RUN undertaken for AR, RARγ and H3S10P, and differentially-enriched sites identified with `csaw`. The number of differentially enriched peaks compared to IgG (n), and the mean logPV (calculated as -log10(p.adj) for each group are indicated.

| Gen.Rx | D | MostSig | Lum | Bas | CoA | CoR | Mix | TF | NR | in.miR | MRE |
|---|---|---|---|---|---|---|---|---|---|---|---|
| Mock_CD437 | N | YAP1 | 0 | 0 | 0 | 0 | 62 | 0 | 0 | 0 | 0 |
| Mock_CD437 | N | PEG10 | 0 | 128 | 0 | 0 | 0 | 0 | 0 | 0 | 0 |
| Mock_CD437 | P | DHRS3 | 0 | 0 | 0 | 0 | 0 | 0 | 0 | 126 | 107 |
| Mock_CD437 | P | EPAS1 | 0 | 0 | 0 | 0 | 0 | 159 | 0 | 0 | 0 |
| Mock_CD437 | P | NR6A1 | 0 | 0 | 0 | 0 | 0 | 159 | 9 | 0 | 0 |
| Mock_CD437 | P | BTG2 | 0 | 0 | 0 | 47 | 0 | 0 | 0 | 0 | 0 |
| Mock_CD437 | P | BAMBI | 188 | 0 | 73 | 0 | 0 | 0 | 0 | 0 | 0 |
| Mock_DHT | P | SHROOM3 | 260 | 0 | 0 | 0 | 0 | 0 | 0 | 153 | 124 |
| Mock_DHT | P | CEBPD | 0 | 0 | 0 | 0 | 0 | 206 | 0 | 0 | 0 |
| Mock_DHT | P | NT5DC3 | 0 | 155 | 0 | 0 | 0 | 0 | 0 | 0 | 0 |
| Mock_DHT | P | YAP1 | 0 | 0 | 0 | 0 | 61 | 0 | 0 | 0 | 0 |
| Mock_DHT | P | CD36 | 0 | 0 | 0 | 60 | 0 | 0 | 0 | 0 | 0 |
| Mock_DHT | P | FZD5 | 0 | 0 | 88 | 0 | 0 | 0 | 0 | 0 | 0 |
| Mock_DHT | P | RARB | 0 | 0 | 0 | 0 | 0 | 0 | 10 | 0 | 0 |
| Mock_ENZA | N | BTG1 | 0 | 0 | 0 | 0 | 0 | 141 | 0 | 0 | 0 |
| Mock_ENZA | N | TMEFF2 | 163 | 0 | 0 | 0 | 0 | 0 | 0 | 0 | 0 |
| Mock_ENZA | N | STEAP2 | 0 | 0 | 0 | 0 | 0 | 0 | 0 | 0 | 89 |
| Mock_ENZA | N | BARD1 | 0 | 0 | 0 | 0 | 50 | 0 | 0 | 0 | 0 |
| Mock_ENZA | N | NR3C2 | 0 | 0 | 0 | 0 | 0 | 0 | 13 | 0 | 0 |
| Mock_ENZA | P | GALNT10 | 0 | 0 | 0 | 0 | 0 | 0 | 0 | 106 | 0 |
| Mock_ENZA | P | PRKD1 | 0 | 0 | 60 | 0 | 0 | 0 | 0 | 0 | 0 |
| Mock_ENZA | P | TOB1 | 0 | 0 | 0 | 29 | 0 | 0 | 0 | 0 | 0 |
| Mock_ENZA | P | SPRY2 | 0 | 92 | 0 | 0 | 0 | 0 | 0 | 0 | 0 |
| RARg_CD437 | N | KLHL29 | 0 | 128 | 0 | 0 | 0 | 0 | 0 | 0 | 0 |
| RARg_CD437 | P | SHROOM1 | 184 | 0 | 0 | 0 | 0 | 0 | 0 | 0 | 0 |
| RARg_CD437 | P | DHRS3 | 0 | 0 | 0 | 0 | 0 | 0 | 0 | 124 | 112 |
| RARg_CD437 | P | EPAS1 | 0 | 0 | 0 | 0 | 0 | 199 | 0 | 0 | 0 |
| RARg_CD437 | P | BAMBI | 0 | 0 | 77 | 0 | 0 | 0 | 0 | 0 | 0 |
| RARg_CD437 | P | NRIP1 | 0 | 0 | 0 | 0 | 64 | 0 | 0 | 0 | 0 |
| RARg_CD437 | P | BTG2 | 0 | 0 | 0 | 44 | 0 | 0 | 0 | 0 | 0 |
| RARg_CD437 | P | NR6A1 | 0 | 0 | 0 | 0 | 0 | 0 | 20 | 0 | 0 |
| RARg_DHT | P | ELOVL2 | 186 | 0 | 0 | 0 | 0 | 0 | 0 | 0 | 0 |
| RARg_DHT | P | NT5DC3 | 0 | 97 | 0 | 0 | 0 | 0 | 0 | 0 | 0 |
| RARg_DHT | P | KLF9 | 0 | 0 | 0 | 0 | 0 | 120 | 0 | 0 | 0 |
| RARg_DHT | P | TMEM164 | 0 | 0 | 0 | 0 | 0 | 0 | 0 | 100 | 0 |
| RARg_DHT | P | SHROOM3 | 0 | 0 | 0 | 0 | 0 | 0 | 0 | 0 | 92 |
| RARg_DHT | P | FZD5 | 0 | 0 | 49 | 0 | 0 | 0 | 0 | 0 | 0 |
| RARg_DHT | P | FZD1 | 0 | 0 | 0 | 0 | 33 | 0 | 0 | 0 | 0 |
| RARg_DHT | P | TFCP2L1 | 0 | 0 | 0 | 41 | 0 | 0 | 0 | 0 | 0 |
| RARg_DHT | P | RARB | 0 | 0 | 0 | 0 | 0 | 0 | 3 | 0 | 0 |
| RARg_ENZA | N | DKK1 | 0 | 62 | 0 | 26 | 0 | 0 | 0 | 0 | 0 |
| RARg_ENZA | P | DHRS3 | 0 | 0 | 0 | 0 | 0 | 0 | 0 | 82 | 63 |
| RARg_ENZA | P | ATP1B1 | 123 | 0 | 0 | 0 | 0 | 0 | 0 | 0 | 0 |
| RARg_ENZA | P | BAMBI | 123 | 0 | 34 | 0 | 0 | 0 | 0 | 0 | 0 |
| RARg_ENZA | P | EPAS1 | 0 | 0 | 0 | 0 | 0 | 111 | 0 | 0 | 0 |
| RARg_ENZA | P | NRIP1 | 0 | 0 | 0 | 0 | 28 | 0 | 0 | 0 | 0 |
| RARg_ENZA | P | PPARA | 0 | 0 | 0 | 0 | 0 | 0 | 10 | 0 | 0 |
| RARg_veh | N | PLEKHG5 | 0 | 100 | 0 | 0 | 0 | 0 | 0 | 0 | 0 |
| RARg_veh | N | AGO1 | 0 | 0 | 44 | 0 | 0 | 0 | 0 | 0 | 0 |
| RARg_veh | N | HNF4A | 0 | 0 | 0 | 0 | 0 | 0 | 8 | 0 | 0 |
| RARg_veh | P | CLU | 0 | 0 | 0 | 0 | 0 | 0 | 0 | 79 | 0 |
| RARg_veh | P | SOX9 | 0 | 0 | 0 | 0 | 0 | 119 | 0 | 0 | 71 |

| | | | | | | | | | | |
|---|---|---|---|---|---|---|---|---|---|---|
| RARg_veh | P | SLC12A2 | 139 | 0 | 0 | 0 | 0 | 0 | 0 | 0 | 0 |
| RARg_veh | P | LEF1 | 0 | 0 | 0 | 0 | 33 | 0 | 0 | 0 | 0 |
| RARg_veh | P | PPID | 0 | 0 | 0 | 25 | 0 | 0 | 0 | 0 | 0 |
| TACC1.RARg_CD437 | P | SHROOM1 | 261 | 0 | 0 | 0 | 0 | 0 | 0 | 0 | 0 |
| TACC1.RARg_CD437 | P | ABLIM3 | 0 | 0 | 136 | 0 | 0 | 0 | 0 | 0 | 0 |
| TACC1.RARg_CD437 | P | NRIP1 | 0 | 0 | 0 | 0 | 95 | 0 | 0 | 0 | 0 |
| TACC1.RARg_CD437 | P | CORO2A | 0 | 0 | 0 | 86 | 0 | 0 | 0 | 0 | 0 |
| TACC1.RARg_CD437 | P | PPARD | 0 | 0 | 0 | 0 | 0 | 305 | 19 | 0 | 0 |
| TACC1.RARg_CD437 | P | EPAS1 | 0 | 0 | 0 | 0 | 0 | 0 | 0 | 0 | 178 |
| TACC1.RARg_CD437 | P | UPP1 | 0 | 191 | 0 | 0 | 0 | 0 | 0 | 0 | 0 |
| TACC1.RARg_CD437 | P | DHRS3 | 0 | 0 | 0 | 0 | 0 | 0 | 0 | 226 | 0 |
| TACC1.RARg_DHT | P | ELOVL2 | 110 | 0 | 0 | 0 | 0 | 0 | 0 | 0 | 0 |
| TACC1.RARg_DHT | P | KCNJ11 | 110 | 0 | 0 | 0 | 0 | 0 | 0 | 0 | 0 |
| TACC1.RARg_DHT | P | TFEB | 0 | 0 | 0 | 0 | 0 | 59 | 0 | 0 | 0 |
| TACC1.RARg_DHT | P | NT5DC3 | 0 | 40 | 0 | 0 | 0 | 0 | 0 | 0 | 0 |
| TACC1.RARg_DHT | P | FSTL1 | 0 | 0 | 0 | 0 | 0 | 0 | 0 | 61 | 0 |
| TACC1.RARg_DHT | P | SHROOM3 | 0 | 0 | 0 | 0 | 0 | 0 | 0 | 0 | 52 |
| TACC1.RARg_DHT | P | YAP1 | 0 | 0 | 0 | 0 | 21 | 0 | 0 | 0 | 0 |
| TACC1.RARg_DHT | P | FZD5 | 0 | 0 | 26 | 0 | 0 | 0 | 0 | 0 | 0 |
| TACC1.RARg_DHT | P | RARB | 0 | 0 | 0 | 0 | 0 | 0 | 3 | 0 | 0 |
| TACC1.RARg_DHT | P | PIAS1 | 0 | 0 | 0 | 20 | 0 | 0 | 0 | 0 | 0 |
| TACC1.RARg_ENZA | N | DKK1 | 0 | 16 | 0 | 0 | 0 | 0 | 0 | 0 | 0 |
| TACC1.RARg_ENZA | P | CORO2A | 0 | 0 | 0 | 11 | 0 | 0 | 0 | 0 | 0 |
| TACC1.RARg_ENZA | P | SHROOM1 | 72 | 0 | 0 | 0 | 0 | 0 | 0 | 0 | 0 |
| TACC1.RARg_ENZA | P | ABLIM3 | 0 | 0 | 15 | 0 | 0 | 0 | 0 | 0 | 0 |
| TACC1.RARg_ENZA | P | NRIP1 | 0 | 0 | 0 | 0 | 12 | 0 | 0 | 0 | 0 |
| TACC1.RARg_ENZA | P | DHRS3 | 0 | 0 | 0 | 0 | 0 | 0 | 0 | 29 | 37 |
| TACC1.RARg_ENZA | P | PPARD | 0 | 0 | 0 | 0 | 0 | 39 | 6 | 0 | 0 |
| TACC1.RARg_veh | N | KNTC1 | 0 | 0 | 0 | 0 | 0 | 352 | 0 | 0 | 0 |
| TACC1.RARg_veh | N | SMC4 | 0 | 0 | 0 | 0 | 0 | 0 | 0 | 224 | 0 |
| TACC1.RARg_veh | N | ZNF367 | 0 | 0 | 0 | 0 | 0 | 0 | 0 | 0 | 217 |
| TACC1.RARg_veh | N | TMEM237 | 0 | 194 | 0 | 0 | 0 | 0 | 0 | 0 | 0 |
| TACC1.RARg_veh | N | CDK2 | 0 | 0 | 143 | 0 | 0 | 0 | 0 | 0 | 0 |
| TACC1.RARg_veh | N | TIMELESS | 0 | 0 | 0 | 108 | 0 | 0 | 0 | 0 | 0 |
| TACC1.RARg_veh | N | BRIP1 | 0 | 0 | 0 | 0 | 99 | 0 | 0 | 0 | 0 |
| TACC1.RARg_veh | P | SUSD2 | 295 | 0 | 0 | 0 | 0 | 0 | 0 | 0 | 0 |
| TACC1.RARg_veh | P | RARG | 0 | 0 | 0 | 0 | 0 | 0 | 16 | 0 | 0 |

**Supplementary Table 6A** – **Categorization of DEGs.** LNCaP, 22Rv1-Mock, 22Rv1-RAR$\gamma$ and 22Rv1-RAR$\gamma$-TACC1 cells in triplicate were treated with vehicle or DHT (10nM), ENZA (10mM), or CD437 (400nM) for 24 h and RNA-Seq undertaken. Each DEG list was annotated as to whether the gene was contained within a luminal (Lum) or basal prostate gene signature (Bas), a coactivator (CoA), corepressor (CoR), mixed function coregulator (mixed), a transcription factor (TF), a nuclear receptor (NR), an intron-encoded miRNA (in.miR) or contained a miR-96 binding site identified by IMPACT-Seq and was differentially expressed either at the mRNA or protein level (MRE). The number of genes in each category, in each cell background and treatment is indicated, and the most significant gene in each class given.

| Gen.Rx | Overlap | logPV | Threshold |
|---|---|---|---|
| RARg_CD437 | 20 | 4.85 | Significant |
| TACC1.RARg_ENZA | 6 | 2.59 | Significant |
| Mock_ENZA | 13 | 2.24 | Significant |
| RARg_ENZA | 10 | 1.80 | Significant |
| TACC1.RARg_CD437 | 19 | 1.68 | Significant |
| RARg_veh | 8 | 1.03 | Significant |
| Mock_DHT | 10 | 0.43 | NS |
| Mock_CD437 | 9 | 0.39 | NS |
| TACC1.RARg_DHT | 3 | 0.24 | NS |
| TACC1.RARg_veh | 16 | 0.19 | NS |
| RARg_DHT | 3 | 0.04 | NS |

**Supplementary Table 6B – Enrichment of NRs in DEGs.** LNCaP, 22Rv1-Mock, 22Rv1-RARγ and 22Rv1-RARγ-TACC1 cells in triplicate were treated with vehicle or DHT (10nM), ENZA (10mM), or CD437 (400nM) for 24 h and RNA-Seq undertaken. Each DEG list was annotated to identify nuclear receptors (NR). A hypergeometric test was undertaken to examine if the number of NR were significantly enriched in each DEGs list.

| Gen.Rx | Overlap | logPV | Threshold |
|---|---|---|---|
| TACC1.RARg_ENZA | 37 | 3.94 | Significant |
| RARg_DHT | 92 | 2.11 | Significant |
| TACC1.RARg_DHT | 52 | 1.39 | Significant |
| TACC1.RARg_CD437 | 178 | 0.72 | NS |
| RARg_CD437 | 112 | 0.71 | NS |
| RARg_veh | 71 | 0.64 | NS |
| Mock_ENZA | 89 | 0.51 | NS |
| Mock_DHT | 124 | 0.44 | NS |
| RARg_ENZA | 63 | 0.22 | NS |
| Mock_CD437 | 107 | 0.18 | NS |
| TACC1.RARg_veh | 217 | 0.04 | NS |

**Supplementary Table 6C – Enrichment of miR96 bound and regulated genes in DEGs.** LNCaP, 22Rv1, 22Rv1-RARγ and 22Rv1-RARγ-TACC1 cells in triplicate were treated with vehicle or DHT (10nM), ENZA (10mM), or CD437 (400nM) for 24 h and RNA-Seq undertaken. Each DEG list was annotated to identify miR96 bound and regulated genes. A hypergeometric test was undertaken to examine if the number of miR96 bound and regulated genes were significantly enriched in each DEGs list.

| Cell.Rx | class | NumberGenes | MostSignificant |
|---|---|---|---|
| Mock_DHT | other | 127 | CNTROB |
| Mock_DHT | TF | 13 | TFEB |
| Mock_DHT | miR96.target | 7 | SEMA3C |
| Mock_DHT | Mixed | 5 | TANC2 |
| Mock_DHT | intragenic | 5 | WIZ |
| Mock_DHT | CoA | 4 | SMARCE1 |
| Mock_DHT | CoR | 2 | SIRT7 |
| Mock_DHT | NR | 1 | NR1H3 |
| Mock_ENZA | other | 207 | CNTROB |
| Mock_ENZA | TF | 20 | AR |
| Mock_ENZA | CoA | 17 | SMARCE1 |
| Mock_ENZA | intragenic | 13 | AATK |
| Mock_ENZA | Mixed | 7 | UHRF2 |
| Mock_ENZA | miR96.target | 6 | PLA2G6 |
| Mock_ENZA | CoR | 5 | PCGF2 |
| Mock_ENZA | NR | 2 | AR |
| Mock_CD437 | other | 105 | PI4KAP2 |
| Mock_CD437 | CoA | 8 | MED12L |
| Mock_CD437 | CoR | 6 | NCOR2 |
| Mock_CD437 | intragenic | 6 | NCOR2 |
| Mock_CD437 | miR96.target | 4 | JPT2 |
| Mock_CD437 | TF | 2 | RERE |
| Mock_CD437 | Mixed | 1 | KRIT1 |
| RARg_DHT | other | 109 | DLD |
| RARg_DHT | TF | 13 | SOX13 |
| RARg_DHT | CoR | 6 | GFI1 |
| RARg_DHT | intragenic | 6 | MAD1L1 |
| RARg_DHT | miR96.target | 4 | SEMA3C |
| RARg_DHT | Mixed | 4 | ASXL1 |
| RARg_DHT | CoA | 2 | KAT5 |
| RARg_ENZA | other | 131 | SLC25A36 |
| RARg_ENZA | TF | 11 | AR |
| RARg_ENZA | intragenic | 9 | MAD1L1 |
| RARg_ENZA | CoA | 6 | HYAL2 |
| RARg_ENZA | miR96.target | 6 | ARL1 |
| RARg_ENZA | CoR | 5 | NCOR2 |
| RARg_ENZA | Mixed | 3 | TBL1XR1 |
| RARg_ENZA | NR | 1 | AR |
| RARg_CD437 | other | 477 | TBL1XR1 |
| RARg_CD437 | CoA | 40 | SMARCA4 |
| RARg_CD437 | TF | 37 | ZNF544 |
| RARg_CD437 | intragenic | 36 | TLE3 |
| RARg_CD437 | miR96.target | 19 | ARPC1B |
| RARg_CD437 | CoR | 16 | TLE3 |
| RARg_CD437 | Mixed | 12 | TBL1XR1 |
| RARg_CD437 | NR | 3 | NR1H3 |
| RARg.Mock_DHT | other | 157 | SPAG5 |
| RARg.Mock_DHT | TF | 16 | AR |
| RARg.Mock_DHT | intragenic | 10 | UHRF1 |
| RARg.Mock_DHT | Mixed | 8 | CHD8 |
| RARg.Mock_DHT | miR96.target | 7 | UHRF1 |
| RARg.Mock_DHT | CoR | 5 | UHRF1 |

| | | | |
|---|---|---|---|
| RARg.Mock_DHT | CoA | 3 | SMARCD3 |
| RARg.Mock_DHT | NR | 2 | AR |
| RARg.Mock_ENZA | other | 153 | SPAG5 |
| RARg.Mock_ENZA | TF | 14 | RARG |
| RARg.Mock_ENZA | intragenic | 9 | ABR |
| RARg.Mock_ENZA | miR96.target | 8 | RARG |
| RARg.Mock_ENZA | CoA | 6 | PRMT2 |
| RARg.Mock_ENZA | Mixed | 4 | UHRF2 |
| RARg.Mock_ENZA | NR | 2 | RARG |
| RARg.Mock_ENZA | CoR | 2 | SNX6 |
| RARg.Mock_CD437 | other | 199 | SPAG5 |
| RARg.Mock_CD437 | TF | 23 | AR |
| RARg.Mock_CD437 | intragenic | 15 | ZBTB20 |
| RARg.Mock_CD437 | CoA | 10 | RRN3 |
| RARg.Mock_CD437 | CoR | 9 | ZBTB20 |
| RARg.Mock_CD437 | Mixed | 7 | TBL1XR1 |
| RARg.Mock_CD437 | miR96.target | 5 | RARG |
| RARg.Mock_CD437 | NR | 3 | AR |

**Supplementary Table 6D – Categorization of DETs in 22Rv1 variants.** 22Rv1, 22Rv1-RAR$\gamma$ and 22Rv1-RAR$\gamma$-TACC1 cells in triplicate were treated with DHT (10nM), ENZA (10mM), or CD437 (400nM) for 24 h and RNA-Seq undertaken. FASTQ files were processed with a transcript-aware alignment tool (`Salmon`) and differentially-enriched transcripts (DETs) were identified. The DET lists was annotated to identify coregulators (CoA, CoR, Mixed) and TFs; miR96 bound and regulated genes (miR96.target); or protein-coding genes that contained miRNA (intragenic). The most significant DET in each cell, treatment and category is indicated.

| Cell.Rx | class | NumberGenes | MostSignificant |
|---|---|---|---|
| ENZA_miR96 | other | 82 | PROM2 |
| ENZA_miR96 | intragenic | 9 | EVL |
| ENZA_miR96 | CoA | 7 | NCK1 |
| ENZA_miR96 | TF | 6 | TGIF1 |
| ENZA_miR96 | Mixed | 5 | HNRNPA3 |
| ENZA_miR96 | miR96.target | 2 | DUT |
| ENZA_miR96 | CoR | 1 | BCLAF1 |
| ENZA_veh | other | 10 | ENST00000570727 |
| ENZA_veh | TF | 2 | ZNF136 |
| ENZA_veh | CoA | 1 | U2AF1 |
| veh_miR96 | other | 65 | EIF3M |
| veh_miR96 | CoA | 7 | PPP1R12A |
| veh_miR96 | TF | 7 | TGIF1 |
| veh_miR96 | intragenic | 5 | CPSF6 |
| veh_miR96 | miR96.target | 3 | OAS3 |
| veh_miR96 | CoR | 2 | ELP2 |
| veh_miR96 | Mixed | 1 | ING4 |

**Supplementary Table 7A – Categorization of DETs following miR-96 antagomir plus Enza.** 22Rv1 cells in triplicate were treated miR-96 antagomir (50nM) or ENZA (10μM), or the combination or vehicle for 24h and RNA-Seq undertaken. FASTQ files were processed with a transcript-aware alignment tool (`Salmon`) and differentially-enriched transcripts (DETs) were identified. The DET lists was annotated to identify coregulators (CoA, CoR, Mixed) and TFs; miR96 bound and regulated genes (miR96.target); or protein-coding genes that contained miRNA (intragenic). The most significant DET in each cell, treatment and category is indicated.

| Cell.Rx | class | hgnc_symbol | logPV | Threshold |
|---|---|---|---|---|
| ENZA_miR96 | CoA | COPA | 1.33 | Significant |
| ENZA_miR96 | CoA | NCK1 | 1.33 | Significant |
| ENZA_miR96 | CoA | CDC40 | 1.33 | Significant |
| ENZA_miR96 | CoA | TCEA1 | 1.33 | Significant |
| ENZA_miR96 | CoA | BLOC1S2 | 1.33 | Significant |
| ENZA_miR96 | CoA | SMARCE1 | 1.33 | Significant |
| ENZA_miR96 | CoA | SMARCD2 | 1.33 | Significant |
| ENZA_miR96 | CoA | SMARCA4 | 1.33 | Significant |

**Supplementary Table 7B – Enrichment of functional categories in DETs.** 22Rv1 cells in triplicate were treated with vehicle or miR-96 antagomir (50nM) or ENZA (10μM), or the combination for 24h or vehicle, and RNA-Seq undertaken. FASTQ files were processed with a transcript-aware alignment tool (Salmon) and differentially-enriched transcripts (DETs) were identified. The DET lists was annotated to identify coregulators (CoA, CoR, Mixed) and TFs; miR96 bound and regulated genes (miR96.target); or protein-coding genes that contained miRNA (intragenic). A hypergeometric test was undertaken to examine if the number of DETs in each category was significantly enriched, and the results for CoAs, which was significant, are shown.
